# Engineering a Biosynthetic Pathway for the Production of (+)-Brevianamides A and B in *Escherichia coli*

**DOI:** 10.1101/2024.12.10.627567

**Authors:** Casandra P Sandoval Hurtado, Samantha P Kelly, Vikram Shende, Makayla Perez, Brian J Curtis, Sean A Newmister, Kaleb Ott, Filipa Pereira, David H Sherman

## Abstract

The privileged fused-ring system comprising the bicyclo[2.2.2]diazaoctane (BDO) core is prevalent in diketopiperazine (DKP) natural products with potent and diverse biological activities, with some being explored as drug candidates. Typically, only low yields of these compounds can be extracted from native fungal producing strains and the available synthetic routes remain challenging due to their structural complexity. BDO-containing DKPs including (+)-brevianamides A and B are assembled via multi-component biosynthetic pathways incorporating non-ribosomal peptide synthetases, prenyltransferases, flavin monooxygenases, cytochrome P450s and semi-pinacolases. To simplify access to this class of alkaloids, we designed an engineered biosynthetic pathway in *Escherichia coli*, composed of six enzymes sourced from different kingdoms of life. The pathway includes a cyclodipeptide synthase (NascA), a cyclodipeptide oxidase (DmtD2/DmtE2), a prenyltransferase (NotF), a flavin-dependent monooxygenase (BvnB), and kinases (PhoN and IPK). Cultivated in glycerol supplemented with prenol, the engineered *E. coli* strain produces 5.3 mg/L of (-)-dehydrobrevianamide E (**4**), which undergoes a terminal, *ex vivo* lithium hydroxide catalyzed rearrangement reaction to yield (+)-brevianamides A and B with a 46% yield and a 92:8 diastereomeric ratio. Additionally, titers of **4** were increased eight-fold by enhancing NADPH pools in the engineered *E. coli* strain. Our study combines synthetic biology, biocatalysis and synthetic chemistry approaches to provide a five-step engineered biosynthetic pathway for producing complex indole alkaloids in *E. coli*.

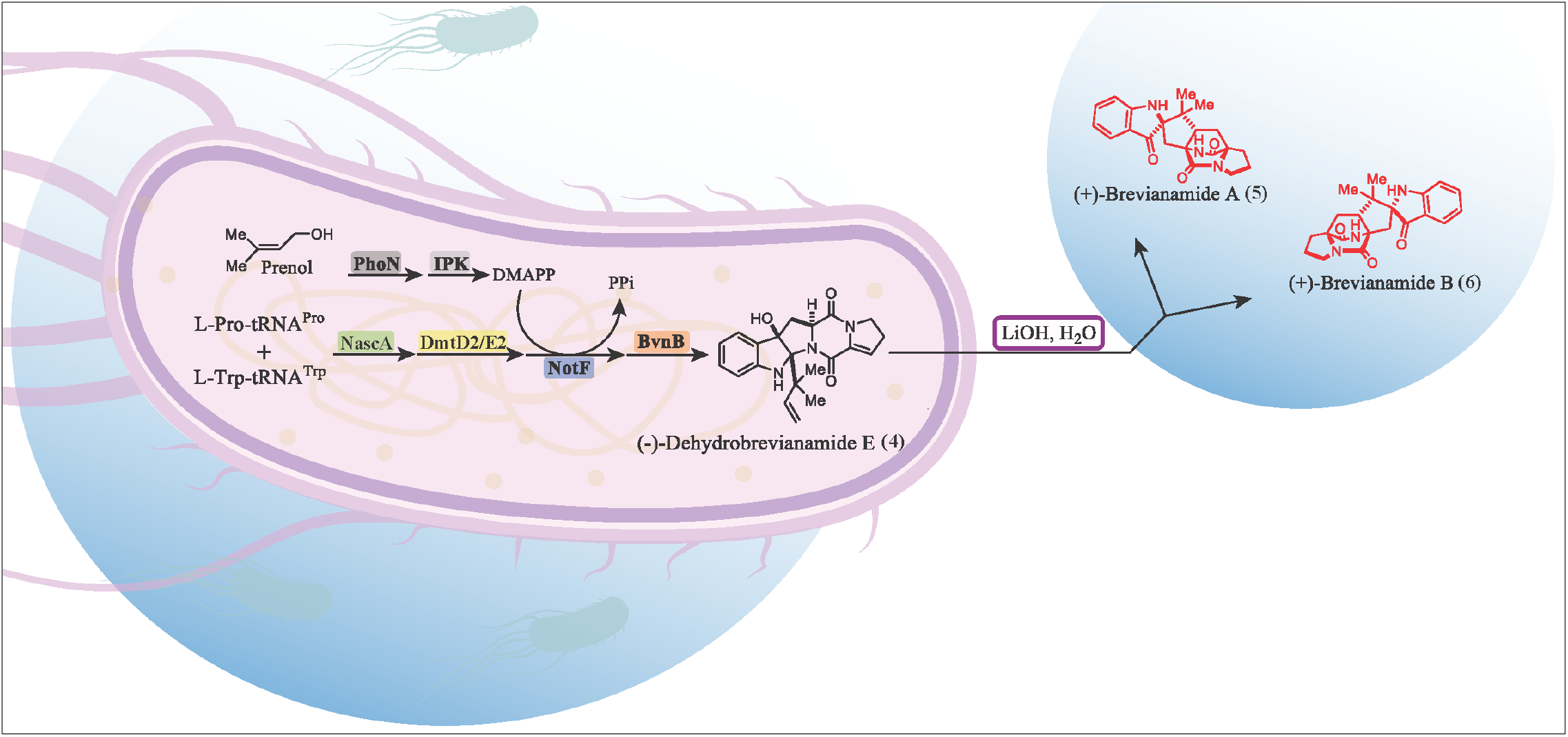

## Introduction

Indole alkaloids with bicyclo[2.2.2]diazaoctane (BDO) cores have long captivated natural product scientists due to their unique structures and biological properties.^1,2^ This group of natural products are subdivided into diketopiperazines (DKPs) or monoketopiperazines (MKPs) in accordance with the number of carbonyl groups on their piperazine scaffold (Scheme 1). BDO products form either *anti-* or *syn*-conformations and are further classified as α or β enantiomers depending on the orientation of the dienophile to the diene.^2^ This group of natural products include the commercial anthelmintic Derquantal^3^, the anti-cancer agent (-)-stephacidin A^4^, and the insecticidal agent (+)-brevianamide A (**5**)^5^, which was the first discovered BDO-containing DKP from *Penicillium brevicompactum*.^2,6,7,8^ The brevianamide DKP BDO core is assembled by a non-enzymatic intramolecular Diels-Alder (IMDA) reaction from the azadiene and prenyl group dienophile moieties.^9,10^

**Scheme 1.**
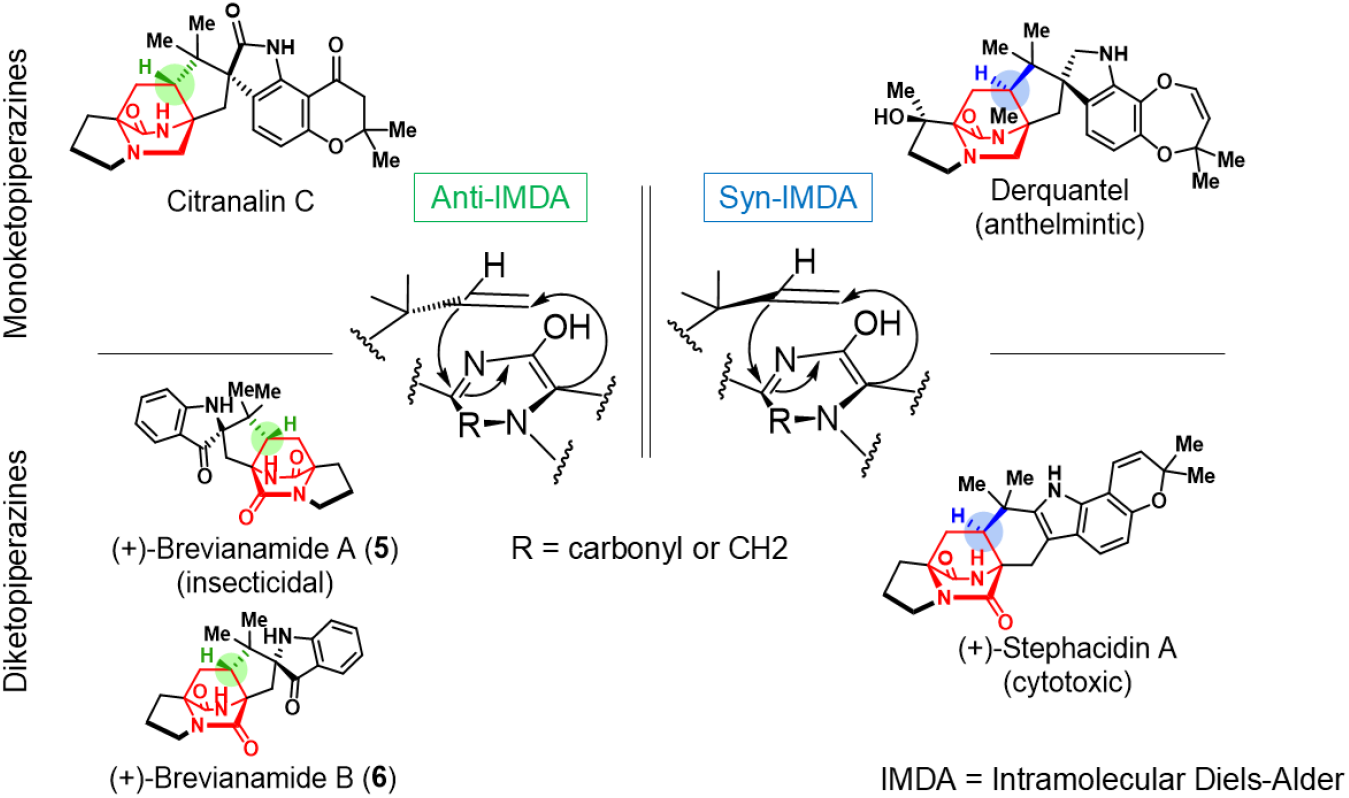
Examples of *anti-* and *syn-*mono and diketopiperazines with bicyclo[2.2.2]diazaoctane cores.

While common in synthetic chemistry, Diels-Alder reactions are relatively rare in nature, with only a handful of Diels-Alderases (DAases) discovered.^11^Notable examples are the recently characterized α-anti selective CtdP and the bifunctional NADPH-dependent reductase/DAase homologs, PhqE and MalC.^12,13^ Of the DAases identified, none have been found in DKP biosynthetic gene clusters (BGCs), including the BGC of (+)-brevianamide A (**5**) and B (**6**) (Figure 1a).^14^ Compounds **5** and **6** contain anti-BDO conformations and arise from a spontaneous IMDA reaction that preferentially forms **5** over **6** in a ratio of 10:1.^15^ Studies on the energetics of the brevianamides have revealed the intrinsic preference of **5** over **6** is due to the favorable intramolecular hydrogen bonds in the transition state of **5**, which are not present in the transition state of **6**.^15,16,17^ In addition, the *syn* isomers of **5** and **6** are not formed due to the geometric constraints imposed by the formation of the 5-membered ring during the IMDA reaction.^9^

**Figure 1.**
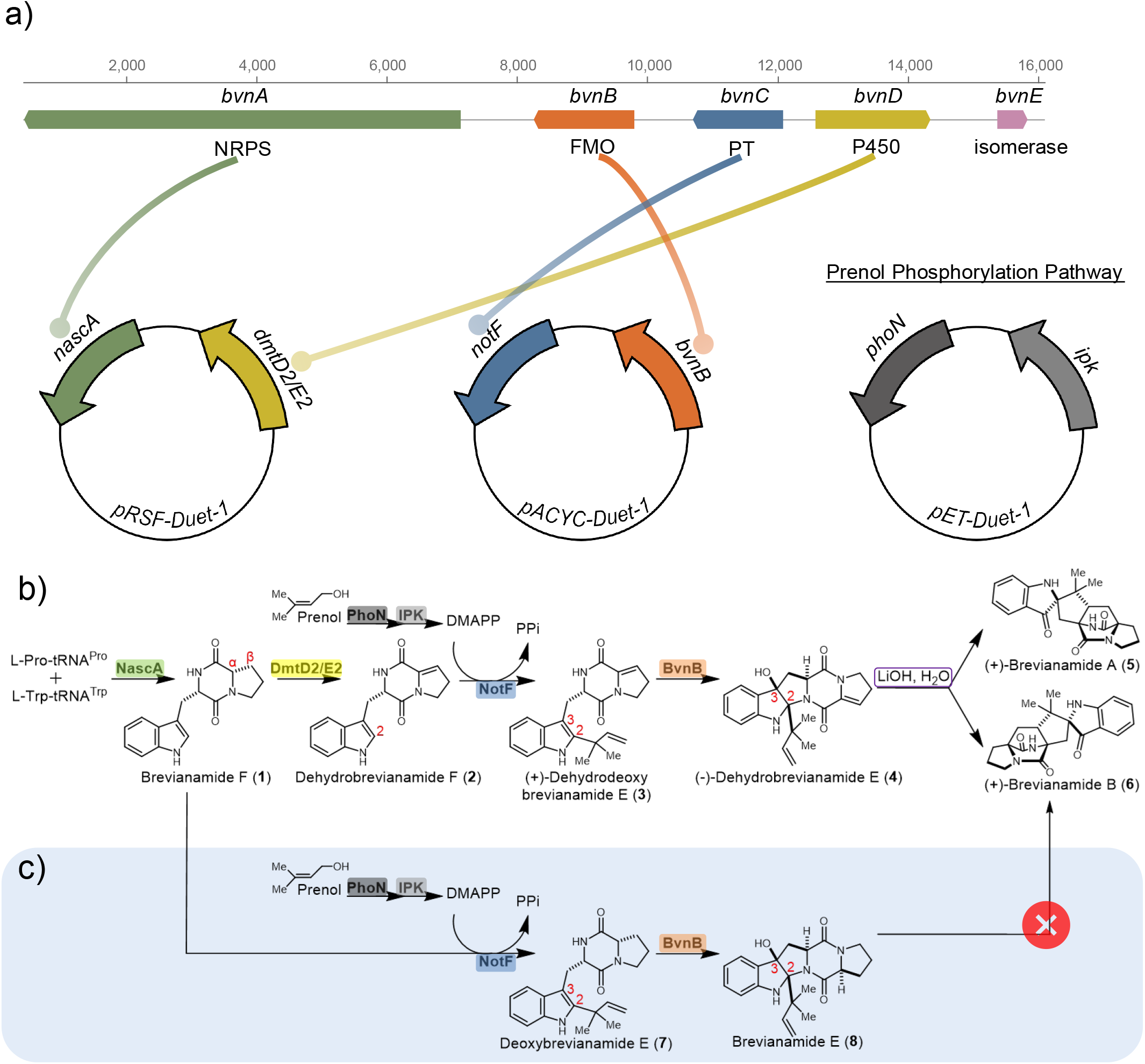
*De novo* bio(synthesis) of diketopiperazines. **a**) Comparison between native brevianamide biosynthetic gene cluster and engineered brevianamide plasmid system. NRPS = non-ribosomal peptide synthetase, FMO = flavin-dependent monooxygenase, PT = prenyltransferase. **b**) Target pathway for the total synthesis of (+)-Brevianamide A (**5**) and (+)-Brevianamide B (**6**). **c**) Shunt pathway resulting from a desaturation on the proline ring.

Although **5** is made in excess amounts over **6** in Nature, the diastereoselective synthesis of **5** has eluded scientists for decades until recently when Lawrence *et al*.^18^ and Bisai *et al*. ^19^ published two approaches for the total synthesis of **5** (Figure S1a-b). In the final two steps of the Lawrence group synthesis, (+)-dehydrodeoxybrevianamide E (**3**) is oxidized to (-)- and (+)-dehydrobrevianamide E (**4**) and (**9**), which are congeners of the native brevianamide pathway shunt intermediate, brevianamide E (**8**) (Figure S2)^18^ The double bond on hydroxypyrroloindolines, **4** and **9**, is key in the base-catalyzed rearrangement cascade wherein an azadiene intermediate is formed followed by a spontaneous [4 + 2] cycloaddition reaction (Figure S3). The Bisai group’s total synthesis begins with compound **10**, a mesylate-containing analog of **3**, and ultimately leads to intermediate **4** after the –OMs elimination step.^19^

Taking inspiration from the synthesis of **5**, we developed an *in vitro* biocatalytic cascade to generate hydroxypyrroloindoline (**8**) and (-)-eurotiumin A by employing enzymes from different DKP biosynthetic pathways, including a prenyltransferase from the notoamide pathway (NotF), and a flavin-dependent monooxygenase (FMO) from the brevianamide pathway (BvnB) (Figure S1c).^20^ Thus, brevianamide F (**1**) was converted into deoxybrevianamide E (**7**) by NotF, followed by a highly stereoselective reaction catalyzed by BvnB to yield **8** as the sole product. This stereoselective approach prevented racemization caused by conventional synthetic oxidizing agents and enabled us to explore product diversification through the substrate flexibility of the biocatalysts. In this investigation, we utilize NotF and BvnB in an expanded biocatalytic cascade genetically embedded in *Escherichia coli* to produce compound **4**. As a final step, we employed a lithium hydroxide catalyzed rearrangement strategy^18^ to produce (+)-brevianamide A (**5**) and B (**6**) (Figure 1b), effectively reconstituting the brevianamide pathway.

The engineered biosynthetic pathway for producing **5** and **6** operates within the engineered *E. coli* strain by overexpressing enzymes from across multiple kingdoms of life and utilizes cost-effective glycerol and prenol as starting materials. As a result, we achieved a 5.3 mg/L titer of IMDA precursor **4**, which was subjected to the LiOH reaction *ex vivo* to achieve a 46% yield of **5** and **6** with a 92:8 diastereomeric ratio. During our studies, we also identified saturated congeners of key intermediates, highlighting the flexibility of downstream engineered biosynthetic pathway enzymes. Furthermore, by upregulating NADPH levels through a CRISPR-Cas9 knock-out of phosphofructokinase A (Δ*pfkA*), we increased the production of (-)-dehydrobrevianamide E (**4**) by eight-fold.

## Results and Discussion

### Overcoming Challenges in the Brevianamide Pathway: NascA and DmtD2/DmtE2 as Surrogate Biocatalysts in DKP Scaffold Formation

In native fungal indole alkaloid biosynthesis, amino acids are condensed to form the DKP scaffold by NRPS enzymes.^15^ NRPSs are large, multi-domain enzymes that are often insoluble in *E. coli*, especially those derived from eukaryotic sources.^12^ Moreover, NRPS thiolation domains require an additional phosphopantetheinyl transferase for post-translational modification. Additional engineering may also be required to deliver sufficient titers of DKPs for downstream tailoring enzymes.^21,22^ Cyclodipeptide synthases (CDPSs) serve as ideal NRPS replacements for heterologous expression in bacteria, as they are bacterial in origin and efficiently produce DKP scaffolds. CDPSs form the DKP ring by catalyzing cyclization of two aminoacyl-tRNAs.^23^ After screening candidate brevianamide F producing CDPS enzymes, we observed heterologous expression of the wild-type CDPS (NascA) from *Streptomyces sp*.CMB-MQ030^24^ efficiently generates brevianamide F (**1**) presumably from endogenous pools of L-Trp-tRNA^Trp^ and L-Pro-tRNA^Pro^ in *E. coli* (Figure S4).

In the native brevianamide pathway, the *bvnD* encoded fungal P450 is presumed to catalyze oxidation followed by spontaneous dehydration and tautomerization of the DKP ring to generate an unstable azadiene, which is central to the formation of the BDO core (Figures 1a, S2).^15^ Fungal P450s, such as BvnD, are typically membrane-bound or membrane-associated,^25^ and their functional expression in bacteria has not been reported.^26^ In addition, they also require cognate reductase partners to be identified and co-expressed. As an alternative approach, we considered the strategy of Lawrence and coworkers, which incorporates desaturation in the DKP core and preserves it until the final tautomerization cascade generates the key azadiene (Figure S3).^18^ One group of enzymes known to catalyze Cα,Cβ-desaturation of DKPs is bacterial cyclodipeptide oxidases (CDOs).^27,28,29^ While dehydrogenating the original DKP scaffold would reorder the biosynthetic steps relative to the native brevianamide pathway, we envisioned using a CDO as a functional surrogate for the native fungal P450 to produce a key pathway precursor, dehydrobrevianamide F (**2**). The genes that form a functional CDO dyad are often found as two overlapping open reading frames, and previous reports have noted separation of these two genes for co-expression often results in insoluble protein.^30^ As such, we obtained a set of synthetic CDO genes with minimal codon optimization to avoid disrupting internal ribosomal binding sites. Whole cell biotransformation reactions were conducted in *E. coli* and screened for production of **2** from brevianamide F (**1**). These assays revealed CDO pair DmtD2/DmtE2 (hereon referred to as DmtD2/E2) from *Streptomyces* sp. F5123^31^ exclusively generated **2** from **1** with no trace of double dehydrogenation (Figures S5). After purifying **2** using semi-preparatory HPLC, we sought to validate NotF and BvnB’s ability to function in our proposed engineered biosynthetic pathway (Figure 1b).

### *In Vitro* Analysis Reveals that NotF and BvnB Can Accept Non-Native Substrates

NotF is a prenyltransferase from the notoamide pathway that utilizes dimethylallyl pyrophosphate (DMAPP) to add a reverse prenyl group at the C-2 position on the DKP indole ring.^32^ In a recent study, we demonstrated that NotF is a versatile enzyme that accepts highly diverse indole-containing DKPs.^20^ These results motivated our use of NotF, rather than the native prenyltransferase of the brevianamide pathway to catalyze the formation of (+)-dehydrodeoxybrevianamide E (**3**). In order to convert **3** into the penultimate substrate **4** for the final LiOH reaction, we chose the BvnB indole oxidase, an FMO that catalyzes initial β-face indole C2-C3 epoxidation followed by N–C bond formation.^15,33^ In the native *P. brevicompactum* pathway, the product of BvnB is immediately oxidized by the BvnD P450 enzyme. However, if the gene coding for the P450 is disrupted, the BvnB product spontaneously cyclizes to form the shunt product, brevianamide E (**8**).^15^ Thus, based on the presumed substrate flexibility of this FMO, we hypothesized BvnB could facilitate the conversion of **3** to the related hydroxypyrroloindoline, (-)- dehydrobrevianamide E (**4**).

The compatibility of NotF and BvnB as components of a one-pot *in vitro* biocatalytic cascade has been validated previously (Figure S1c).^20^ In addition, both enzymes have displayed substrate flexibility by accepting non-native substrates, *cyclo*-L-Trp-L-Ala and preechinulin, *en route* to production of (-)-eurotinium. Since the CDPS and CDO that are used in this study are expected to accept their native substrates from the engineered pathway, NotF and BvnB were tested for their ability to accept non-native substrates, **2** and (+)-dehydrodeoxybrevianamide E (**3**). Thus, an *in vitro* biocatalytic cascade was conducted wherein the DmtD2/E2 product, **2**, was incubated with either NotF and DMAPP or NotF, DMAPP, and BvnB. Both of the desired products, **3** and (-)-dehydrobrevianamide E (**4**), were identified and confirmed using HPLC and LC-MS (Figure 2a Top, S19-20).

**Figure 2.**
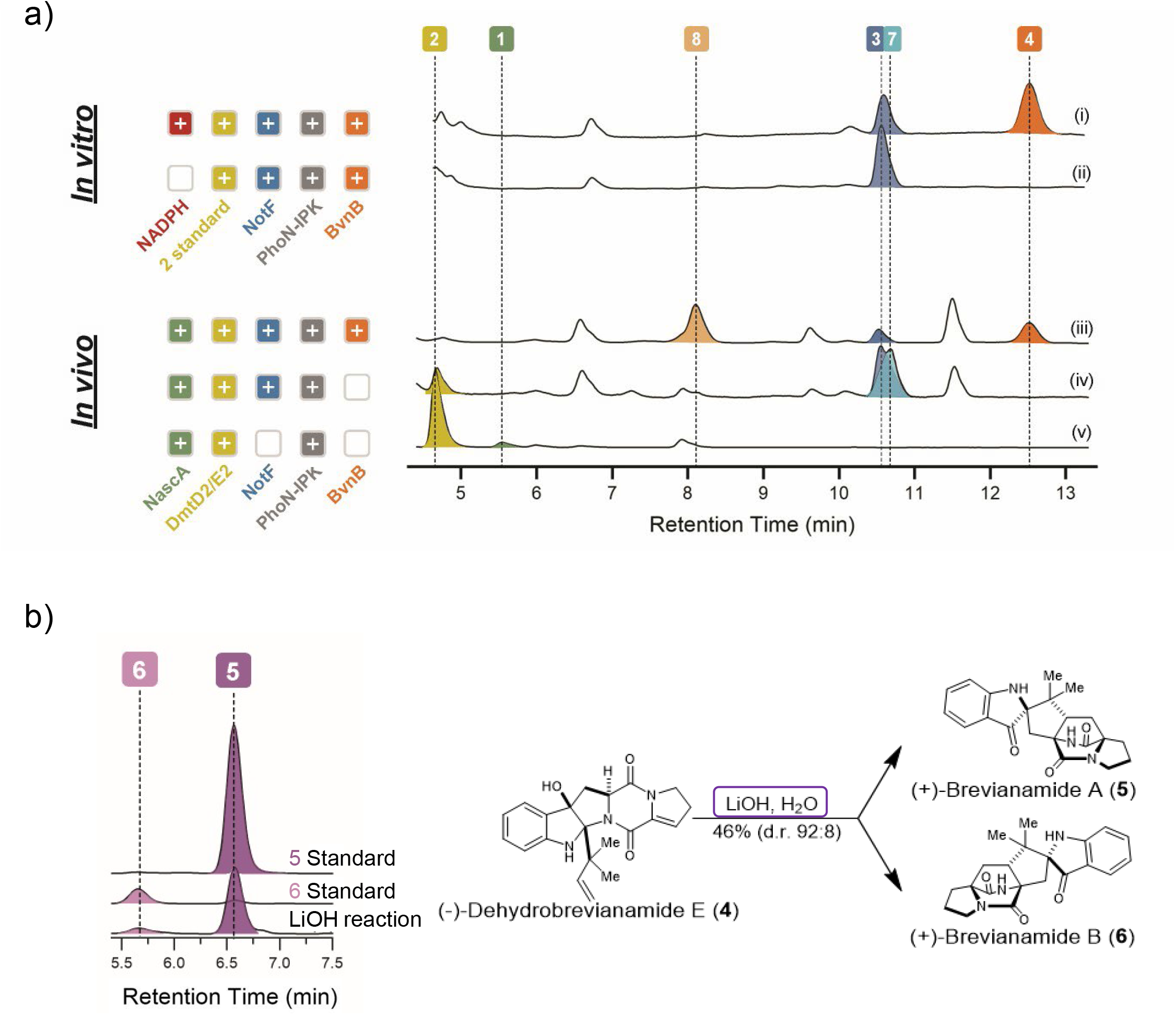
Validation and isolation of all diketopiperazines in target and shunt pathways. **a**) Top: *In vitro* HPLC traces of NotF and BvnB with **2**. (i) **2** + NotF, BvnB, and NADPH (ii) **2** + NotF and BvnB, no NADPH. Bottom: *In vivo* HPLC traces of engineered biosynthetic pathway. (iii) Expression of NascA, DmtD2/E2, PhoN, IPK, NotF, and BvnB (iv) Expression of NascA, DmtD2/E2, PhoN, IPK, and NotF (v) Expression of NascA and DmtD2/E2. All HPLC trace alignments with isolated samples shown in Figures S6, S8. **b**) Lithium hydroxide reaction with **4**. Reaction yields and ratio of **5** and **6**.

### Engineering *E. coli* for *In Vivo* Production of (-)-Dehydrobrevianamide E (4)

Given the efficient *in vivo* production of compound **2** and its compatibility with biosynthetic enzymes NotF and BvnB, we reasoned that expressing NascA, DmtD2/DmtE2, NotF, and BvnB would enable the *in vivo* production of precursor, **4**, which can be subsequently converted into **5** and **6**.

NotF requires DMAPP as a prenyl donor for its reaction. However, *E. coli* has a minimal supply of isoprenoid building blocks, including low, endogenous amounts of DMAPP.^34^ To enable production of this molecule on a large scale, a variety of pathways based on the mevalonate and mevalonate-independent pathways have been developed.^35,36,37^ Recently, a new pathway has been reported wherein only two kinases are necessary to generate biosynthetically useful titers of DMAPP from inexpensive, and non-toxic prenol.^38^ Thus, we tested the two-kinase cassette composed of bacterial PhoN and archaeal IPK reported by Williams and coworkers.^38^ To evaluate if these two kinases can supply a sufficient concentration of DMAPP in the context of our engineered biosynthetic pathway, we fed native substrate **1** to a strain of *E. coli* co-expressing both kinases and NotF. Significant conversion of **1** to deoxybrevianamide E (**7**) was observed (Figure S6). Furthermore, in *E. coli* bioconversion studies wherein **1** is fed to the organism expressing BvnB, NotF, and the kinases, we observed production of **8** (Figure S6) which motivated us to validate the engineered biosynthetic pathway *in vivo*.

Three *pDuet* constructs containing the six genes encoding for the new biosynthetic pathway (*nascA, dmtD2*/*dmtE2, phoN, ipk, notF*, and *bvnB*) were created through Gibson assembly (Figure 1a).^39,40,41^ A *pRSFDuet-1* plasmid contained genes encoding NascA and DmtD2/E2, a *pETDuet-1* carried genes for the PhoN and IPK kinases, and a third plasmid, *pACYCDuet-1*, harbored genes for NotF and BvnB. We also routinely employed a *pRARE* plasmid to accommodate rare *E. coli* codons (Figure S8).^42^

Engineering using combinations of the different constructs were performed in *E. coli* BL21 (DE3). To enable straightforward isolation of upstream intermediates, we expressed the enzymes sequentially. As such, in one culture the first two enzymes in the engineered pathway, NascA and DmtD2/E2, were expressed and compounds **1** and **2** were detected via LC-MS (Figure S21-22). In the cultures where all pathway enzymes except for BvnB were expressed, product **3** was detected along with analog **7** from the shunt pathway (Figures 1c, S23-24). Similarly, **8** was identified in the cultures where all the enzymes were expressed (Figure S26). These results were expected as **1** and **7** are the native substrates of NotF and BvnB, respectively. As in the native brevianamide biosynthetic pathway, compound **8** functions as a terminal shunt metabolite in our engineered pathway because it lacks the necessary oxidation state for subsequent tautomerization in the IMDA reaction (Figure S3). Further analysis through LC-MS revealed production of preferred compound **4** in cultures expressing all input biocatalyst genes (Figure S25).

We next sought to confirm the production of pathway intermediates using HPLC. However, in cultures harboring the full enzymatic pathway, BvnB products, **4** and **8** were produced below quantification limit (only detected using LC-MS). To circumvent this, we optimized the culture and gene expression conditions by lowering the incubation temperature from 30 °C to 18 °C and increasing to four days of growth before quenching the reactions (Figures S11-12). Cultivation of engineered *E. coli* in optimized conditions resulted in the detection and isolation of BvnB products, **4** and **8** (Figure 2a, Bottom).

The six purified biocatalytic compounds, **1**-**4** and **7**-**8**, were characterized using HPLC (Figure 2a), MS (Figure S19-30) and NMR (Figure S31-59), with spectra matching previously published literature.^18,43,44,45,46^ Finally, treatment of compound **4** with *ex vivo* synthetic lithium hydroxide readily converts it into the final brevianamide products, **5** and **6** (Figure 2b), through a proposed retro-5-*exo* trig, alkyl shift, and tautomerization cascade (Figure S3).^18^ Overall, introduction of this *de novo* biosynthetic pathway in *E. coli* enabled the isolation of **4** and the production of complex diketopiperazines, **5** and **6**, in a 46% yield and a diastereomeric ratio of 92:8 from endogenous and commercially available starting materials.

### Engineering NADPH Upregulation via a *pfkA* Knock-Out Enhances Yields of Pathway Products

During fermentation scale-up and compound isolation, we quantified the purified yields of compounds **3**–**4** and **7**–**8** and determined the product ratios for **3**:**7** with NotF and **4**:**8** with BvnB (Figure 3a). These results demonstrate that NotF preferentially converts compound **2** into **3**, albeit by a slightly greater margin than its conversion of **1** into **7**. In contrast, BvnB exhibits the opposite preference, favoring the production of compound **8** over **4** in a ratio of 2.5:1. This led us to investigate BvnB’s preference for the saturated substrate **8**, which lacks the Cα–Cβ desaturation.

**Figure 3.**
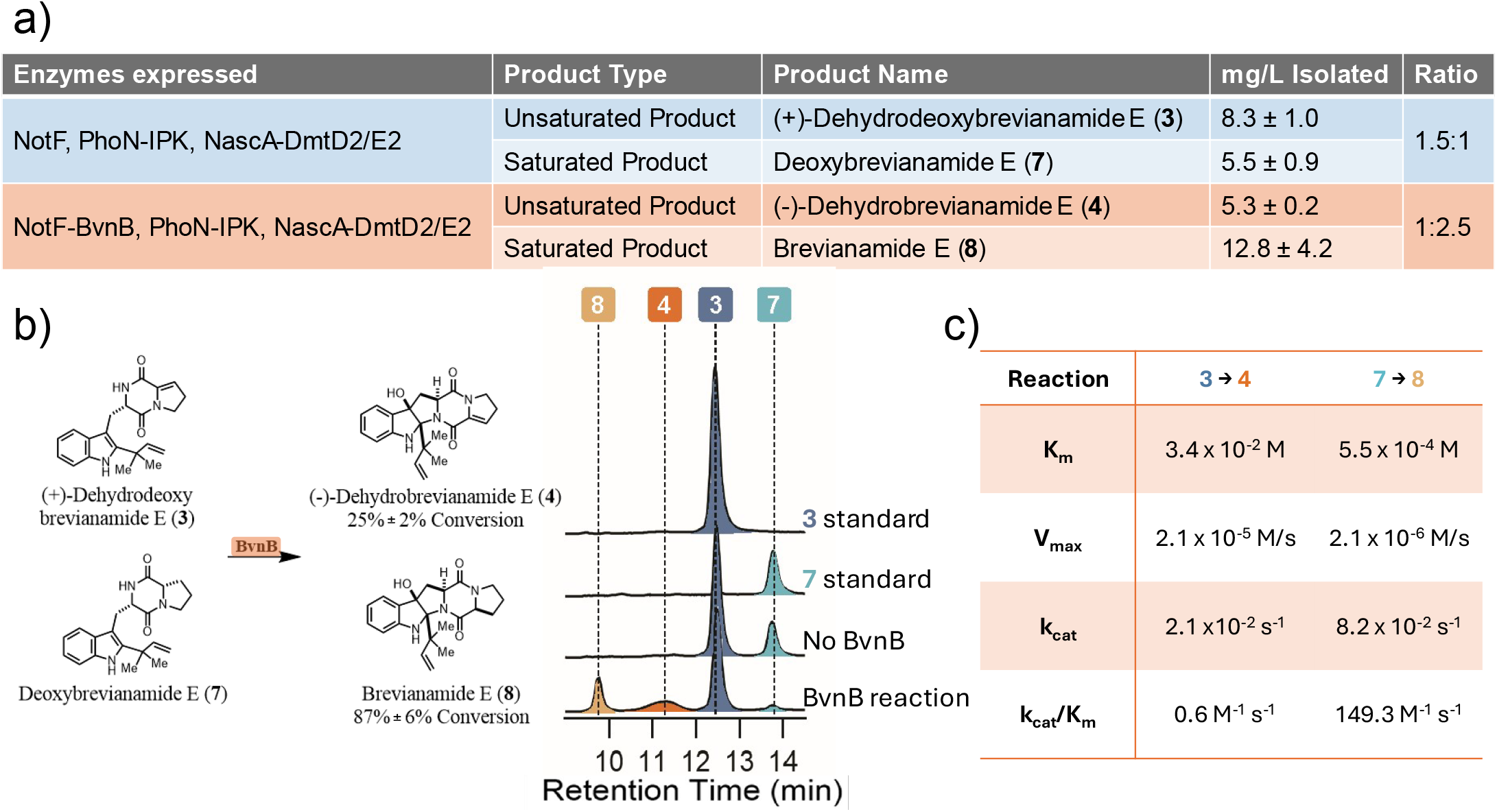
Assessment of BvnB’s preference for **7. a**) Quantification of unsaturated and saturated NotF and BvnB products from the engineered biosynthetic pathway. **b**) In tandem biocatalytic reactions with BvnB through incubation with compounds **3** and **7**. HPLC traces of (i) 400 µM standard of **3** (ii) 400 µM standard of **7** (iii) No enzyme control (iv) BvnB reaction. **c**) Kinetic parameters for BvnB reactions.

To determine the extent of BvnB’s preference for **3** vs **7**, we performed *in vitro* assays testing BvnB’s conversion of **3** to **4** compared to its conversion of **7** to **8**. First, we incubated each substrate with BvnB in separate reactions and observed comparable conversion for both reactions (Figure S13). However, when we incubated **3** and **7** with BvnB in the same reaction, we observed an 87% conversion of **7** to **8** and a 24% conversion of **3** to **4** (Figure 3b). Further corroboration came from BvnB kinetic studies, which revealed a k_cat_/K_m_ for **3** approximately two orders of magnitude less than **7** (Figures 3c, S14). BvnB’s preference for **7** over **3** aligns with the *in vivo* results and underscores the role of this FMO as a bottleneck in the pathway. This finding also illustrates the imbalance in enzyme substrate promiscuity within biosynthetic pathways, as NotF more readily converts non-native substrates into high-yielding products compared to BvnB.^47,48^,^20^

To enhance the efficiency of BvnB, we reexamined the FMO’s mechanism, which relies on a redox system composed of an enzyme-bound FAD and NADPH cofactors.^49^ We tested the effect of NADPH on BvnB production of **4** *in vitro* and found higher concentrations of NADPH correlated with greater conversion of **3** into compound **4** (Figure S15). These findings prompted us to upregulate NADPH in our engineered *E. coli* strain.

Several strategies exist to upregulate NADPH *in vivo*^50–53^, including a genetic knock-out (KO) of the first committed step in glycolysis: conversion of fructose-6-phosphate to fructose-6-bisphosphate by phosphofructokinase (PFK).^54^ This KO would redirect metabolic flux towards the oxidative portion of the pentose phosphate pathway (ox-PPP), thus providing higher levels of NADPH within the cell (Figure 4a). We employed a *pEcCas*/*pEcgRNA* CRISPR-Cas9 system^55^ from Li *et al*. as a strategy for complete deletion of the dominant PFK gene, *pfkA*, in the *E. coli* genome.

**Figure 4.**
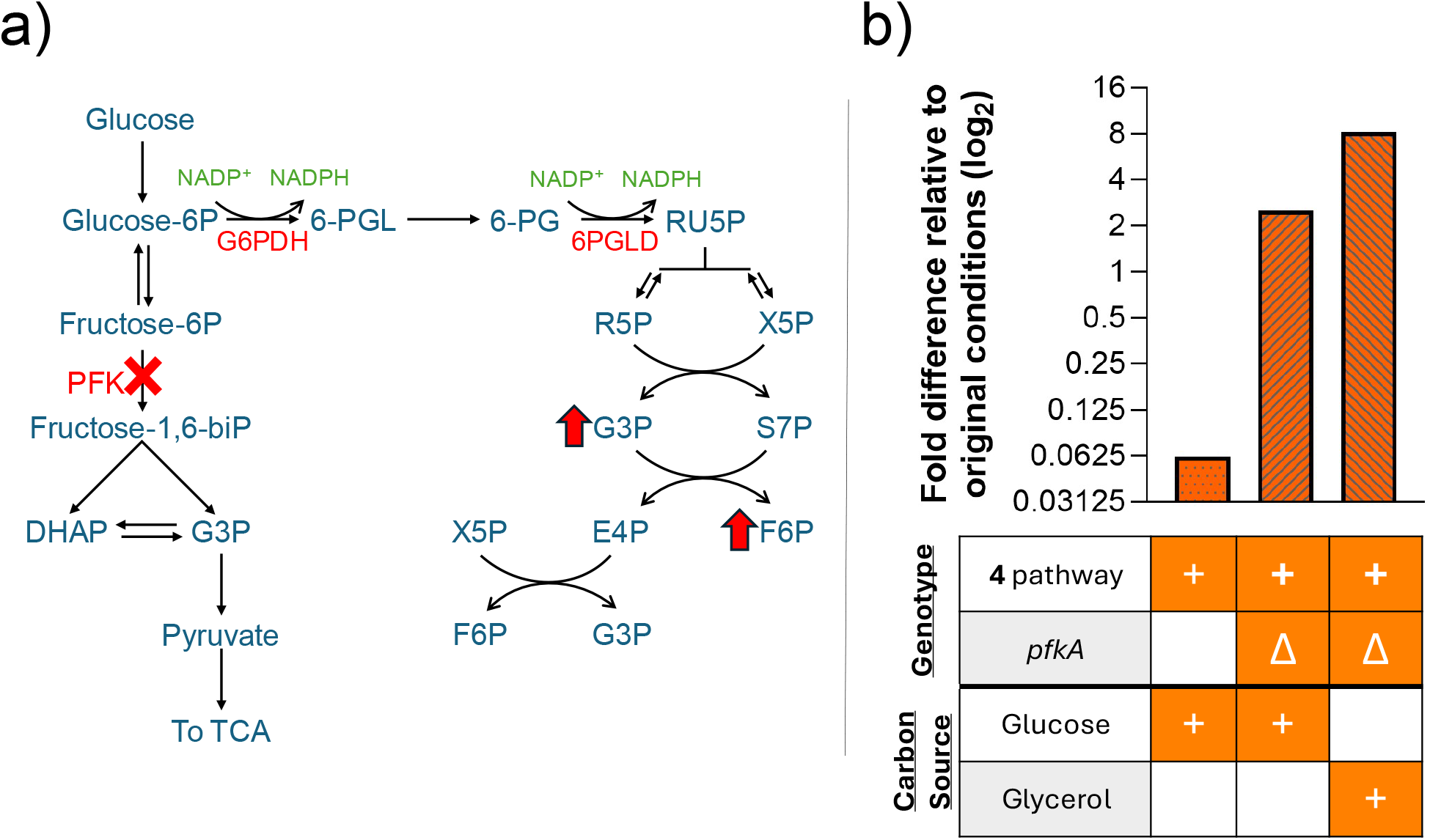
NADPH upregulation strain via CRISPR-Cas9 knock-out of dominant phosphofructokinase (*pfk*) gene. **a**) Simplified glycolysis and pentose phosphate pathway, highlighting NADPH production through the oxidative branch of the PPP after knock-out of *pfk*. Red arrows indicate proposed upregulation induced by addition of glycerol. Glucose-6P = Glucose-6-phosphate, Fructose-6P = F6P = Fructose-6-phosphate, Fructose-1,6-biP = Fructose-1,6-bisphosphate, DHAP = dihydroxyacetone phosphate, G3P = glyceraldehyde-3-phosphate, TCA = tricarboxylic acid, G6PDH = Glucose-6-phosphate dehydrogenase, 6-PGL = 6-phosphogluconolactone, 6-PG = 5-phosphogluconate, 6PGLD = 6-phosphogluconolactonase dehydrogenase, RU5P = ribulose-5-phosphate, R5P = ribose-5-phosphate, X5P = xylulose-5-phosphate, S7P = sedoheptulose-7-phosphate, E4P = erythrose-4-phosphate. **b**) Average AUC fold differences for the production of **4** across conditions relative to the original engineered biosynthetic pathway strain under glycerol conditions. Data presented in a log_2_ scale. **4** pathway refers to the expression of all enzymes and Δ refer to the KO of *pfkA*.

Initially, we used glycerol as the carbon source for all cultivations to ensure rapid growth of *E. coli* cells, especially since they are cultured in the presence of four antibiotics to select each plasmid.^52,56^ Glycerol has also been shown to promote viability of host cells compared to glucose.^57^ However, we envisioned that the Δ*pfkA* mutant would require glucose as a carbon source in order to feed into the ox-PPP. We compared the production of downstream pathway intermediates between the original strain harboring the engineered biosynthetic pathway and *ΔpfkA* mutant strains, expressing all enzymes in glucose and glycerol media. In the first day of incubation, the Δ*pfkA* strain exhibited the expected trend, producing higher levels of compound **4** in glucose compared to glycerol media (Figure S17). However, after four days of fermentation, we enhanced the production of **4** using the Δ*pfkA* mutant by eight-fold compared to the original strain with glycerol as the preferred carbon source. We speculate that the shift in carbon source preference is attributed to glycerol’s ability to metabolize into glucose-3-phosphate, which feeds back into the PPP and acts as a substrate for generating fructose-6-phosphate (Figure 4a). Fructose-6-phosphate can be converted into glucose-6-phosphate, which enters the ox-PPP generating NADPH. The delay in the production of NADPH in glycerol media compared to glucose media may be due to the direct route that glucose takes to enter the PPP.

Furthermore, the mutant strain produced greater amounts of compounds **3** and **8** (Figure S18), mirroring the trend of **4** production. This suggests that upregulation of NADPH not only benefits BvnB, but also enhances overall cellular production of the desired metabolites. It is likely that BvnB competes against native enzymes that also consume NADPH. Once this co-factor is upregulated, there is sufficient availability for both BvnB catalysis and cell growth leading to elevated enzyme production and pathway intermediates.

## Conclusion

In this study, we report the design of a five-step engineered biosynthetic pathway starting from simple precursors to generate 5.3 mg/L of (-)-dehydrobrevianamide E (**4**) devoid of racemization. Subsequently, compound **4** underwent an aqueous synthetic reaction to produce (+)-brevianamides A (**5**) and B (**6**), in a 46% yield and with a 92:8 diastereomeric ratio. This closely resembles the product profile ratio from the native fungal strain^15^ and is in agreement with previous energetic studies on **5** and **6**.^15–17^ Using our engineered biosynthetic pathway, we can generate compounds **5** and **6** from **4** in amounts comparable to the titers produced by the native *P. brevicompactum* pathway (Table S3). In addition, we showed that NADPH upregulation through *pfkA* gene KO leads to the increased production of precursor **4** by eight-fold.

To engineer an *E. coli* strain capable of producing brevianamides **5** and **6**, we selected enzymes with known substrate flexibility. Specifically, we utilized a CDPS and a CDO to optimize precursor assembly. Though the use of a CDO deviates from a strict biomimetic approach for brevianamide assembly, we were motivated to pursue this alternative path to overcome known restrictions on expression of fungal-derived P450s in bacteria. We also included an endogenous source of DMAPP that avoided incorporation of 1-deoxy-D-xylulose 5-phosphate or metabolic engineering of the mevalonate pathway.^35,36^ Given these considerations, we combined enzymes from different biosynthetic pathways, and across the biological kingdoms of Bacteria, Fungi, and Archaea using heterologous expression in *E. coli*. While a previous study has utilized the strategy of co-expressing bacterial CDPS and fungal prenyltransferase genes^58^, our study extends further to access highly complex indole alkaloids. Our findings highlight the significant degree of flexibility available for *de novo* pathway design when specific roadblocks must be overcome.

Broad efforts are currently underway to obtain previously inaccessible complex molecules using directed evolution^59–61^, whole BGC heterologous expression^62–64^, and combinatorial biosynthetic approaches.^48,65–67^ Our pathway is amenable to further diversification due to its modular design, which enables ready substitution of CDPS and CDO enzymes with varying substrate preferences. We observed production of saturated congeners of the native substrates for NotF and BvnB that reflect the flexibility of these biocatalysts. Efforts to develop this system further by generating a diverse range of DKPs in a high-throughput fashion through both DKP feeding studies and combinatorial biosynthetic strategies are in progress and could lead to the formation of bioactive and life-saving molecules.

## Supporting Information

Experimental details, materials, and methods including NMR spectra, HPLC and MS traces, and figures (SI).

## Author Information

### Notes

The authors declare no competing financial interest.

## Acknowledgements

We are grateful for support from NIH grant R35GM118101 and the Hans W. Vahlteich Professorship (to D.H.S.). The authors acknowledge Rajani Arora, UM LSI Multimedia and Social Media specialist, for her contribution to figures in this journal article, Natalia Harris for her assistance in editing the document, and the Summer Research Opportunity Program (SROP) for funding co-author Makayla Perez. Research reported in this publication was supported by the University of Michigan BioNMR Core Facility (U-M BioNMR). U-M BIoNMR Core is grateful for support from U-M including the College of Literature, Sciences and Arts, Life Sciences Institute, College of Pharmacy and the Medical School along with the U-M Biosciences Initiative.

This article references 67 other publications.

## Notes

### Competing Interest Statement

The authors have declared no competing interest.

### Summary of Updates

Formatting changes, particularly to Figure 2, are corrected.

