## Supplementary Information for "Engineering a Biosynthetic Pathway for the Production of (+)-Brevianamides A and B in *Escherichia coli*"

### Supporting Information

Table of contents

**Instrumentation** ........ S4
**Construct design** ........ S4
**Supplementary Table 1.** Primers used in this study ........ S5
**Partially codon-optimized sequence of *dmtD2/E2* used in this study** ........ S6
**Supplementary Table 2.** Expected [M+H] of pathway compounds ........ S6
**Supplementary Figure 1.** Previous work exploring the synthesis of hydroxypyrrolindolines ........ S7
**Supplementary Figure 2.** Native Brevianamide Pathway[6] ........ S7
**Supplementary Figure 3.** Mechanism of LiOH reaction with (-)-dehydrobrevianamide E (**4**) as proposed by Lawrence and co-workers[4] ........ S8
**Supplementary Figure 4.** *In vivo* production of brevianamide F (**1**) by CDPS, NascA. ........ S8
**Whole cell biotransformation of brevianamide F (1) with DmtD2/E2** ........ S8
**Supplementary Figure 5.** *In vivo* conversion of brevianamide F (**1**) to dehydrobrevianamide F (**2**). ........ S9
**Whole cell biotransformation of brevianamide F (1) with NotF and PhoN/IPK kinases** ........ S9
**Supplementary Figure 6.** HPLC traces showing *in vivo* conversion of brevianamide F (**1**) to deoxybrevianamide E (**7**) and brevianamide E (**8**) through co-expression of PhoN-IPK with NotF and NotF-BvnB, respectively. ........ S9
**Reaction conditions for *in vitro* biocatalytic cascades (adapted from Kelly *et al.*)**[3] ........ S10
**Supplementary Figure 7.** *In vitro* reaction of NotF and BvnB through incubation with **2** with and without the addition of NADPH using a cellulose column. ........ S10
**Supplementary Figure 8.** Effect of *pRARE* on NotF activity in *E. coli* ........ S11
**General fermentation and engineered biosynthetic pathway intermediate isolation procedures** ........ S11
Characterization of brevianamide F (**1**). ........ S12
Characterization of dehydrobrevianamide F (**2**). ........ S12
Characterization of (+)-dehydrodeoxybrevianamide E (**3**). ........ S13
Characterization of deoxybrevianamide E (**7**). ........ S13
Characterization of (-)-dehydrobrevianamide E (**4**). ........ S14
Characterization of brevianamide E (**8**). ........ S14
**Isolation of (+)-Brevianamides A and B from native organism *Penicillium brevicompactum* (adapted from Ying *et al.*)**[6] ........ S14
Characterization of (+)-brevianamide A (**5**) isolated from native fungi. ........ S15
Characterization of (+)-brevianamide B (**6**) isolated from native fungi. ........ S15
**Lithium hydroxide reaction of (-)dehydrobrevianamide E (4).** ........ S16
Characterization of (+)-brevianamide A (**5**) isolated from LiOH reaction. ........ S16
Characterization of (+)-brevianamide B (**6**) isolated from native fungi. ........ S16
**Supplementary Figure 9.** Individual HPLC traces of *in vivo* engineered pathway intermediates with standards. ........ S18
**Supplementary Figure 10.** HPLC traces of *in vivo* and *in vitro* reactions using unoptimized BvnB conditions at 2 days incubation at 30 °C with a phenylhexyl column. ........ S19

**Supplementary Figure 11.** Temperature optimization ........ S19
**Supplementary Figure 12.** Incubation time optimization ........ S20
**Supplementary Figure 13.** Percent conversion results. ........ S21
**BvnB kinetic assay conditions for reactions with 3 and 7** ........ S22
**Supplementary Figure 14.** Kinetic Data from BvnB reactions with **3** and **7**. ........ S23
**Supplementary Figure 15.** Concentration-dependent NADPH effect on BvnB activity. ........ S24
***pfkA* CRISPR-Cas9 knock-out** ........ S24
***pfkA* HDR template** ........ S25
**Colony PCR product sequencing data for *pfka* KO** ........ S25
**Supplementary Figure 16.** DNA gel of colony PCR product for KO of *pfkA*. A) DNA ladder, B) colony PCR product. Expected size ~ 1600bp. ........ S26
**Supplementary Figure 17.** Fold differences of **4** (a), **8** (b), and **3** (c) in different conditions relative to original conditions (**4** pathway in glycerol) after **a single day** of incubation. ........ S27
**Supplementary Figure 18.** Fold differences of **4** (a), **8** (b), and **3** (c) in different conditions relative to original conditions (**4** pathway in glycerol) after **four days** of incubation. ........ S28
**Supplementary Table 3.** Titers (mg/L) of (+)-brevianamides A (**5**) and B (**6**) using native pathway and LiOH reaction ........ S28
**Supplementary Figure 19.** Mass spectrum (TOF LC/MS) of **3** from NotF *in vitro* cascade supplemented with **2**. ........ S28
**Supplementary Figure 20.** Mass spectrum (TOF LC/MS) of **4** from NotF-BvnB *in vitro* cascade supplemented with **2**. ........ S29
**Supplementary Figure 21.** Mass spectrum (TOF LC/MS) of brevianamide F (**1**). ........ S29
**Supplementary Figure 22.** Mass spectrum (TOF LC/MS) of dehydrobrevianamide F (**2**). ........ S29
**Supplementary Figure 23.** Mass spectrum (TOF LC/MS) of (+)-dehydrodeoxybrevianamide E (**3**). ........ S30
**Supplementary Figure 24.** Mass spectrum (TOF LC/MS) of deoxybrevianamide E (**7**). ........ S30
**Supplementary Figure 25.** Mass spectrum (TOF LC/MS) of (-)-dehydrobrevianamide E (**4**). ........ S31
**Supplementary Figure 26.** Mass spectrum (TOF LC/MS) of brevianamide E (**8**). ........ S31
**Supplementary Figure 27.** Mass spectrum (TOF LC/MS) of (+)-brevianamide A (**5**) from fungi extraction... S32
**Supplementary Figure 28.** Mass spectrum (TOF LC/MS) of (+)-brevianamide B (**6**) from fungi extraction. . S32
**Supplementary Figure 29.** Mass spectrum (TOF LC/MS) of (+)-brevianamide A (**5**) from LiOH reaction. .... S32
**Supplementary Figure 30.** Mass spectrum (TOF LC/MS) of (+)-brevianamide B (**6**) from LiOH reaction. .... S33
**Supplementary Figure 31.** $^{1}$H NMR spectrum of brevianamide F (**1**) (599 MHz, $CDCl_3$) ........ S33
**Supplementary Figure 32**. $^{13}$C NMR spectrum of brevianamide F (**1**) (151 MHz, $CDCl_3$) ........ S34
**Supplementary Figure 33.** $^{1}$H NMR spectrum of dehydrobrevianamide F (**2**) (599 MHz, $CDCl_3$) ........ S34
**Supplementary Figure 34.** $^{13}$C NMR spectrum of dehydrobrevianamide F (**2**) (151 MHz, $CDCl_3$) ........ S35
**Supplementary Figure 35.** COSY NMR spectrum of dehydrobrevianamide F (**2**) ($CDCl_3$) ........ S35
**Supplementary Figure 36.** HSQC NMR spectrum of dehydrobrevianamide F (**2**) ($CDCl_3$) ........ S36
**Supplementary Figure 37.** HMBC NMR spectrum of dehydrobrevianamide F (**2**) ($CDCl_3$) ........ S36
**Supplementary Figure 38.** NOESY NMR spectra of dehydrobrevianamide F (**2**) ($CDCl_3$) ........ S37
**Supplementary Figure 39.** $^{1}$H NMR spectrum of dehydrobrevianamide F (**2**) (DMSO-d6) ........ S37
**Supplementary Figure 40.** $^{13}$C NMR spectrum of dehydrobrevianamide F (**2**) (DMSO-d6) ........ S38
**Supplementary Figure 41.** COSY NMR spectrum of dehydrobrevianamide F (**2**) (DMSO-d6) ........ S38
**Supplementary Figure 42.** HSQC NMR spectra of dehydrobrevianamide F (**2**) (DMSO-d6) ........ S39
**Supplementary Figure 43.** HMBC NMR spectra of dehydrobrevianamide F (**2**) (DMSO-d6) ........ S39
**Supplementary Figure 44.** $^{1}$H NMR spectrum of (+)-dehydrodeoxybrevianamide E (**3**) (599 MHz, $CDCl_3$).. S40
**Supplementary Figure 45.** $^{13}$C NMR spectrum of (+)-dehydrodeoxybrevianamide E (**3**) (151 MHz, $CDCl_3$) S40
**Supplementary Figure 46.** $^{1}$H NMR spectrum of deoxybrevianamide E (**7**) (599 MHz, $CDCl_3$) ........ S41
**Supplementary Figure 47.** $^{13}$C NMR spectrum of deoxybrevianamide E (**7**) (151 MHz, $CDCl_3$) ........ S41
**Supplementary Figure 48.** $^{1}$H NMR spectrum of (-)-dehydrobrevianamide E (**4**) (599 MHz, $CDCl_3$) ........ S42
**Supplementary Figure 49.** $^{13}$C NMR spectrum of (-)-dehydrobrevianamide E (**4**) (151 MHz, $CDCl_3$) ........ S42

**Supplementary Figure 50.** $^{1}$H NMR spectrum of brevianamide E (**8**) (599 MHz, $CDCl_3$) ... S43
**Supplementary Figure 51.** $^{13}$C NMR spectrum of brevianamide E (**8**) (151 MHz, $CDCl_3$) ... S43
**Supplementary Figure 52.** $^{1}$H NMR spectrum of (+)-brevianamide A (5) from fungal extraction (599 MHz, $CDCl_3$) ... S44
**Supplementary Figure 53.** $^{13}$C NMR spectrum of (+)-brevianamide A (**5**) from fungal extraction (151 MHz, $CDCl_3$) ... S44
**Supplementary Figure 54.** $^{1}$H NMR spectrum of (+)-brevianamide B (**6**) from fungal extraction (800 MHz, DMSO-d6) ... S45
**Supplementary Figure 55.** $^{13}$C NMR spectrum of (+)-brevianamide B (**6**) from fungal extraction (201 MHz, DMSO-d6) ... S45
**Supplementary Figure 56.** $^{1}$H NMR spectrum of (+)-brevianamide B (**6**) from fungal extraction (599 MHz, $CDCl_3$) ... S46
**Supplementary Figure 57.** $^{1}$H NMR spectrum of (+)-brevianamide A (**5**) from LiOH reaction (599 MHz, $CDCl_3$) ... S46
**Supplementary Figure 58.** $^{13}$C NMR spectrum of (+)-brevianamide A (**5**) from LiOH reaction (151 MHz, $CDCl_3$) ... S47
**Supplementary Figure 59.** $^{1}$H NMR spectrum of (+)-brevianamide B (**6**) from LiOH reaction (599 MHz, $CDCl_3$) ... S47
**References** ... S48

**Instrumentation**
Analytical HPLC data was acquired using a Shimadzu HPLC system comprised of two LC-20ADXR pumps, a SIL-20ACXR autosampler, and an SPD-M20A diode array detector. Preparatory HPLC was performed using a Shimadzu HPLC system comprised of two LC-20AT pumps, a DGU-20A3R degassing unit, SPD-M20A diode array detector, and FRC-10A fraction collector. All NMR spectra were collected on a Bruker 600 spectrometer with Magnex 600/54 active shielded premium magnet, Bruker Prodigy ($^{1}H/^{19}F$)-X broadband probe with Automated Tuning/Matching (ATMA) unit, and Bruker NEO600 NMR System console or a Bruker 800 spectrometer with an Ascend magnet, 24 position SampleCASE and a Bruker NEO console. $^{1}H$ spectra are reported in parts per million on the scale, relative to residual solvent peaks (DMSO-d6: 2.50 or $CDCl_3$). Data are reported as follows: chemical shift [multiplicity (s = singlet, d = doublet, t = triplet, m = multiplet), coupling constant(s) in Hertz, integration]. $^{13}C$ NMR spectra are reported in parts per million on the scale, relative to residual solvent peaks (DMSO-d6: 39.52 or $CDCl_3$: 77.16). Data are reported as follows: chemical shift. High resolution mass spectra (HRMS) were recorded on an Agilent G6545A quadrupole time-of-flight (QTOF) mass spectrometer via electrospray ionization or Agilent G6230B time-of-flight (TOF) mass spectrometer equipped with dual Agilent jet stream electrospray ionization (Dual AJS ESI) source. Optical rotations were obtained using a Jasco P2000 polarimeter at 24 °C with a 100 mm cell. Analytical thin layer chromatography (TLC) was performed using glass plates pre-coated with 0.25 mm 230–400 mesh silica gel impregnated with a fluorescent indicator (254 nm). TLC plates were visualized by exposure to short wave ultraviolet light (254 nm) and long wave ultraviolet light (365 nm). Normal phase flash chromatography was performed on a Biotage Isolera One with Silia Sep Silica HP Flash Cartridges and Biotage Sfär DLV columns.

**Construct design**
The coding sequences of *notF* (accession #: E0Y3X1.1), *bvnB* (accession #: QFZ94975), *phoN*, and *ipk* were cloned into *pET28b*, and *pETDuet-1* as previously described.[1,2] *nascA* was cloned into *pET28b* after amplification from *Streptomyces sp.* CMB-MQ030 gDNA. *dmtD2/dmtE2* (hereon referred to as *dmtD2/E2*) was codon optimized for *E. coli*, but the overlapping region between CDOA and CDOB was not optimized and ordered as a *pET28a* construct from Twist Biosciences.

For cloning into *pACYCDuet-1*, *notF* and *bvnB* were amplified using the primers listed in Table S1. The PCR reaction contained 10X Pfu Ulta II buffer, 0.2 mM dNTPs, 0.4 µM of each primer, 200 ng template DNA, 1 µL Pfu Ultra II DNA polymerase in a total volume of 50 µL. The cycling conditions used were as follows: 1) 95 °C, 3 min; 2) 95 °C, 30 s; 3) 66 °C for *notF*/62 °C for *bvnB*, 1 min; 4) 72 °C, 3 min for *notF*/5 min 20 s for *bvnB*; 5) repeat steps 2-4 30x; 6) 72 °C, 15 min; 7) 4 °C, hold. After amplification the products were purified via PCR cleanup and gel purified (Invitrogen), *notF* was inserted into 'empty' *pACYCDuet-1*, and *bvnB* into *pACYCDuet-1-notF* treated with EcoRI and HindIII for MCS1 and NdeI and XhoI for MCS2 via Gibson Assembly (NEB).

For cloning into *pRSFDuet-1*, *nascA* and *dmtD2/E2* were amplified using the primers listed in Table S1. The PCR reaction contained 10X Pfu Ulta II buffer, 0.2 mM dNTPs, 0.4 µM each primer, 200 ng template DNA, 1 µL Pfu Ultra II DNA polymerase in a total volume of 50 µL. The cycling conditions used were as follows: 1) 95 °C, 3 min; 2) 95 °C, 30 s; 3) 68 °C for *nascA*/64 °C for *dmtD2/E2*, 1 min; 4) 72 °C, 1 min 33 sec for *nascA*/2 min for *dmtD2/E2*; 5) repeat steps 2-4 30x; 6) 72 °C, 15 min; 7) 4 °C, hold. After amplification the products were purified via PCR cleanup and gel purified (Invitrogen), nascA was inserted into 'empty' *pRSFDuet-1*, and *dmtD2/E2* into *pRSFDuet-1-nascA* treated with EcoRI and HindIII for MCS1 and NdeI and XhoI for MCS2 via Gibson Assembly (NEB).

NotF and BvnB enzymes were sourced from previously purified stocks.[3]

Single-stranded oligos from Table S1 were ligated together using the following conditions: 5 µL 20 mM each sgRNA oligo, 5 µL T4 ligase in a total volume of 50 µL. The cycling conditions used were as follows: 1) 95 °C, 5 min; 2) steady 10 °C/min decrease till the temperature reached 16 °C; 3) hold at 16 °C, 10 min. The double-stranded oligo product was diluted 200-fold before ligated into BsaI-linearized pEcgRNA with the following conditions:  2 µL 10X T4 DNA ligase buffer, 50 ng pEcgRNA, 37.5 ng double-standed oligo, and 1 µL of T4 DNA ligase in total volume of 20 µL which was incubated at 16 °C for 1 hour.

To create the *pfkA* upstream and *pfkA* downstream fragments for the HDR template, we used the primers listed in Table S1. The PCR reaction contained 10X Pfu Ulta II buffer, 0.2 mM dNTPs, 0.5 µM each primer, 200 ng template DNA, 0.5 µL Pfu Ultra II DNA polymerase in a total volume of 50 µL. The cycling conditions used were as follows: 1) 98 °C, 30 s; 2) 98 °C, 10 s; 3) 58 °C for *pfkA* upsteam/68 °C for *pfkA* downstream, 10 s; 4) 72 °C, 10 s; 5) repeat steps 2-4 30x; 6) 72 °C, 5 min; 7) 4 °C, hold. After amplification the products were purified via gel extraction (Qiagen).

We used overlap extension PCR to combine *pfkA* upstream and *pfkA* downstream fragments and create the HDR template. The PCR reaction first contained 10X Pfu Ulta II buffer, 0.2 mM dNTPs, 50 ng of each fragment, and 0.5 µL Pfu Ultra II DNA polymerase in a total volume of 50 µL. After 10 extension cycles, 0.2 µM each of *pfka* upstream forward primer and *pfka* downstream reverse primer (Table S1) were added to the reaction. The full cycling conditions were as follows: 1) 98 °C, 30 s; 2) 98 °C, 10 s; 3) 70 °C 10 s; 4) 72 °C, 15 s; 5) repeat steps 2-4 10x, add primers, and then repeat steps 2-4 20x; 6) 72 °C, 5 min; 7) 4 °C, hold. After amplification the products were purified via gel extraction (Qiagen).

Colony PCR was performed to identify colonies with the *pfkA* KO. 10X Standard *Taq* Reaction Buffer, 0.2 mM dNTPs, 0.2 µM each primer (Table S1), 200 ng template DNA, 0.1 µL *Taq* DNA polymerase in a total volume of 20 µL. The cycling conditions used were as follows: 1) 95 °C, 30 s; 2) 95 °C, 30 s; 3) 55 °C, 30 s; 4) 68 °C, 1 min and 40 sec; 5) repeat steps 2-4 30x; 6) 68 °C, 5 min; 7) 4 °C, hold.

**Supplementary Table 1.** Primers used in this study

| Gene | Primers used to amplify |
|---|---|
| *notF* | Forward: GCCAGGATCCGAATTCGAGCTCGATGACGGCCCCAGAGCTCCG<br>Reverse: CTGTTCGACTTAAGCATTATGCGGCCGCTCAATCTTCTTCCCACAGATAGGTGC |
| *bvnB* | Forward: ATTAGTTAAGTATAAGAAGGAGATATACATatgggcCATCATCATCACCATCACCATCAC<br>Reverse: CGCAGCAGCGGTTTCTTTACCAGACTCGAGCTACTCCTGACGATATTT |
| *nascA* | Forward: GCCAGGATCCGAATTCGAGCTCGGTGAACACTTCCCTCGCTGCGG<br>Reverse: CTTAAGCATTATGCGGCCGCAAGCTTTCAGCGTTCGGCCGCCC |
| *dmtD2/E2* | Forward: GTATAAGAAGGAGATATACATATGGGCAGCAGCCATCATCATCATCATCACAGCAGCGGCC |

| | |
|---|---|
| | Reverse: GTTTCTTTACCAGACTCGAGCTCGAGTTATGGCCGTGGAAGAAGATCGACATCGC |
| sgRNA *pfkA* | Forward: TAGTTCTGACATGATCAACCGTGG Reverse: AAACCCACGGTTGATCATGTCAGA |
| *pfkA* upstream HDR template | Forward: CTATATTTTATATAGCGCGTTACGCATGGGATATGAGGCG Reverse: TCTGCCTTTTTCCGAAATCATTACATGACTACCTCTGAACTTTGG |
| *pfkA* downstream HDR template | Forward: ATGTAATGATTTCGGAAAAAGGCAGATTCCTTTACCCTGAAACCGATGACAG Reverse: TTTACCTGAGCCACCGTGTGACTGACGAATCACCACGTTATC |
| *pfkA* colony PCR | Forward: CTGATGTTATGATGAACGGCG Reverse: GCGAGGATAGACTCACTCGG |

**Partially codon-optimized sequence of *dmtD2/E2* used in this study**
ATGGATGCAGTTGACGCAGGTCCCGCAGGTCCCGCAGGCCCAGCAGGTTCAGCTGACGCCGTT
CTCCGCGTATTACGGAGCCGGAGTGTGGTCCGGCAATACACCGGGCGTCAAGTTGATGATGAG
GTTTTAGAAATGCTCGTTTCTGCAATGTTGGCGGCTCCGACTGCGAGCAATAAACAAGCTTGGG
CGTTTGTGGCTGTGCGCGAGAGACGCACACTGCGTCTTTTGGGTGCATTCGCACCGGGTATAAT
CGGTACGCCCCCACTTGTAGTTGCAGCATGTTTTGACCGTTCCCGCCCAGTCGACGAACGCGGT
CCAGGCGAGGGTGGTTGGGATATGGGTTTGCTGTGTGTAGCGATGGCAGTTGAAAACTTATTAC
TGGCTGCTCATGCTCTCGGCTTAGGCGGGTGCCCTGTCGGCGGGTTCCGGGAAGGGCCAGTC
CGCACAGTGTTAAGACTCCCTGCTCATCTCGACCCTGTGTTATTGGTACCAGTGGGACACCCGG
CGGGTCCCCTGCGGCCTACAGATCGACGTGACCGGAACGAGGTGCTCAGACATGACACATGGG
AGCAGTGACAGCGGCCGTATGCGTGAAGAGCTTTTATTACTTGCCGCATTTCTTTTATCTTCTGG
CCGGGGTTTGGCTGATGAACCTGCGGTATATGGTCAAGCCAGATGTCTTGATGCAGCTCGTCGG
ACCCTCGCATTAGTGGAAGGTTTAGGTGGACAAGATCCCGCGGTAACGGGTTTGAGAACAGAAC
TTGAAGCCTTTATGACAGGACCTATCGGTGGTTGTGGTGATATTAATACCCTTTTAGATAGCGCT
TGTGATCGTCTGGCTGAAGTACTCTGTGATCGAGGGCGCGATGTCGATCTTCTTCCACGGCCAT
AACTCGAG

**Supplementary Table 2.** Expected [M+H] of pathway compounds

| Compound | Chemical Formula | Expected [M+H] |
|---|---|---|
| Brevianamide F (**1**) | C16H17N3O2 + H | 284.1394 |
| Dehydrobrevianamide F (**2**) | C16H15N3O2 + H | 282.1237 |
| Deoxybrevianamide E (**7**) | C21H25N3O2 + H | 352.202 |
| Dehydrodeoxybrevianamide E (**3**) | C21H23N3O2 + H | 350.1863 |
| Brevianamide E (**8**) | C21H24N3O3 + H | 368.1969 |
| Dehydrobrevianamide E (**4**) | C21H23N3O3 + H | 366.1812 |

| |
|---|
| Brevianamide A (**5**)<br>Brevianamide B (**6**) |

**Supplementary Figure 1.** Previous work exploring the synthesis of hydroxypyrrolindolines. a) Lawrence *et al.* synthesis of (+)- and (-)-brevianamides A (**5**) and B (**6**).[4] b) Bisai *et al.* synthesis of **5** and **6**.[5] c) Biocatalytic cascade of NotF-BvnB to produce brevianamide E (**8**).[3]

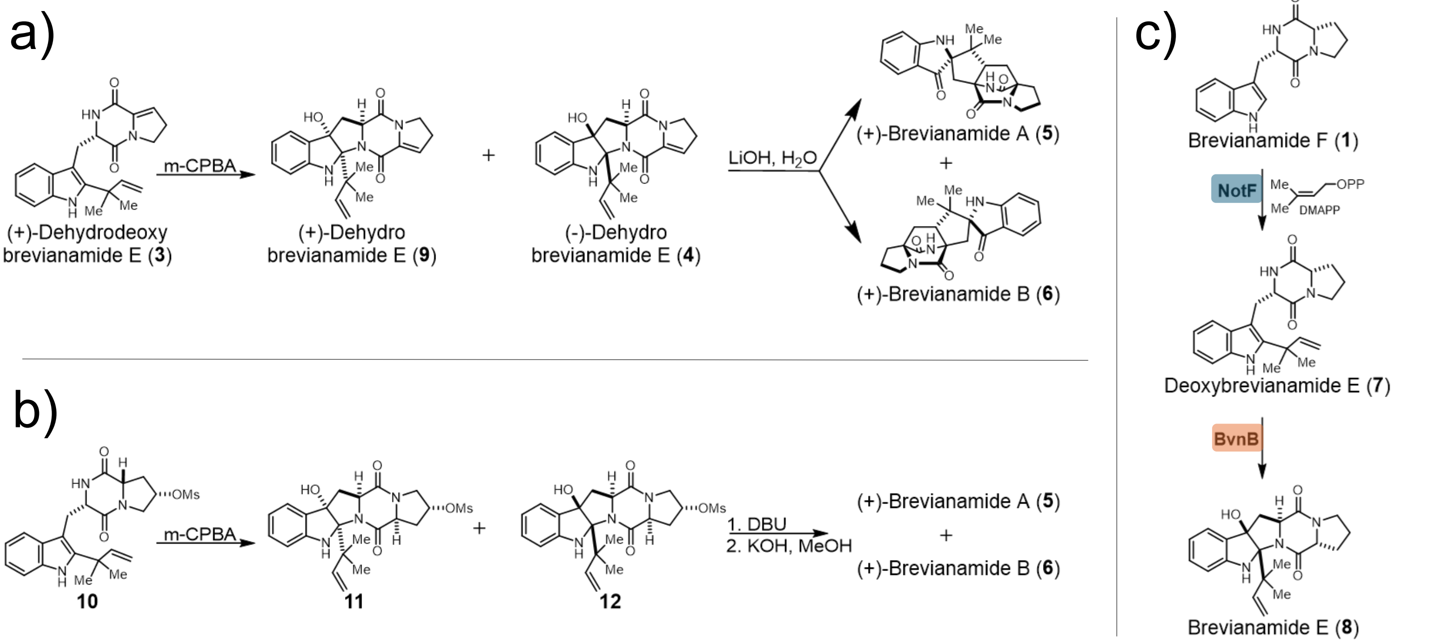

**Supplementary Figure 2.** Native Brevianamide Pathway[6]

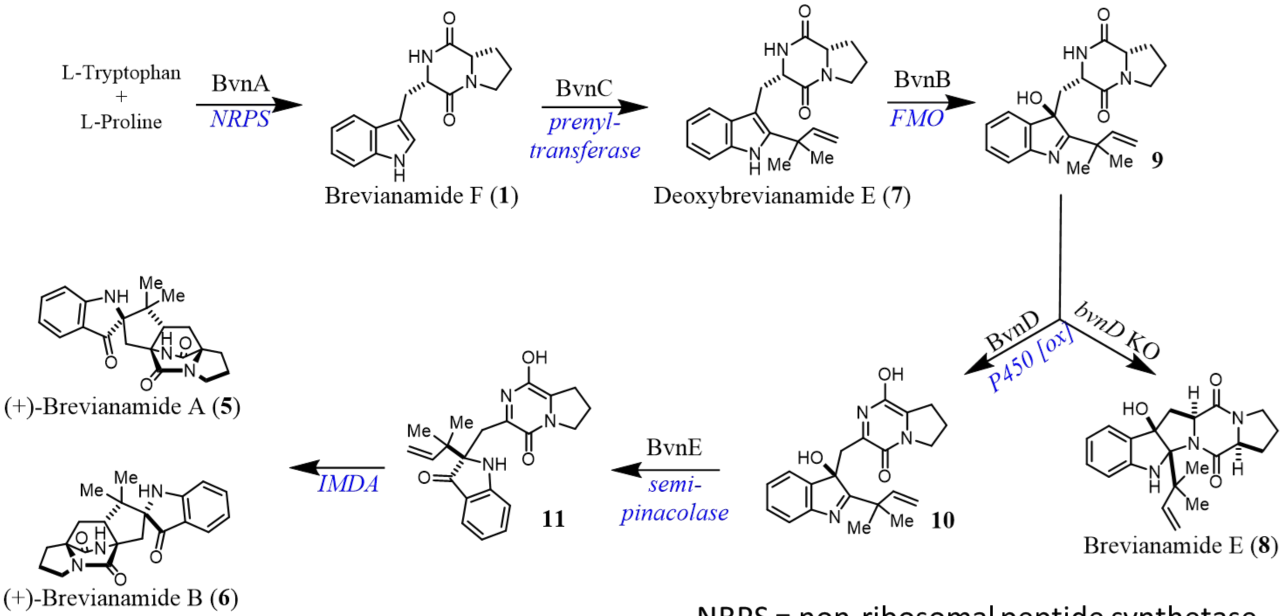

NRPS = non-ribosomal peptide synthetase
FMO = flavin-dependent monooxygenase
IMDA = intramolecular Diels-Alder

**Supplementary Figure 3.** Mechanism of LiOH reaction with (-)-dehydrobrevianamide E (**4**) as proposed by Lawrence and co-workers[4]

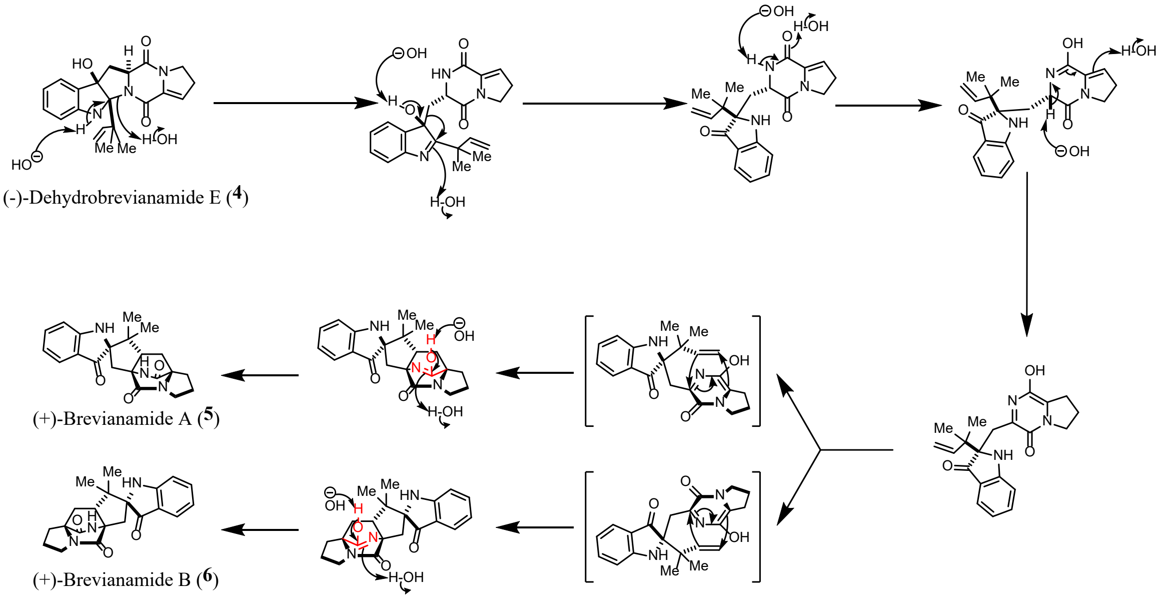

**Supplementary Figure 4.** *In vivo* production of brevianamide F (**1**) by CDPS, NascA. (i) NascA reaction. (ii) **1** standard (iii) MeOH blank.

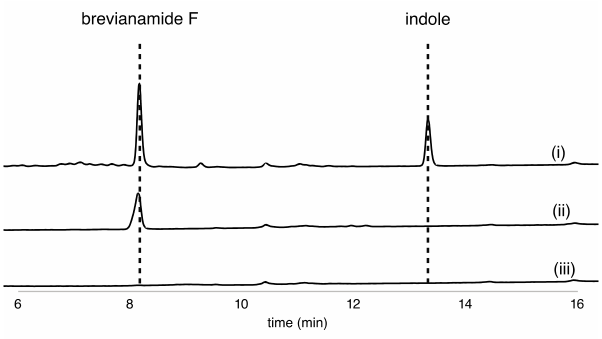

### Whole cell biotransformation of brevianamide F (1) with DmtD2/E2

The *pET28a* plasmid containing *dmtD2/E2* was transformed into chemically competent *E. coli* BL21(DE3) harboring the *pRARE* plasmid for protein overexpression. A single colony was grown in 3 mL SOB (50 µg/mL kanamycin and 50 µg/mL spectinomycin) at 37 °C, and 100 µL of this seed culture was used the following morning to inoculate 10 mL LB (50 µg/mL kanamycin and 50 µg/mL spectinomycin) and grown at 37 °C to OD600 = ~ 0.5. Protein production was induced by addition of 0.4 mM IPTG to a final concentration of 500 µM and 200 µL of 80 mM brevianamide F (**1**) stock in DMSO was added. The protein was expressed overnight at 18 °C. The following morning, the cells were pelleted by centrifugation and the supernatant was quenched with 3x volume methanol, vortexed for 20 s, and chilled at 4 °C for 10 min. The mixture was centrifuged at 17,000 x g for 3 minutes to precipitate solids before injection and purification by preparatory HPLC, using a Lux® cellulose-3 Tris (4-methylbenzoate) semi-preparative HPLC column (5 µm, 250 x 10 mm) and a gradient of 10 – 30% acetonitrile : water at 5 mL/min over 5 min. Fractions containing desired product were pooled and concentrated to obtain **1** as a white solid, clean for NMR characterization.

**Supplementary Figure 5.** *In vivo* conversion of brevianamide F (**1**) to dehydrobrevianamide F (**2**).

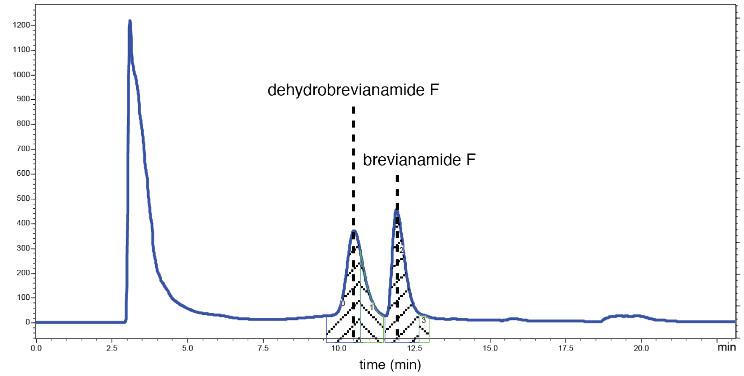

### Whole cell biotransformation of brevianamide F (1) with NotF and PhoN/IPK kinases

The *pET28b-notF* and *pETDuet-1-phoN-ipk* plasmid were co-transformed into chemically competent *E. coli* BL21(DE3) harboring the *pRARE* plasmid for protein overexpression. A single colony was grown in 3 mL SOB (50 µg/mL kanamycin, 100 µg/mL ampicillin, and 50 µg/mL spectinomycin) at 37 °C, and 100 µL of this seed culture was used the following morning to inoculate 10 mL TB (50 µg/mL kanamycin, 100 µg/mL ampicillin, and 50 µg/mL spectinomycin) and grown at 37 °C to OD600 = ~ 0.8. Protein production was induced by addition of 0.4 mM IPTG to a final concentration of 500 µM, prenol to a final concentration of 0.025% v/v, and 100 µL of 40 mM **1** stock in DMSO was added (1.4 mM final concentration). The proteins were expressed for 48 hours at 30 °C. The cells were then pelleted by centrifugation and the supernatant was quenched with 3x volume methanol, vortexed for 20 s, and chilled at 4 °C for 10 min. Prior to HLPC analysis the mixture was centrifuged at 17,000 x g for 3 minutes to precipitate protein, and the supernatant was injected on a Shimadzu HPLC with a Phenomenex Luna® 5µm phenyl hexyl LC column (100 Å, 5 µm, 250 x 4.6 mm) using a linear gradient of 20-80% acetonitrile: water (0.1% formic acid) over 20 min (1.25 mL/min).

**Supplementary Figure 6.** HPLC traces showing *in vivo* conversion of brevianamide F (**1**) to deoxybrevianamide E (**7**) and brevianamide E (**8**) through co-expression of PhoN-IPK with NotF and NotF-BvnB, respectively. (i) Production of **8** through **1** feeding to co-expression of PhoN, IPK, NotF, and BvnB. (ii) Production of **7** through **1** feeding to co-expression of PhoN, IPK, and NotF (iii) **7** standard (iv) **1** standard (v) MeOH blank.

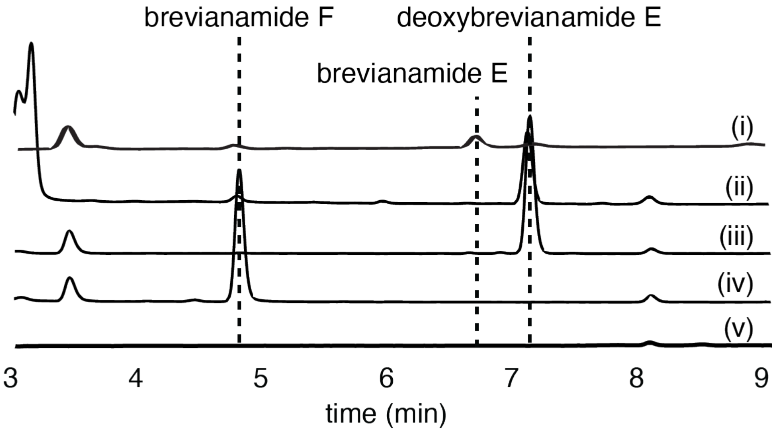

**Reaction conditions for *in vitro* biocatalytic cascades (adapted from Kelly *et al.)*[3]**

The standard enzyme assay containing 5 mM MgCl2, 1.5 mM DMAPP, 1 mM substrate, 5 µM NotF, and 25 µM BvnB in reaction buffer (5% v/v glycerol, 300 mM NaCl, 20 mM Tris pH 7.9) to end volume of 50 µL was initiated with the addition of 5 mM NADPH and incubated at 28 °C overnight for general *in vitro* assays or for 1 h in the case of percent conversion assays. The *in vitro* NotF reactions did not contain NADPH and were initiated with the addition of enzyme. The control reactions for the general *in vitro* assay included all components except for NotF or BvnB. Percent conversion assays contained equimolar (1 mM) amounts of **3** and **7**. Controls for the percent conversion assays included all components except for BvnB. Reactions were quenched with 150 µL HPLC-grade methanol and vortexed at maximum speed for 10 sec. Prior to HLPC analysis, reactions were centrifuged at 17,000 x g for 3 minutes to precipitate protein, and the supernatant from the general *in vitro* assays was injected on a Shimadzu HPLC with a Phenomenex Lux® cellulose-3 Tris (4-methylbenzoate) LC column (100 Å, 5 µm, 100 x 4.6 mm) using a linear gradient of 15-50% acetonitrile: water over 12.5 min (1 mL/min) and a Phenomenex Lux® cellulose-3 Tris (4-methylbenzoate) LC column (100 Å, 5 µm, 250 x 4.6 mm) using a linear gradient of 25-60% acetonitrile: water over 12.5 min (1 mL/min). The supernatant of percent conversion reactions was injected on a Shimadzu HPLC with a Phenomenex Luna® C18(2) LC column (100 Å, 5 µm, 250 x 4.6 mm) using a linear gradient of 35-45 % acetonitrile + 0.1% formic acid: water + 0.1 % formic acid over 15 minutes (1 mL/min) at 40 °C.

**Supplementary Figure 7.** *In vitro* reaction of NotF and BvnB through incubation with **2** with and without the addition of NADPH using a cellulose column. (i) **2** standard (ii) **3** standard (iii) **4** standard (iv) NotF + BvnB + **2** without NADPH (v) NotF + BvnB + **2** + NADPH.

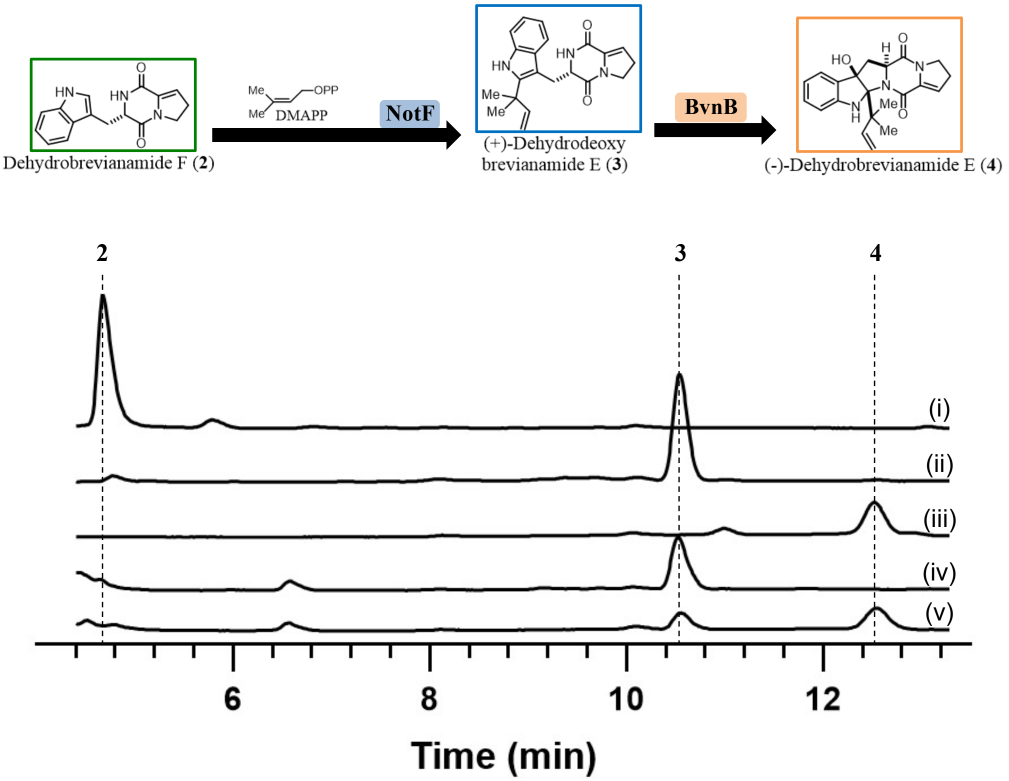

**Supplementary Figure 8.** Effect of *pRARE* on NotF activity in *E. coli*

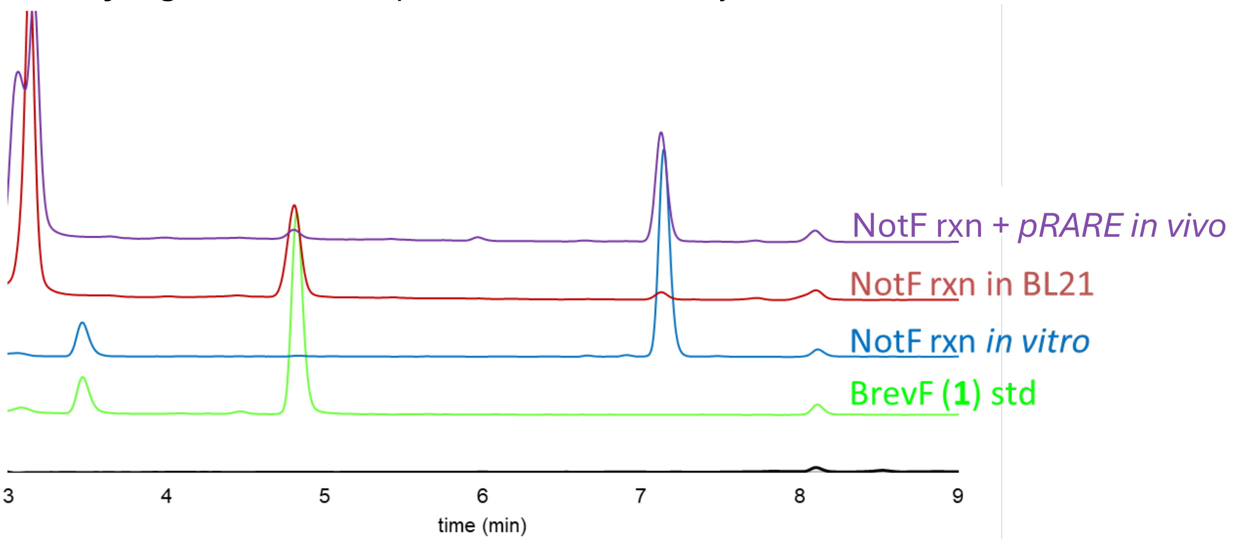

**General fermentation and engineered biosynthetic pathway intermediate isolation procedures**
The plasmid(s) were transformed into chemically competent *E. coli* BL21(DE3) harboring the *pRARE* plasmid and *pETDuet-1-phoN-ipk* plasmid for protein overexpression. A single colony was grown in 3-10 mL SOB with appropriate antibiotics (50 µg/mL kanamycin, 35 µg/mL chloramphenicol in EtOH, 100 µg/mL ampicillin, and 50 µg/mL spectinomycin) at 37 °C, and 500 µL of this seed culture was used the following morning to inoculate TB + 1x potassium phosphate buffer with appropriate antibiotics and grown at 37 °C to OD600 = ~ 0.8. Protein production was induced by addition of 1 M IPTG to a final concentration of 1.25 mM and prenol to a final concentration of 0.025% v/v (if NotF was expressed). The proteins were expressed for 96 hours at 30 °C or 18 °C if BvnB was expressed. The cells were then pelleted by centrifugation and the culture was quenched with 3x volume methanol, vortexed for 20 s, and chilled at 4 °C for 10 min. The mixture was centrifuged at 17,000 x g for 3 minutes to precipitate biomass, and the aqueous layer was extracted with chloroform. The combined organic layers were washed with brine, dried with sodium sulfate, and concentrated.

The dried material was later resuspended in methanol and injected on a Shimadzu HPLC with a Phenomenex Lux® cellulose-3 Tris (4-methylbenzoate) LC column (100 Å, 5 µm, 250 x 4.6 mm) using a linear gradient of 25-60% acetonitrile: water over 12.5 min (1 mL/min) and a Phenomenex Luna® 5 µm phenylhexyl LC column (100 Å, 250 x 4.6 mm) using a linear gradient of 30-95% acetonitrile: water (0.1% formic acid) over 15 min (1 mL/min). The former method was specifically used to separate **3** and **4** while the latter was specifically used to separate **3** and **7**.

Compounds **1** and **2** from NascA-DmtD2/E2 cultures were isolated by injecting a 50-100 mg/mL solution of the extracts in a Shimadzu semi-preparatory HPLC, using a Phenomenex Lux® cellulose-3 Tris (4-methylbenzoate) semi-preparative (5 µm, 250 x 10 mm) HPLC column with an isocratic method of 20% acetonitrile: water at 5 mL/min over 16 min.

Compounds **3** and **7** (and **2**) from NascA-DmtD2/E2, PhoN-IPK, and NotF cultures were isolated by injecting a 50-100 mg/mL solution of the extracts in a Shimadzu semi-preparatory HPLC, using a Phenomenex Luna® semi-preparative phenylhexyl (100 Å, 5 µm, 250 x 10 mm) HPLC column and a gradient of 25-50% acetonitrile: water at 5 mL/min over 20 min.

Compounds **4** and **8** (and **3**) from NascA-DmtD2/E2, PhoN-IPK and NotF-BvnB cultures were isolated by injecting a 50-100 mg/mL solution of the extracts in a Shimadzu semi-preparatory HPLC, using a Phenomenex Lux® cellulose-3 Tris (4-methylbenzoate) semi-preparative (5 μm, 250 x 10 mm) HPLC column, and a step-wise gradient of 15-25% acetonitrile: water over 17 min followed by 30-40% acetonitrile: water over 13 min (5 mL/min).

Fractions containing the desired products were pooled and concentrated to obtain material for NMR characterization.

Characterization of brevianamide F (**1**).

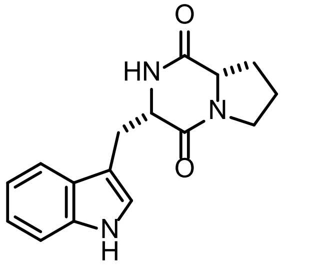

Brevianamide F (**1**)

**HRMS** (ESI-TOF): *m/z* $[M+H]^+$ calculated for $C_{16}H_{17}N_3O_2$ = 284.1394, observed = 284.1428.

**$^1$H NMR** (599 MHz, $CDCl_3$) δ 8.17 (s, 1H), 7.59 (d, *J* = 7.9, 0.9 Hz, 1H), 7.40 (d, *J* = 8.2, 0.9 Hz, 1H), 7.24 (td, 1H), 7.15 (td, *J* = 8.0, 7.0, 1.0 Hz, 1H), 7.13 – 7.12 (m, 1H), 5.72 (s, 1H), 4.38 (dd, *J* = 11.0, 4.2, 1.5 Hz, 1H), 4.11 – 4.05 (m, 1H), 3.77 (ddd, *J* = 15.1, 3.8, 1.1 Hz, 1H), 3.70 – 3.62 (m, 1H), 3.62 – 3.55 (m, 1H), 2.97 (dd, *J* = 15.1, 10.9 Hz, 1H), 2.37 – 2.29 (m, 1H), 2.07 – 1.98 (m, 2H), 1.95 – 1.86 (m, 1H).

**$^{13}$C NMR** (151 MHz, $CDCl_3$) δ 169.47, 165.67, 136.82, 126.86, 123.42, 123.01, 120.22, 118.67, 111.70, 110.21, 59.39, 54.69, 45.57, 28.46, 26.99, 22.79.

Characterization of dehydrobrevianamide F (**2**).

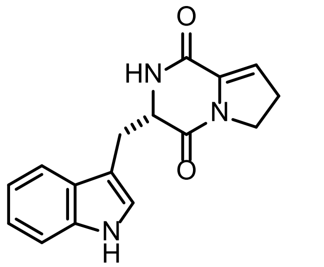

Dehydrobrevianamide F (**2**)

**HRMS** (ESI-TOF): *m/z* $[M+H]^+$ calculated for $C_{16}H_{15}N_3O_2$ = 282.1237, observed = 282.1251.

**$^1$H NMR** 599 MHz, $CDCl_3$) δ 8.16 (s, 1H), 7.64 (d, *J* = 8.0 Hz, 1H), 7.37 (d, *J* = 8.1 Hz, 1H), 7.23 (t, *J* = 7.6 Hz, 1H), 7.15 (t, *J* = 7.5 Hz, 1H), 7.07 (d, *J* = 2.5 Hz, 1H), 5.92 (t, 1H), 5.82 (s, 1H), 4.45 – 4.40 (m, 1H), 3.91 – 3.83 (m, 1H), 3.83 – 3.75 (m, 1H), 3.45 (dd, 1H), 3.28 (dd, *J* = 14.5, 7.8 Hz, 1H), 2.61 – 2.53 (m, 1H), 2.35 – 2.29 (m, 1H).

$^{13}$**C NMR** (151 MHz, $CDCl_3$) δ 162.56, 157.63, 136.56, 132.83, 127.03, 124.48, 122.80, 120.08, 119.07, 118.73, 111.37, 109.13, 57.78, 45.64, 31.77, 27.81.

Characterization of (+)-dehydrodeoxybrevianamide E (**3**).

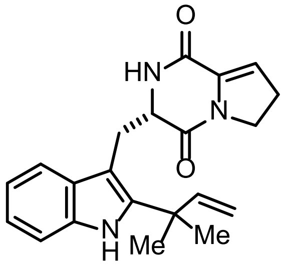

(+)-Dehydrodeoxy
brevianamide E (**3**)

**HRMS** (ESI-TOF): *m/z* $[M+H]^+$ calculated for $C_{21}H_{23}N_3O_2$ = 350.1863, observed = 350.1896.

$[\alpha]_D^{25}$ -34.3° (*c* 0.89, $CHCl_3$).

$^1$**H NMR** (599 MHz, $CDCl_3$) δ 8.05 (s, 1H), 7.53 (d, 1H), 7.32 (d, 1H), 7.17 (ddd, *J* = 8.1, 7.0, 1.2 Hz, 1H), 7.11 (ddd, *J* = 8.0, 7.1, 1.1 Hz, 1H), 6.16 – 6.14 (m, 1H), 6.12 (dd, 1H), 5.67 (s, 1H), 5.20 – 5.15 (m, 2H), 4.52 (dt, *J* = 11.2, 3.2 Hz, 1H), 4.08 (qt, *J* = 12.5, 8.8 Hz, 2H), 3.73 (dd, *J* = 14.7, 3.6 Hz, 1H), 3.23 (dd, *J* = 14.6, 11.3 Hz, 1H), 2.78 (td, *J* = 9.2, 3.1 Hz, 2H), 1.55 (s, 3H), 1.55 (s, 3H).

$^{13}$**C NMR** (151 MHz, $CDCl_3$) δ 162.68, 156.60, 145.78, 141.81, 134.41, 133.23, 128.92, 122.27, 120.26, 119.00, 118.33, 112.60, 110.92, 104.65, 57.57, 45.71, 39.16, 30.92, 28.08, 28.02, 27.92.

Characterization of deoxybrevianamide E (**7**).

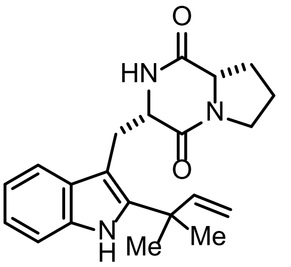

Deoxybrevianamide E (**7**)

**HRMS** (ESI-TOF): *m/z* $[M+H]^+$ calculated for $C_{21}H_{25}N_3O_2$ = 352.202, observed = 352.2052.

$^1$**H NMR** (599 MHz, $CDCl_3$) δ 8.05 (s, 1H), 7.48 (d, *J* = 7.9, 1.0 Hz, 1H), 7.33 (dt, *J* = 8.1, 0.9 Hz, 1H), 7.17 (ddd, *J* = 8.1, 7.1, 1.1 Hz, 1H), 7.11 (ddd, *J* = 8.0, 7.1, 1.0 Hz, 1H), 6.14 (dd, 1H), 5.74 (s, 1H), 5.21 – 5.15 (m, 2H), 4.45 (ddd, *J* = 11.7, 4.3, 1.6 Hz, 1H), 4.10 – 4.04 (m, 1H), 3.75 (dd, *J* = 15.3, 4.1 Hz, 1H), 3.73 – 3.66 (m, 1H), 3.60 (ddd, *J* = 12.0, 8.9, 2.9 Hz, 1H), 3.19 (dd, *J* = 15.3, 11.7 Hz, 1H), 2.39 – 2.31 (m, 1H), 2.11 – 2.02 (m, 2H), 1.98 – 1.86 (m, 1H), 1.56 (d, *J* = 1.1 Hz, 6H).

$^{13}$**C NMR** (151 MHz, $CDCl_3$) δ 169.42, 165.99, 145.73, 141.61, 134.47, 129.22, 122.33, 120.27, 118.02, 112.99, 111.02, 104.73, 59.37, 55.07, 45.54, 39.16, 28.51, 28.10, 28.01, 26.12, 22.79.

Characterization of (-)-dehydrobrevianamide E (**4**).

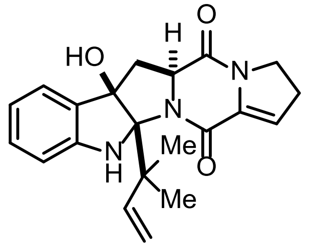

(-)-Dehydrobrevianamide E (**4**)

**HRMS** (ESI-TOF): *m/z* $[M+H]^+$ calculated for $C_{21}H_{23}N_3O_3$ = 366.1812, observed = 366.1841.

$[\boldsymbol{a}]_{\boldsymbol{D}}^{\mathbf{24.8}}$-197.6° (*c* 0.25, EtOH).

**$^1$H NMR** (599 MHz, $CDCl_3$) δ 7.25 (d, *J* = 7.2 Hz, 1H), 7.18 (t, *J* = 7.7 Hz, 1H), 6.82 (t, *J* = 7.4 Hz, 1H), 6.74 (d, *J* = 7.9 Hz, 1H), 6.42 – 6.40 (m, 1H), 6.38 (dd, *J* = 17.6, 10.8 Hz, 1H), 6.16 (t, *J* = 3.1 Hz, 1H), 5.15 (d, *J* = 17.7 Hz, 1H), 5.08 (d, *J* = 10.9 Hz, 1H), 4.10 – 4.03 (m, 1H), 3.92 – 3.84 (m, 1H), 3.82 (dd, *J* = 11.5, 7.3 Hz, 1H), 2.82 – 2.68 (m, 4H), 2.39 (s, 1H), 1.34 (s, 3H), 1.27 (s, 3H).

**$^{13}$C NMR** (151 MHz, $CDCl_3$) δ 163.70, 162.09, 149.39, 144.71, 136.45, 131.00, 129.78, 123.82, 120.63, 120.18, 113.45, 111.24, 91.49, 89.28, 60.33, 45.78, 45.04, 36.76, 29.04, 27.64, 23.10.

Characterization of brevianamide E (**8**).

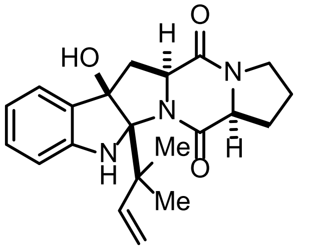

Brevianamide E (**8**)

**HRMS** (ESI-TOF): *m/z* $[M+H]^+$ calculated for $C_{21}H_{25}N_3O_3$ = 368.1969, observed = 368.1999.

**$^1$H NMR** (599 MHz, $CDCl_3$) δ 7.28 (d, *J* = 7.5 Hz, 1H), 7.19 (t, *J* = 7.7, 1.2 Hz, 1H), 6.84 (t, *J* = 7.4 Hz, 1H), 6.74 (d, *J* = 7.9 Hz, 1H), 6.36 – 6.33 (m, 1H), 6.32 (dd, *J* = 17.7, 10.9 Hz, 1H), 5.11 (d, *J* = 17.6, 1.3 Hz, 1H), 5.07 (d, *J* = 10.9, 1.3 Hz, 1H), 3.92 (t, *J* = 8.1 Hz, 1H), 3.73 (dd, *J* = 11.3, 7.4 Hz, 1H), 3.63 – 3.54 (m, 1H), 3.51 (ddd, *J* = 11.7, 8.1, 4.0 Hz, 1H), 2.90 (dd, *J* = 13.1, 11.3 Hz, 1H), 2.65 (dd, *J* = 13.2, 7.4 Hz, 1H), 2.30 (ddd, *J* = 13.0, 6.9, 4.1 Hz, 1H), 2.27 (s, 1H), 2.16 – 2.07 (m, 1H), 1.99 (ddd, *J* = 14.1, 7.3, 3.3 Hz, 1H), 1.94 – 1.85 (m, 1H), 1.28 (s, 3H), 1.25 (s, 3H).

**$^{13}$C NMR** (151 MHz, $CDCl_3$) δ 172.74, 165.95, 149.14, 144.59, 130.90, 130.39, 123.84, 120.25, 113.26, 111.55, 91.21, 89.42, 61.93, 59.83, 45.14, 44.98, 35.40, 28.55, 26.87, 23.86, 23.36.

**Isolation of (+)-Brevianamides A and B from native organism *Penicillium brevicompactum* (adapted from Ying *et al.*)[6]**

*P. brevicompactum* was cultured on potato dextrose agar (PDA) plates at 28 °C for seven days from a glycerol stock. Five (5) L of solid Czapek–Dox agar (CDA) media were inoculated with the spore suspension and incubated at 28 °C for six days. The resulting mold was incubated with ethyl acetate: MeOH 85:15 before being liquid-liquid extraction with water: ethyl acetate. The resulting supernatant was incorporated into silica through vacuum filtration. The silica mixture was packed into a cartridge that was attached to a silica column and subjected to normal-phase flash chromatography using an isocratic method of 20% hexanes and 80% 1:1 ethyl acetate:acetone for 13-17 column volumes on a Biotage.

Further purification was conducted by semi-preparative HPLC using a Phenomenex Lux® cellulose-3 Tris (4-methylbenzoate) semi-preparative (5 µm, 250 x 10 mm) HPLC column and a gradient of 40-52% acetonitrile: water at 5 mL/min over 14 min. Fractions containing the desired products were pooled and concentrated for NMR characterization.

Characterization of (+)-brevianamide A (**5**) isolated from native fungi.

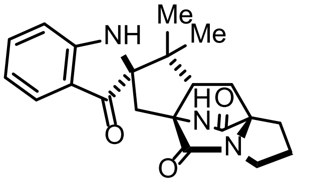

(+)-Brevianamide A (**5**)

**HRMS** (ESI-TOF): *m/z* $[M+H]^+$ calculated for $C_{21}H_{23}N_3O_3$ = 366.1812, observed = 366.1823.

**$^1$H NMR** (599 MHz, $CDCl_3$) δ 7.56 (d, 1H), 7.45 (ddd, *J* = 8.4, 7.1, 1.4 Hz, 1H), 6.85 – 6.78 (m, 2H), 6.49 (s, 1H), 4.93 (s, 1H), 3.52 – 3.42 (m, 2H), 2.81 – 2.74 (m, 2H), 2.39 (dd, 1H), 2.32 (d, *J* = 15.6 Hz, 1H), 2.09 – 1.99 (m, 2H), 1.96 – 1.80 (m, 3H), 1.12 (s, 3H), 0.93 (s, 3H).

**$^{13}$C NMR** (151 MHz, $CDCl_3$) δ 202.20, 172.42, 169.72, 160.15, 137.79, 124.83, 121.14, 119.40, 112.11, 78.81, 69.53, 67.84, 55.73, 48.39, 44.05, 37.48, 29.16, 28.98, 25.09, 24.22, 20.00.

Characterization of (+)-brevianamide B (**6**) isolated from native fungi.

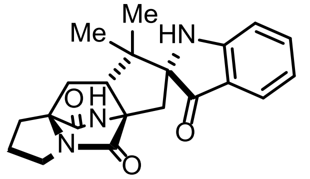

(+)-Brevianamide B (**6**)

**HRMS** (ESI-TOF): *m/z* $[M+H]^+$ calculated for $C_{21}H_{23}N_3O_3$ = 366.1812, observed = 366.1814.

**$^1$H NMR** (800 MHz, DMSO) δ 8.67 (s, 1H), 7.42 (ddd, *J* = 8.3, 7.0, 1.4 Hz, 1H), 7.34 (d, *J* = 7.7 Hz, 1H), 7.30 (s, 1H), 6.90 (d, *J* = 8.2 Hz, 1H), 6.65 (t, *J* = 7.3 Hz, 1H), 3.30 – 3.21 (m, 2H), 3.01 (dd, *J* = 10.3, 7.5 Hz, 1H), 2.67 (d, *J* = 15.2 Hz, 1H), 2.46 (dd, *J* = 12.3, 6.2 Hz, 1H), 2.00 – 1.91 (m, 3H), 1.82 – 1.76 (m, 2H), 1.64 (dd, *J* = 13.0, 7.5 Hz, 1H), 1.05 (s, 3H), 0.63 (s, 3H).

**$^{13}$C NMR** (201 MHz, DMSO) δ 205.24, 173.17, 161.79, 137.48, 124.32, 118.66, 117.51, 111.90, 78.03, 68.67, 65.78, 49.34, 46.52, 43.75, 33.97, 28.88, 28.04, 24.89, 22.40, 20.25.

**$^{1}$H NMR** (599 MHz, $CDCl_3$) δ 7.56 (d, *J* = 7.7, 4.4 Hz, 1H), 7.43 (t, *J* = 7.5 Hz, 1H), 6.84 – 6.77 (m, 2H), 5.96 (s, 1H), 4.78 (s, 1H), 3.51 – 3.45 (m, 2H), 3.30 (dd, 1H), 3.26 (d, *J* = 15.6 Hz, 1H), 2.75 (ddd, *J* = 26.1, 14.3, 6.7 Hz, 1H), 2.13 – 1.90 (m, 3H), 1.90 – 1.76 (m, 2H), 1.72 (d, *J* = 15.6 Hz, 1H), 1.12 (s, 3H), 0.83 (s, 3H).

**Lithium hydroxide reaction of (-)dehydrobrevianamide E (4).**
The lithium hydroxide reaction was conducted on **4** as reported previously.[4] The dried material was later resuspended in MeOH to a concentration of 1.5 mg/mL and injected on a Shimadzu HPLC with a Phenomenex Lux® cellulose-3 Tris (4-methylbenzoate) LC column (100 Å, 5 μm, 250 x 4.6 mm) using a linear gradient of 25-60% acetonitrile: water over 12.5 min (1 mL/min) and a Phenomenex Lux® cellulose-3 Tris (4-methylbenzoate) semi-preparative (5 μm, 250 x 10 mm) HPLC column and a isocratic method of 20% acetonitrile: water over 10 min (5 mL/min).

Characterization of (+)-brevianamide A (**5**) isolated from LiOH reaction.

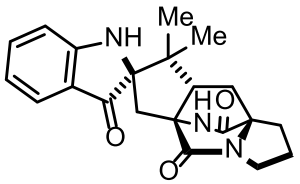

(+)-Brevianamide A (**5**)

**Rf** 0.26 (3:7 THF/$CHCl_3$).

**HRMS** (ESI-TOF): *m/z* $[M+H]^+$ calculated for $C_{21}H_{23}N_3O_3$ = 366.1812, observed = 366.1816.

$[\alpha]_D^{24.8}$ +392° (*c*, EtOH).

**$^{1}$H NMR** (599 MHz, $CDCl_3$) δ 7.57 (d, *J* = 7.7 Hz, 1H), 7.45 (t, *J* = 7.5 Hz, 1H), 6.86 – 6.79 (m, 2H), 6.46 (s, 1H), 4.92 (s, 1H), 3.50 – 3.43 (m, 2H), 2.81 – 2.73 (m, 2H), 2.39 (dd, *J* = 9.8, 7.2 Hz, 1H), 2.31 (d, *J* = 15.6 Hz, 1H), 2.09 – 1.99 (m, 2H), 1.93 (dd, *J* = 13.3, 9.9 Hz, 1H), 1.90 – 1.81 (m, 2H), 1.12 (s, 3H), 0.93 (s, 3H).

**$^{13}$C NMR** (151 MHz, $CDCl_3$) δ 202.23, 172.39, 169.72, 160.16, 137.79, 124.82, 121.13, 119.39, 112.11, 78.80, 69.53, 67.83, 55.73, 48.39, 44.04, 37.48, 29.16, 28.99, 25.08, 24.23, 20.01.

Characterization of (+)-brevianamide B (**6**) isolated from native fungi.

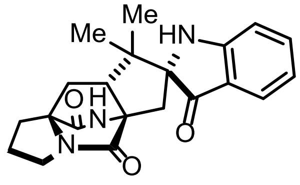

(+)-Brevianamide B (**6**)

**Rf** 0.09 (3:7 THF/$CHCl_3$).

**HRMS (**ESI-TOF): *m/z* $[M+H]^+$ calculated for $C_{21}H_{23}N_3O_3$ = 366.1812, observed = 366.1815.

**$^1$H NMR** (599 MHz, $CDCl_3$) δ 7.55 (d, *J* = 7.5 Hz, 1H), 7.43 (ddd, *J* = 8.0 Hz, 1H), 6.85 – 6.75 (m, 2H), 5.94 (s, 1H), 3.48 (t, *J* = 6.8 Hz, 2H), 3.33 – 3.28 (m, 1H), 3.26 (dd, *J* = 15.9, 6.6 Hz, 1H), 2.73 (dt, *J* = 12.7, 6.4 Hz, 1H), 2.07 – 1.94 (m, 3H), 1.89 – 1.75 (m, 2H), 1.77 – 1.69 (m, 1H), 1.13 (s, 3H), 0.83 (s, 3H).

**Supplementary Figure 9.** Individual HPLC traces of *in vivo* engineered pathway intermediates with standards.

**a)** HPLC traces of NascA and DmtD2/E2 enzymatic products. (i) **1** standard (ii) **2** standard (iii) Expression of NascA and DmtD2/E2.

**b)** HPLC traces of NotF enzymatic products. (i) **7** standard (ii) **3** standard (iii) Expression of NascA, DmtD2/E2, PhoN, IPK, and NotF.

**c)** HPLC traces of NotF and BvnB products (i) **4** standard (ii) **8** standard (iii) **7** standard (vi) Expression of NascA, DmtD2/E2, PhoN, IPK, NotF, and BvnB.

Separation between peaks in **a)** and **c)** was achieved using a cellulose column while separation of peaks in **b)** could only be accomplished using a phenylhexyl column.

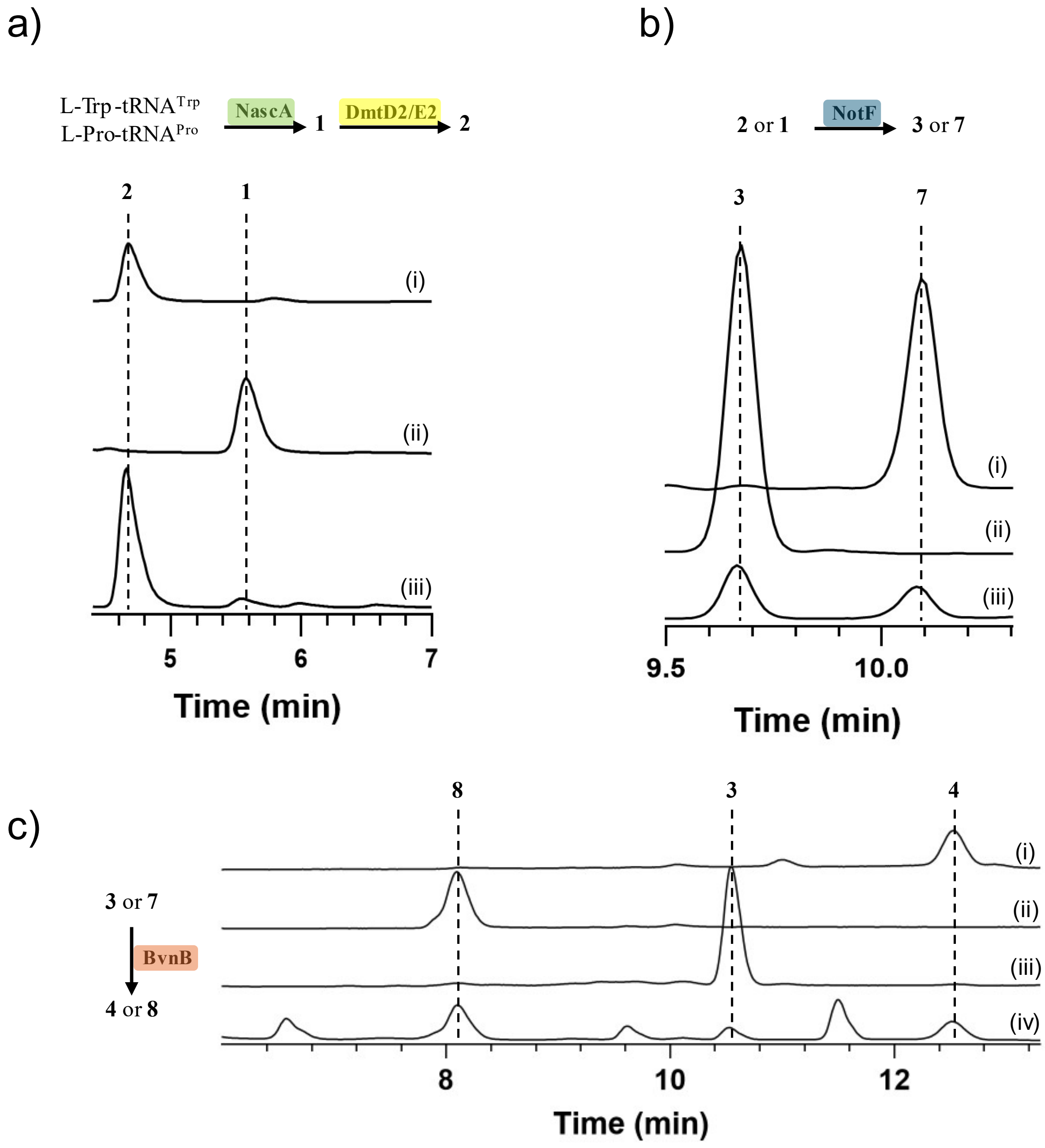

**Supplementary Figure 10.** HPLC traces of *in vivo* and *in vitro* reactions using unoptimized BvnB conditions at 2 days incubation at 30 °C with a phenylhexyl column. (i) **7** standard (ii) Expression of NascA and DmtD2/E2 (iii) Expression of NascA, DmtD2/E2, PhoN, IPK, and NotF (iv) Expression of NascA, DmtD2/E2, PhoN, IPK, NotF, and BvnB (v) no enzyme control, incubation with **2** (vi) *in vitro* reaction of NotF with **2** (vii) *in vitro* reaction of NotF with **2** and BvnB with **3**.

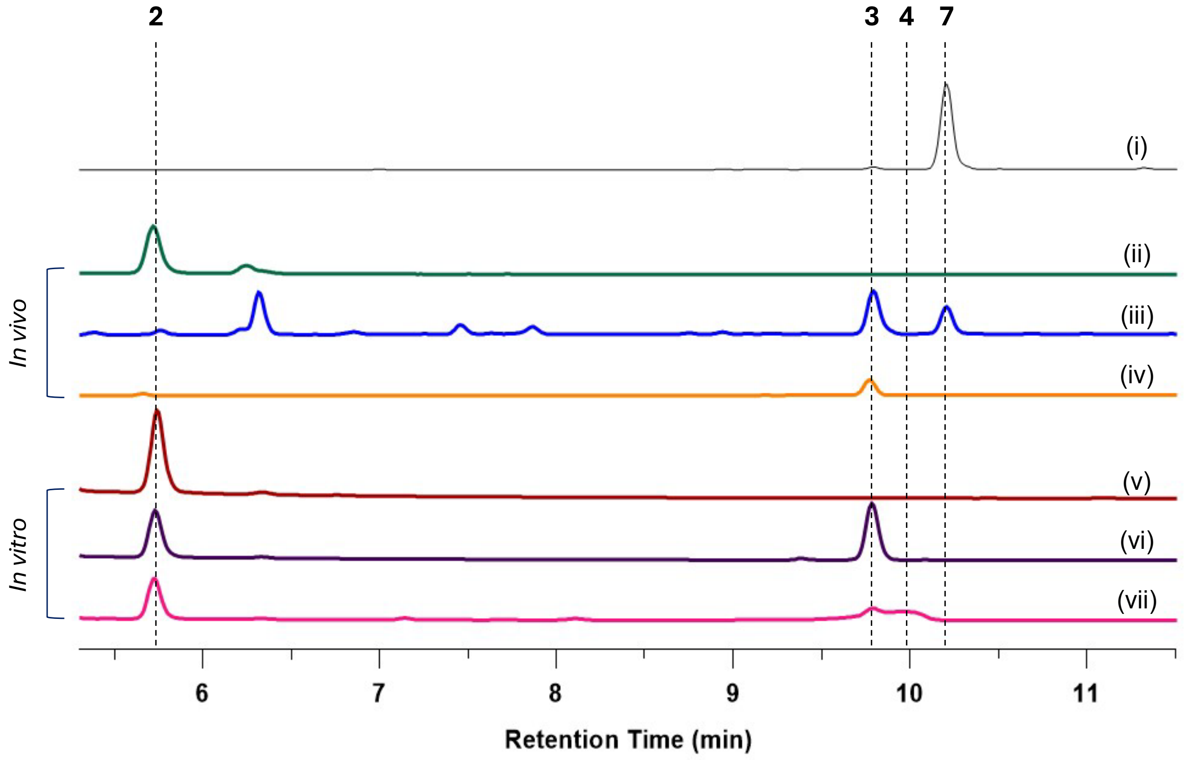

**Supplementary Figure 11.** Temperature optimization of *in vivo* (-)-dehydrobrevianamide E (**4**) production with NascA-DmtD2/E2, NotF-BvnB, and PhoN/IPK co-transformant. (i) **4** standard (ii) Expression of NascA, DmtD2/E2, PhoN, IPK, NotF, and BvnB at 30 °C, (iii) Expression of NascA, DmtD2/E2, PhoN, IPK, NotF, and BvnB at 18 °C (iv) *in vitro* reaction of **2** with NotF-BvnB.

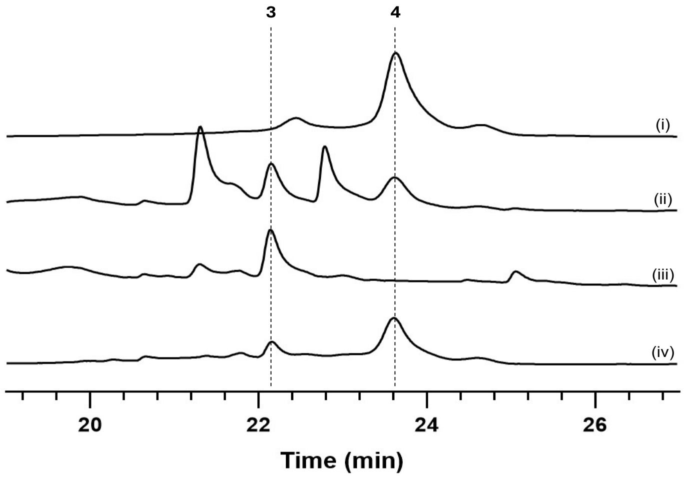

**Supplementary Figure 12.** Incubation time optimization of *in vivo* (-)-dehydrobrevianamide E (**4**) production with NascA-DmtD2/E2, NotF-BvnB, and PhoN-IPK co-transformant.

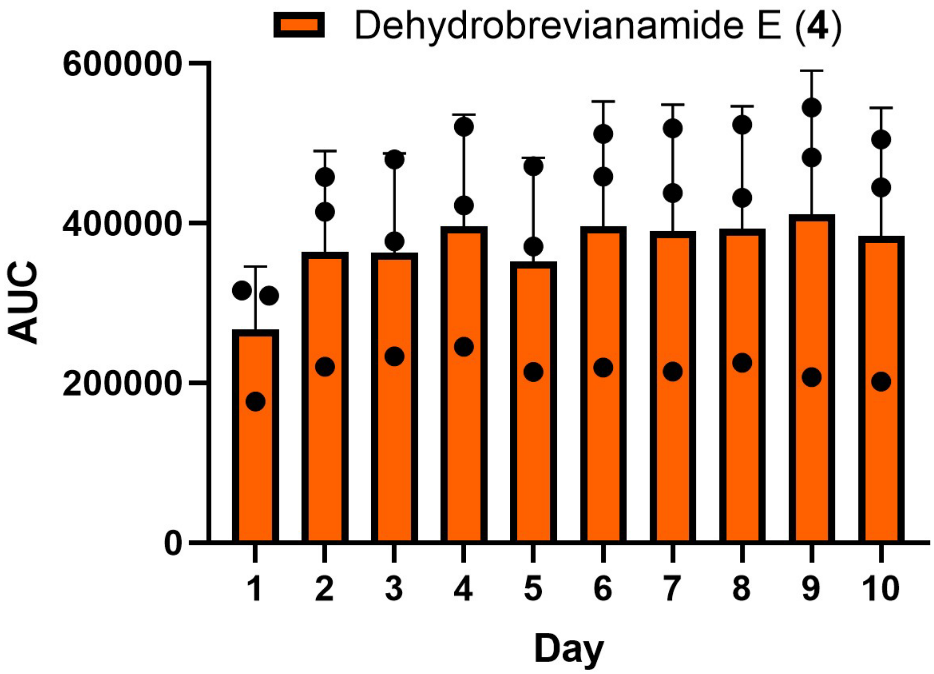

**Supplementary Figure 13.** Percent conversion results.

**a)** BvnB reaction of **7** to **8**. HPLC traces of (i) **7** standard (ii) No enzyme control (iii) Reaction of BvnB with **7** replicate-1 (iv) Reaction of BvnB with **7** replicate-2 (v) Reaction of BvnB with **7** replicate-3 (vi) MeOH blank.

**b)** BvnB reaction of **3** to **4**. HPLC traces of (i) **3** standard (ii) No enzyme control (iii) Reaction of BvnB with **3** replicate-1 (iv) Reaction of BvnB with **3** replicate-2 (v) Reaction of BvnB with **3** replicate-3 (vi) MeOH blank.

**c)** In tandem BvnB reaction of **7** and **3** to **8** and **4**. HPLC traces of (i) **3** standard (ii) **7** standard (iii) No enzyme control (iv) Reaction of BvnB with **3** and **7** replicate-1 (v) Reaction of BvnB with **3** and **7** replicate-2 (vi) Reaction of BvnB with **3** and **7** replicate-3 (vii) MeOH blank.

**d)** Calibration curves of compound **3** and **7** which were used to calculate percent conversions which are listed in tables.

a)

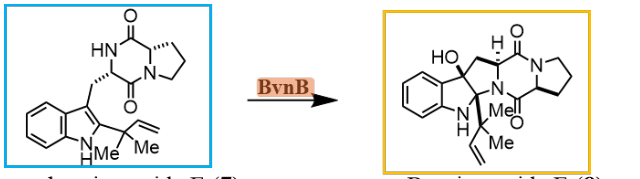

Deoxybrevianamide E (**7**)

Brevianamide E (**8**)
87.3% ± 5.1 % conversion

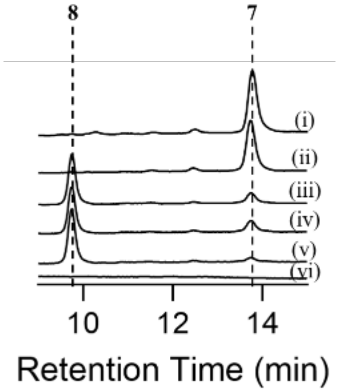

| Name | RT (min) | AUC | Concentration (µM) | % Conversion | Average (%) | Standard Deviation (%) |
|---|---|---|---|---|---|---|
| DBE 1 | 13.756 | 23422 | 48.9 | 84.8 | **87.3** | 5.1 |
| DBE 2 | 13.745 | 26818 | 55.9 | 82.6 | | |
| DBE 3 | 13.738 | 8462 | 17.9 | 94.4 | | |
| DBE ctrl | 13.734 | 155021 | 321.5 | | | |

b)

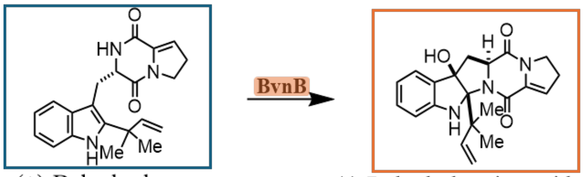

(+)-Dehydrodeoxy brevianamide E (**3**)

(-)-Dehydrobrevianamide E (**4**)
83.3% ± 1.1% conversion

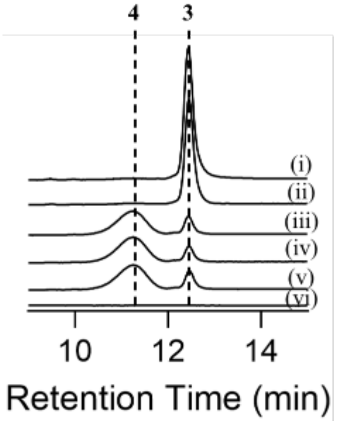

| Name | RT (min) | AUC | Concentration (µM) | % Conversion | Average (%) | Standard Deviation (%) |
|---|---|---|---|---|---|---|
| dDBE 1 | 12.438 | 90439 | 50.7 | 82.9 | **83.3** | 1.1 |
| dDBE 2 | 12.452 | 80334 | 45.1 | 84.8 | | |
| dDBE 3 | 12.456 | 95238 | 53.3 | 82.0 | | |
| dDBE ctrl | 12.448 | 538674 | 296.9 | | | |

c)

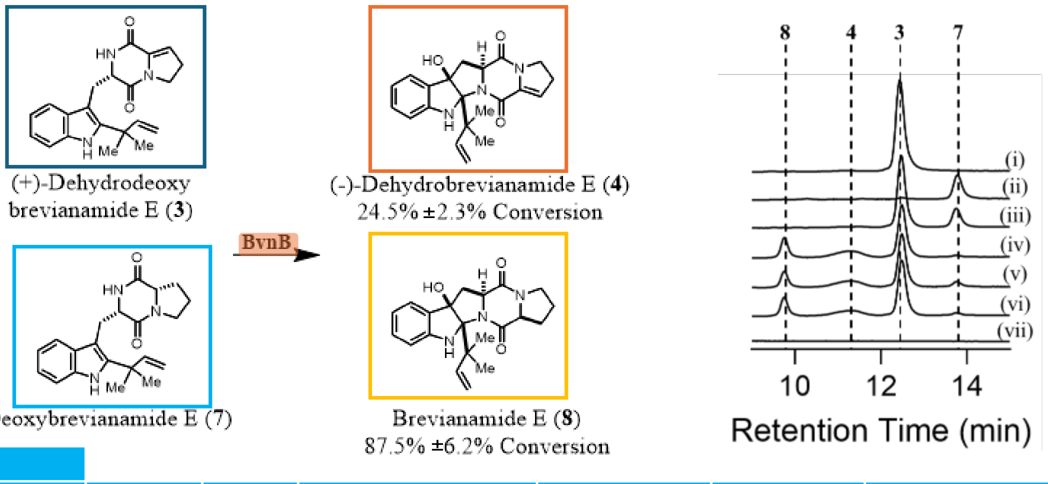

| DBE | | | | | | |
|---|---|---|---|---|---|---|
| **Name** | **RT (min)** | **AUC** | **Concentration (µM)** | **% Conversion** | **Average (%)** | **Standard Deviation (%)** |
| dDBE-DBE 1 | 13.756 | 9283 | 19.6 | 93.7 | **87.5** | 6.2 |
| dDBE-DBE 2 | 13.763 | 31348 | 65.3 | 79.1 | | |
| dDBE-DBE 3 | 13.752 | 15144 | 31.8 | 89.8 | | |
| dDBE-DBE ctrl | 13.748 | 150538 | 312.2 | | | |
| **dDBE** | | | | | | |
| **Name** | **RT (min)** | **AUC** | **Concentration (µM)** | **% Conversion** | **Average (%)** | **Standard Deviation (%)** |
| dDBE-DBE 1 | 12.483 | 388657 | 214.5 | 26.5 | **24.5** | 2.3 |
| dDBE-DBE 2 | 12.483 | 393109 | 216.9 | 25.7 | | |
| dDBE-DBE 3 | 12.481 | 416352 | 229.7 | 21.3 | | |
| dDBE-DBE ctrl | 12.469 | 529729 | 291.9 | | | |

d)

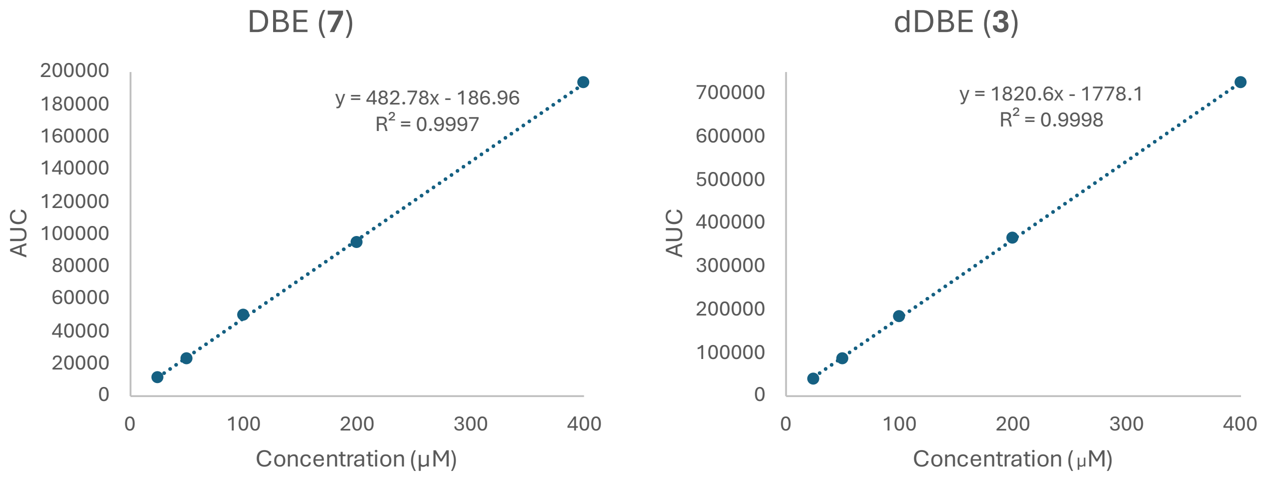

**BvnB kinetic assay conditions for reactions with 3 and 7**

Reactions for kinetics assays were assembled on a Corning® 96-well Clear Flat Bottom Polystyrene Microplates (product number 3596) using 10 µM FAD, 500 µM MgCl2, varying concentrations of substrate, 1 mM (experiments with **3**) or 25 µM (experiments with **7**) BvnB, and 400 µM NADPH in reaction buffer (5% v/v glycerol, 300 mM NaCl, 20 mM Tris pH 7.9) to an end volume of 100 µL. The concentration series of **3** is as follows: 6 mM, 3 mM, 1 mM, 0.5 mM. The concentration of **7** is as follows:

250 mM, 175 mM, 100 mM, 75 mM, 50 mM, 37.5 mM, 25 mM, 18.75 mM, 12.5 mM, 6.25 mM. NADPH was added to the reactions and the plate was immediately placed in a SpectraMax M5 spectrophotometer where it was shaken for 15 seconds before the first read. 340 nm absorbance was measured over 20 (experiments with **3**) or 15 (experiments with **7**) minutes at 15 (experiments with **3**) or 10 (experiments with **7**) second intervals at room temperature. The differences in absorbances of no substrate controls over time were used to normalize data and account for the spontaneous oxidation of NADPH. The pathlength of NADPH was ascertained to be 0.25 cm. Concentration of product was determined by calculating concentration of NADPH consumed and plotted against time. The velocities of each reaction were plotted against substrate concentration to construct Michaelis-Menten curves. The $K_m$ and $V_{max}$ values were obtained using nonlinear regression analysis in GraphPad Prism software.

**Supplementary Figure 14.** Kinetic Data from BvnB reactions with **3** and **7**.

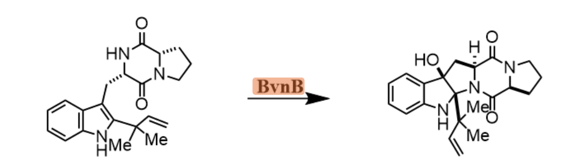

Deoxybrevianamide E (**7**) → Brevianamide E (**8**)

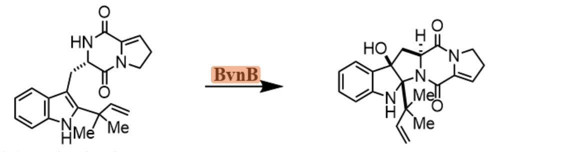

(+)-Dehydrodeoxy brevianamide E (**3**) → (-)-Dehydrobrevianamide E (**4**)

$K_m = 5.5 \times 10^{-4}$ M
$V_{max} = 2.1 \times 10^{-6}$ M/s
$k_{cat} = 8.2 \times 10^{-2}$ $s^{-1}$
$k_{cat}/K_m = 149.3$ $M^{-1}$ $s^{-1}$
N = 2

$K_m = 3.4 \times 10^{-2}$ M
$V_{max} = 2.1 \times 10^{-5}$ M/s
$k_{cat} = 2.1 \times 10^{-2}$ $s^{-1}$
$k_{cat}/K_m = 0.6$ $M^{-1}$ $s^{-1}$
N = 3

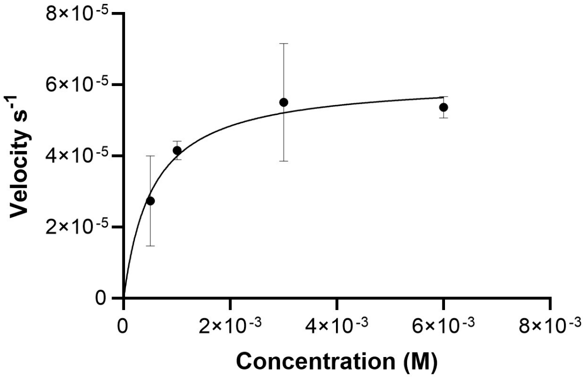

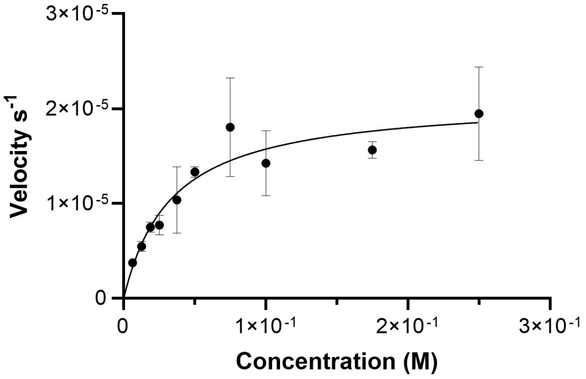

**Supplementary Figure 15.** Concentration-dependent NADPH effect on BvnB activity.

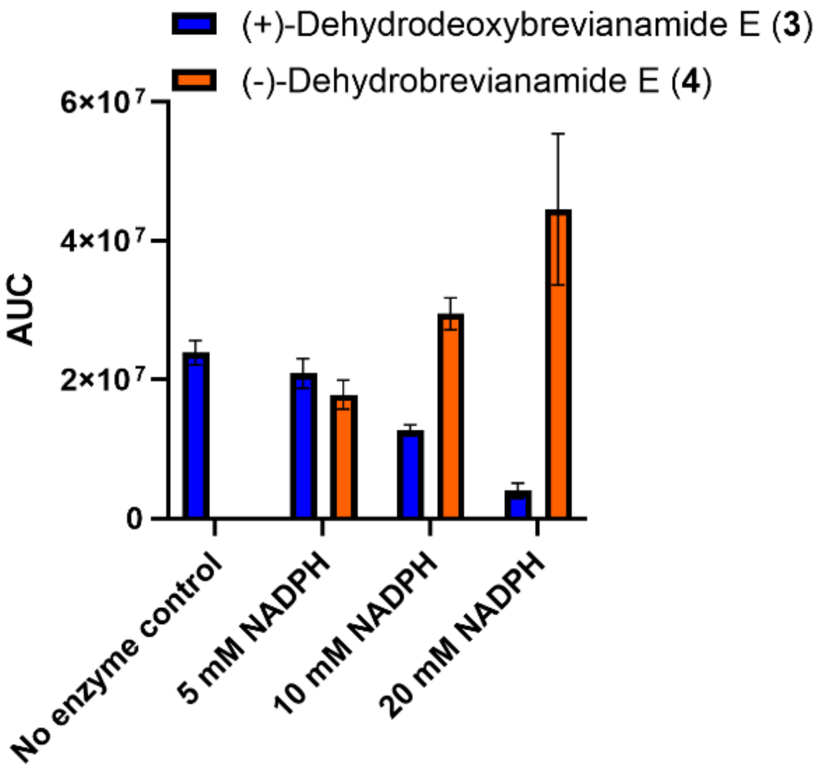

#### ***pfkA*** **CRISPR-Cas9 knock-out**

Gene editing was accomplished using modifications to the protocol provided by Li *et al.*[7] The components for gene editing consisted of a plasmid containing Cas9, *pEcCas* (Addgene #73227), a plasmid containing sgRNA, *pEcgRNA* (Addgene #166581), and a 1000 bp HDR (Homologous Directed Repair) template. Single stranded sgRNA oligos were chosen via webtool CHOPCHOP, annealed, and then inserted into linearized *pEcgRNA* as previously described.[7,8] The HDR template was constructed through overlap extension PCR where two amplified fragments upstream and downstream of *pfkA* with a small overlap region were combined into one fragment. Next, we transformed *pEcCas* into chemically competent BL21(DE3) cells. The next day, we fermented a single colony in LB media with kanamycin and induced the expression of λ-red genes. We harvested the cells, washed with $CaCl_2$, and resuspended with 100 mM $CaCl_2$. These newly made chemically competent cells were co-transformed with *pEcgRNA* containing the *pfkA* sgRNA and the *pfkA* HDR template and plated onto LB-agar containing 50 μg/mL kanamycin and 50 μg/mL spectinomycin. We performed colony PCR with individual colonies and observed the expected size (~1600bp). These colonies were suspended in LB media containing 50 μg/mL kanamycin and 10 mM rhamnose to remove the *pEcgRNA* plasmid and thereafter plated on LB agar with 50 μg/mL kanamycin. Individual colonies were plated on LB agar containing 50 μg/mL kanamycin and 50 μg/mL spectinomycin to determine *pEcgRNA* curing. Successfully cured colonies were resuspended in LB and 5 g/L glucose and thereafter plated on LB agar with 5 g/L glucose and 10 g/L sucrose to remove the *pEcCas* plasmid. Finally, we transformed *pRARE*, *pETDuet-1-phoN-ipk*, *pRSFDuet-1-nascA-dmtd2/e2*, and *pACYCDuet-1-notF-bvnB* into the *ΔpfkA* strain in a step-wise fashion and incubated for one to four days before quenching the reactions with three times the culture amount with methanol and injected these samples into LC-MS to compare the production of **4** in mutant vs the wild-type using AUCs.

**_pfkA_ HDR template**

CTATATTTTATATAGCGCGTTACGCATGGGATATGAGGCGGTACAGTCATTACTGGATCGCGCATTGCCT
GATGAGGAACGGCAAGAAATTATTGATATCGTGACTTCCTGGCCGGGTGTTAGCGGCGCTCACGATCTTC
GCACGCGGCAGTCAGGGCCGACCCGCTTTATTCAGATTCATTTGGAAATGGAAGACTCTCTGCCTTTGGT
TCAGGCACATATGGTGGCGGATCAGGTAGAGCAGGCTATTTTACGGCGTTTTCCGGGATCGGATGTAATT
ATCCATCAGGACCCCTGTTCCGTCGTACCCAGGGAGGGTAAACGGTCTATGCTTTCATAATCAGTATAAA
AGAGAGCCAGACCCGCATTTTGTGTATAAAATACCGCCATTTGGCCTGACCTGAATCAATTCAGCAGGAA
GTGATTGTTATACTATTTGCACATTCGTTGGATCACTTCGATGTGCAAGAAGACTTCCGGCAACAGATTT
CATTTTGCATTCCAAAGTTCAGAGGTAGTCATG
TAATGATTTCGGAAAAAGGCAGATTCCTTTACCCTGAAACCGATGACAGAAGCAAAAATGCCTGATGCGC
TTCGCTTATCAGGCCTACATGAATTCTGCAATTTATTGAATTTGCAAACTTTTGTAGGCCGGATAAGGCG
TTCGCGCCGCATCCGGCATGGACAAAGCGCACTTTGTCAGCAATATGAGGCGGATTTCTTCCGCCTTTTT
AATCCCTCAACATATACCCGCAAGTTATAGCCAATCTTTTTTTATTCTTTAATGTTTGGTTAACCTTCTG
GCACGCTTTGCTCATCACAACACAACATAAGAGAGTCGGGCGATGAACAAGTGGGGCGTAGGGTTAACAT
TTTTGCTGGCGGCAACCAGCGTTATGGCAAAGGATATTCAGCTTCTTAACGTTTCATATGATCCAACGCG
CGAATTGTACGAACAGTACAACAAGGCATTCAGCGCCCACTGGAAACAGCAAACTGGTGATAACGTGGTG
ATTCGTCAGTCACACGGTGGCTCAGGTAAA

**Colony PCR product sequencing data for _pfka_ KO**

attctgctggcgctggggttgtcctggtacggctggcatcgcgccgatgctctgtttgcattgggaatcggcatctatattttatatagcgcgttacgc
atgggatatgaggcggtacagtcattactggatcgcgcattacctgatgaggaacggcaagaaattattgatatcgtgacttcctggccgggtgt
tagcggcgctcacgatcttcgcacgcggcagtcagggccgacccgctttattcagattcatttggaaatggaagactctctgcctttggttcaggc
acatatggtggcggatcaggtagagcaggctattttacggcgttttccgggatcggatgtgattatccatcaggacccctgttccgtcgtacccag
ggagggtaaacggtctatgctttcataattagtataaaagagagccagacccgcattttgtgtataaaataccgccatttggcctgacctgaatc
aattcagcaggaagtgattgttatactatttgcacattcgttggatcacttcgatgtgcaagaagacttccggcaacagatttcattttgcattccaa
agttcagaggtagtcatgtaatgatttcggaaaaaggcagattcctttagcctgaaaccgatgacagaagcaaaaatgcctgatgcgcttcgct
tatcaggcctacgtgaattctgcaatttattgaatttacaaatttttgtaggtcggataaggcgttcgcgccgcatccggcatcgataaagcgcactt
tgtcagcaatatgaggcggatttcttccgcctttttaattcctcaacatatacccgcaagttatagccaatctttttttattctttaatgtttggttaaccttct
ggcacgctttgctcatcacaacacaacataagagagtcgggcgatgaacaagtggggcgtagggttaacatttttgctggcggcaaccagcg
ttatggcaaaggatattcagcttcttaacgtttcatatgatccaacgcgcgaattgtacgaacagtacaacaaggcattcagcgcccactggaa
acagcaaactggcgataacgtggtgatccgtcagtcccacggtggttcaggcaaacaagcgacgtcggtaatcaacggtattgaagctgatg
ttgtcacgctggctctggcctatgacgtggacgcaattgcggaacgcgggcggattgataaagagtggatcaaacgtctgccggataactccg
caccgtacacttccaccattgttttcctggtgcgtaagggcaatccgaagcagatccatgactggaacgatctgattaaaccgggtgtttcggtg
atcacgcctaatccgaaaagctctggtggcgcacgctggaactacctggctgcctggggctacgcgctgcatcacaacaacaacgatcagg
caaaagcacaggattttgttcgggcactgtataaaaacgtcgaagttctggattctggcgcgcgcggctccactaacacttttgtcgagcgcgg
aattggcgatgtactgatcgcctgggaaaacgaagctctgttggcagcgaatgaactggggaaagataaattcgaaatcgtc

**Supplementary Figure 16.** DNA gel of colony PCR product for KO of *pfkA*. A) DNA ladder, B) colony PCR product. Expected size ~ 1600bp.

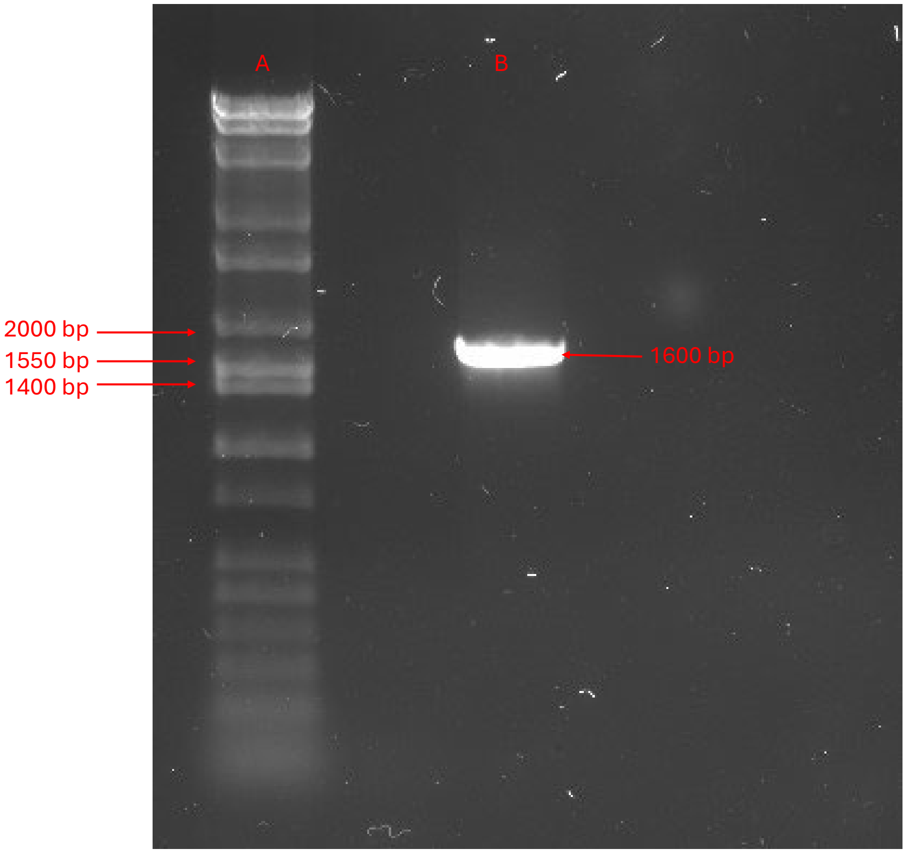

**Supplementary Figure 17.** Fold differences of **4** (a), **8** (b), and **3** (c) in different conditions relative to original conditions (**4** pathway in glycerol) after **a single day** of incubation. Tables list gly and glu as abbreviations for glycerol and glucose. "gly" refers to the original conditions.

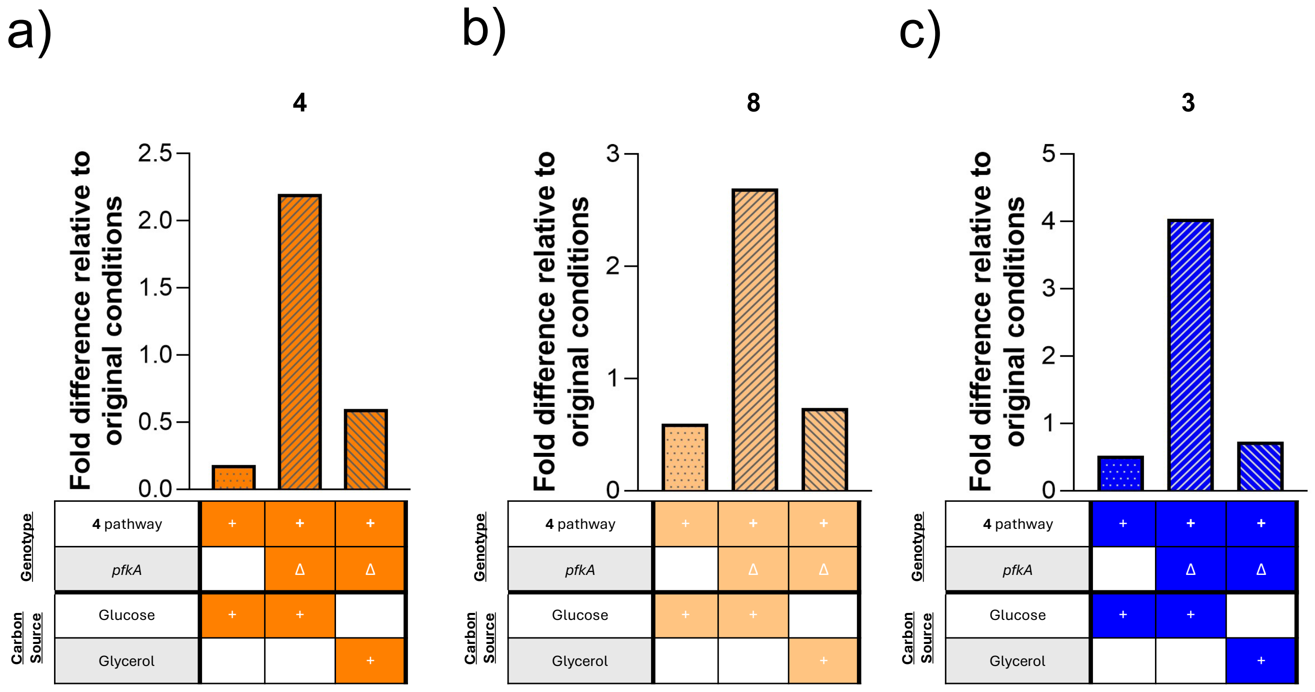

| | ΔpfkA glu | ΔpfkA gly | glu | gly |
|---|---|---|---|---|
| Mean AUC | 437971.1 | 118885.1 | 36460.35 | 199272.5 |
| Standard Deviation | 34470.66 | 28550.75 | 11945.7 | 73761.77 |
| # of samples | 3 | 3 | 3 | 3 |

| | ΔpfkA glu | ΔpfkA gly | glu | gly |
|---|---|---|---|---|
| Mean AUC | 7413372 | 2034764 | 1646376 | 2755855 |
| Standard Deviation | 929048.4 | 1536862 | 338321.5 | 777183.3 |
| # of samples | 3 | 3 | 3 | 3 |

| | ΔpfkA glu | ΔpfkA gly | glu | gly |
|---|---|---|---|---|
| Mean AUC | 3419305 | 617064 | 440574.4 | 846618.2 |
| Standard Deviation | 256853.3 | 120857.7 | 132774.6 | 261793.8 |
| # of samples | 3 | 3 | 3 | 3 |

**Supplementary Figure 18.** Fold differences of **4** (a), **8** (b), and **3** (c) in different conditions relative to original conditions (**4** pathway in glycerol) after **four days** of incubation. Tables list gly and glu as abbreviations for glycerol and glucose. “gly” refers to the original conditions.

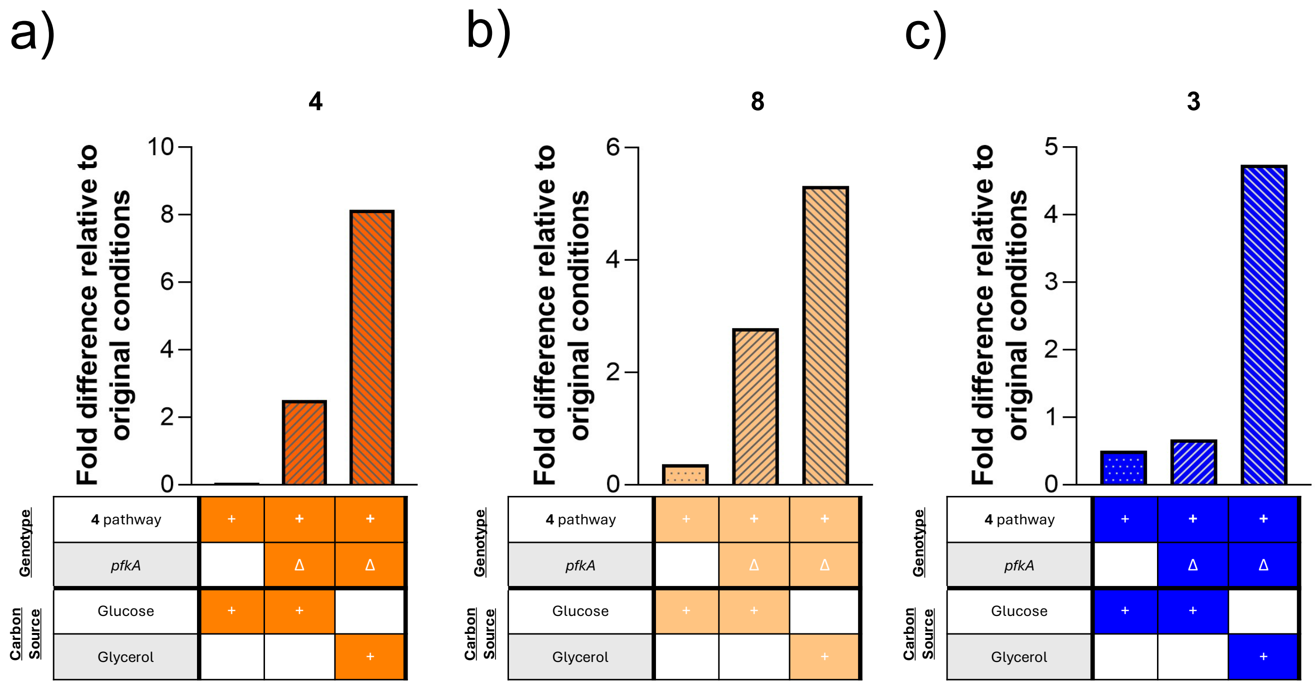

| | ΔpfkA glu | ΔpfkA gly | glu | gly |
|---|---|---|---|---|
| Mean AUC | 5276263 | 17171117 | 130476.1 | 2108088 |
| Standard Deviation | 344876 | 3091507 | 97722.4 | 1289861 |
| # of samples | 3 | 3 | 3 | 3 |

| | ΔpfkA glu | ΔpfkA gly | glu | gly |
|---|---|---|---|---|
| Mean AUC | 15497973 | 29533514 | 2034067 | 5560387 |
| Standard Deviation | 2600475 | 3246197 | 1576082 | 3278112 |
| # of samples | 3 | 3 | 3 | 3 |

| | ΔpfkA glu | ΔpfkA gly | glu | gly |
|---|---|---|---|---|
| Mean AUC | 874653.3 | 6174394 | 658857.2 | 1302223 |
| Standard Deviation | 124350.1 | 1404110 | 555477.3 | 1162394 |
| # of samples | 3 | 3 | 3 | 3 |

**Supplementary Table 3.** Titers (mg/L) of (+)-brevianamides A (**5**) and B (**6**) using native pathway and LiOH reaction

| | (+)-Brevianamide A (**5**) | (+)-Brevianamide B (**6**) |
|---|---|---|
| Native *P. brevicompactum* | 2.02 | 0.08 |
| LiOH rxn from 5.3 mg/L titers of **4** | 2.37 | 0.207 |

**Supplementary Figure 19.** Mass spectrum (TOF LC/MS) of **3** from NotF *in vitro* cascade supplemented with **2**.

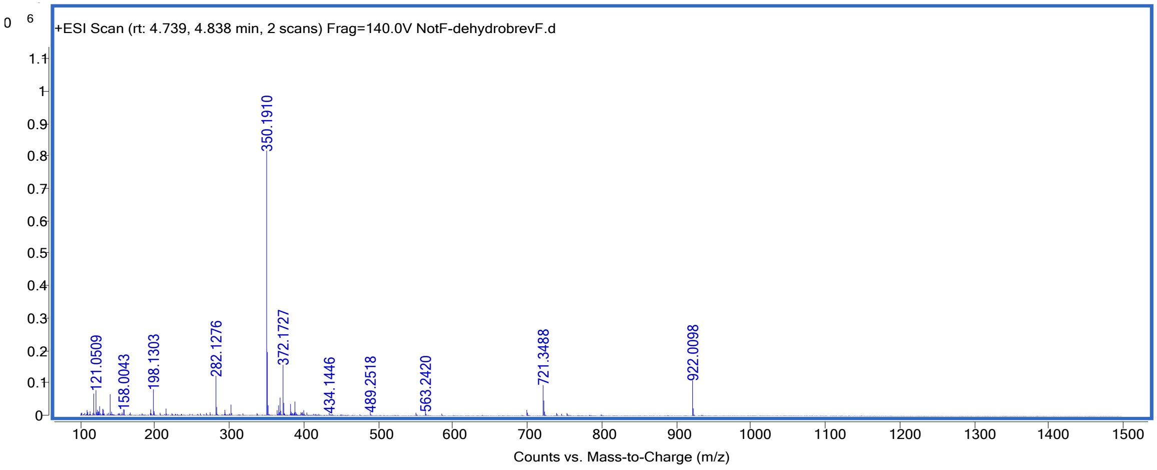

**Supplementary Figure 20.** Mass spectrum (TOF LC/MS) of **4** from NotF-BvnB *in vitro* cascade supplemented with **2**.

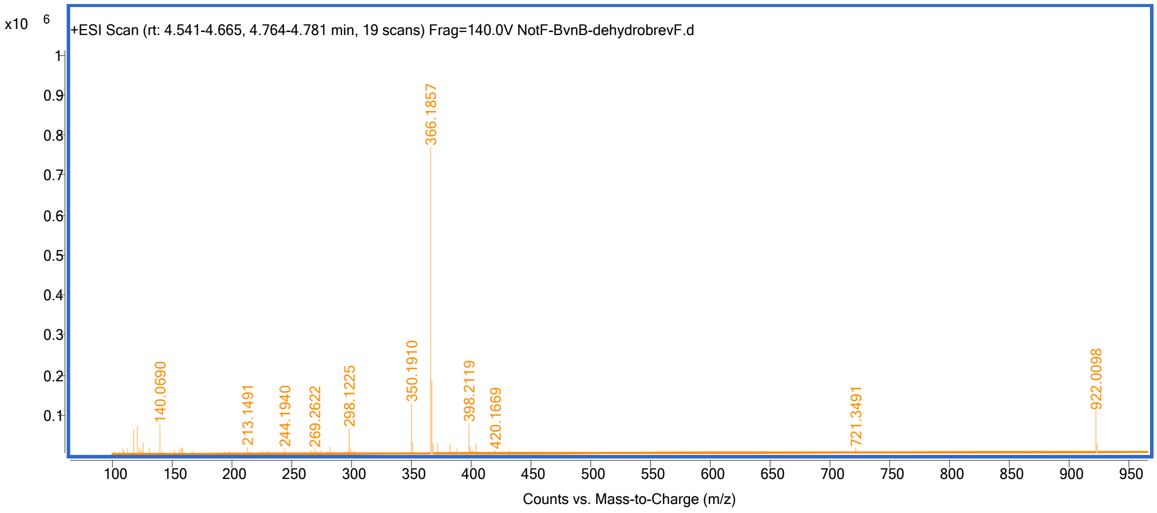

**Supplementary Figure 21.** Mass spectrum (TOF LC/MS) of brevianamide F (**1**).

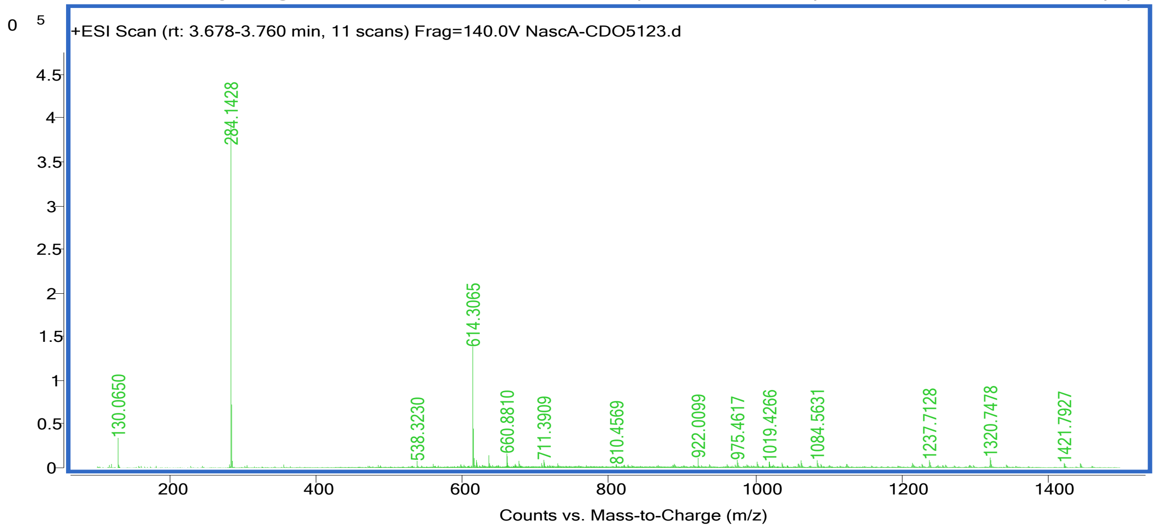

**Supplementary Figure 22.** Mass spectrum (TOF LC/MS) of dehydrobrevianamide F (**2**).

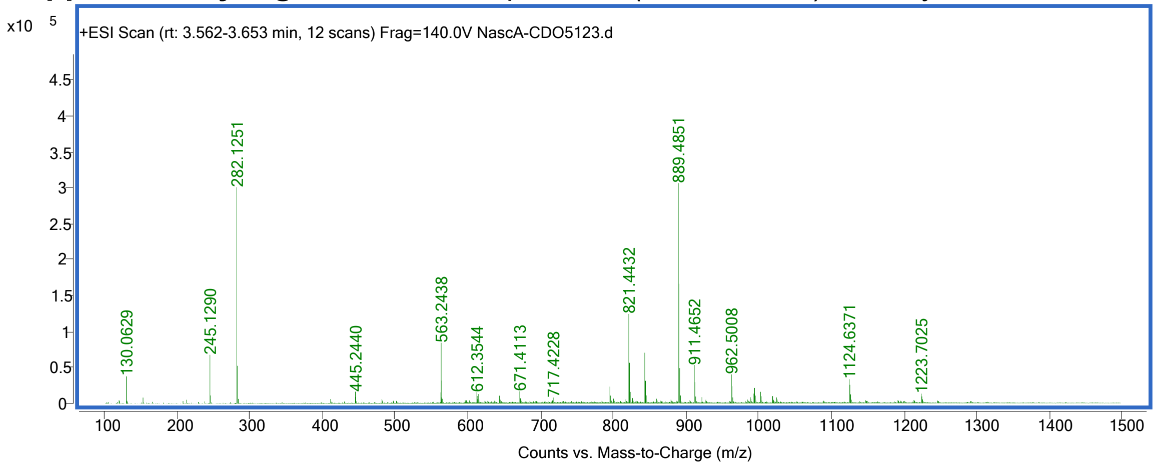

**Supplementary Figure 23.** Mass spectrum (TOF LC/MS) of (+)-dehydrodeoxybrevianamide E (**3**).

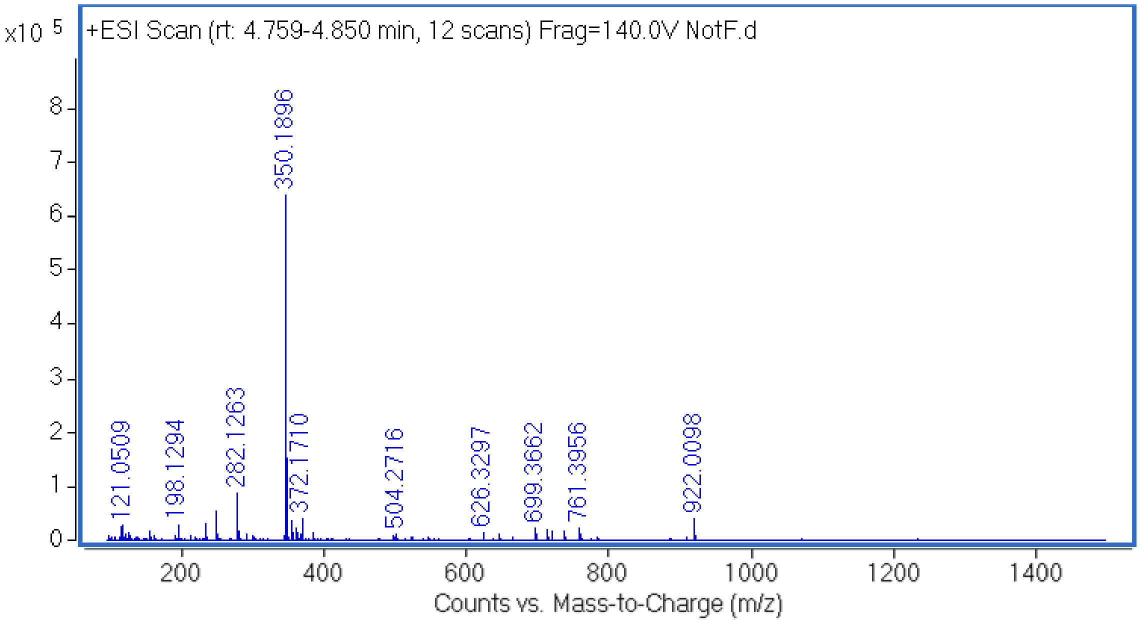

**Supplementary Figure 24.** Mass spectrum (TOF LC/MS) of deoxybrevianamide E (**7**).

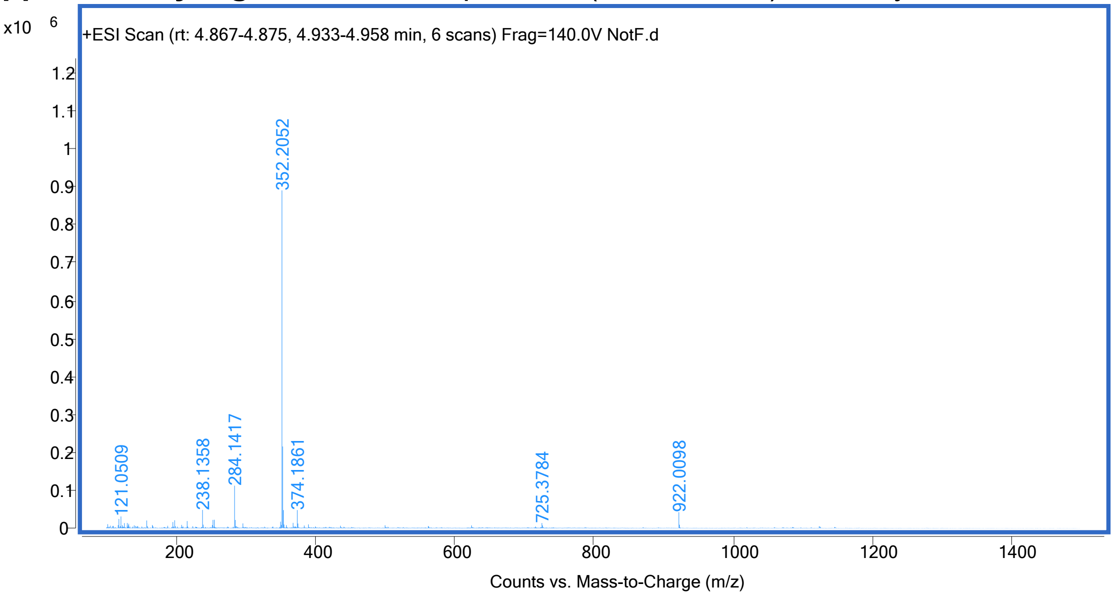

**Supplementary Figure 25.** Mass spectrum (TOF LC/MS) of (-)-dehydrobrevianamide E (**4**).

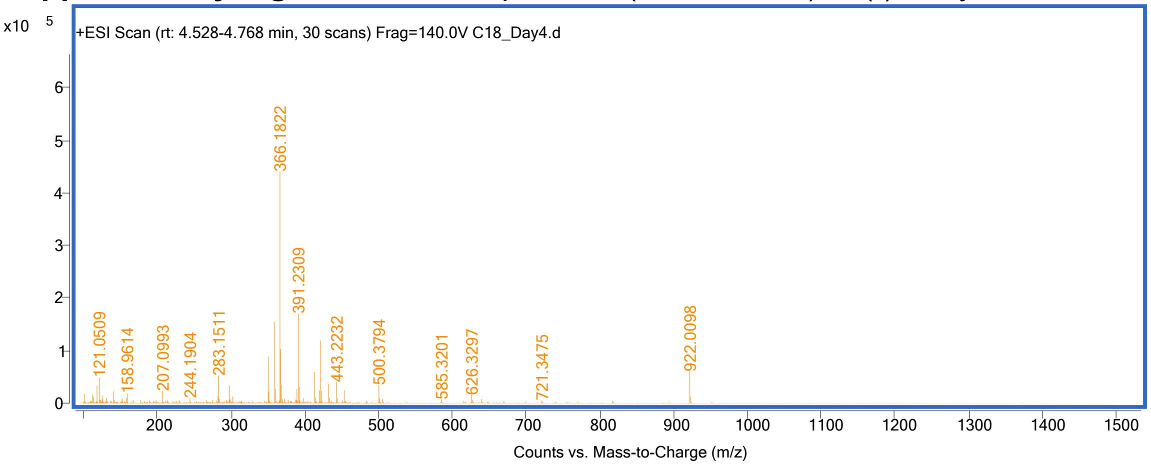

**Supplementary Figure 26.** Mass spectrum (TOF LC/MS) of brevianamide E (**8**).

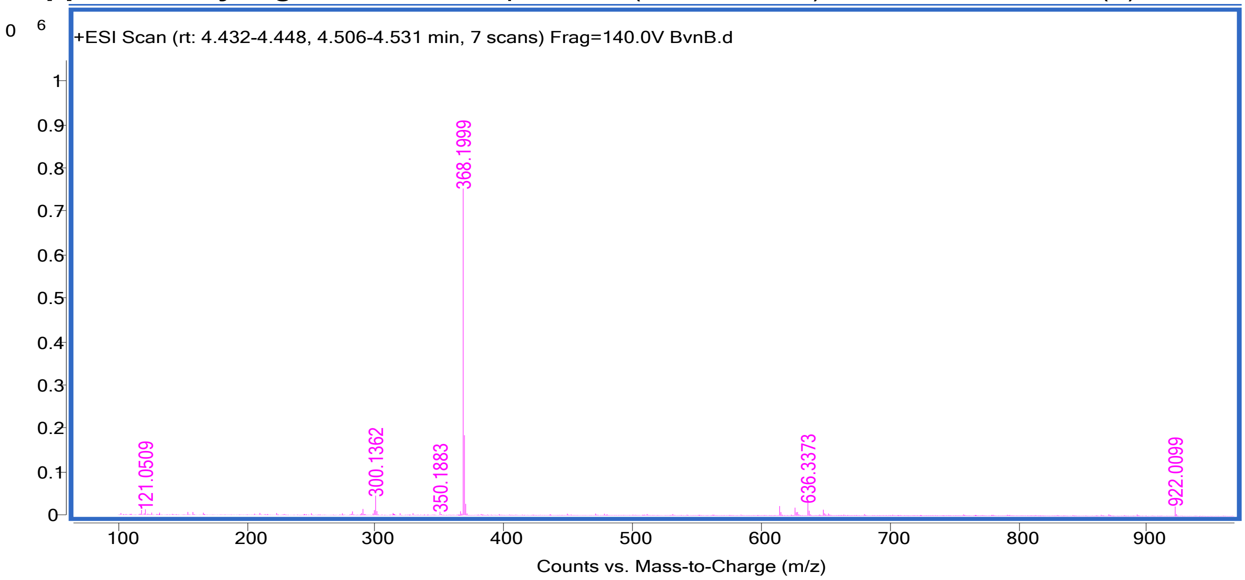

**Supplementary Figure 27.** Mass spectrum (TOF LC/MS) of (+)-brevianamide A (**5**) from fungi extraction.

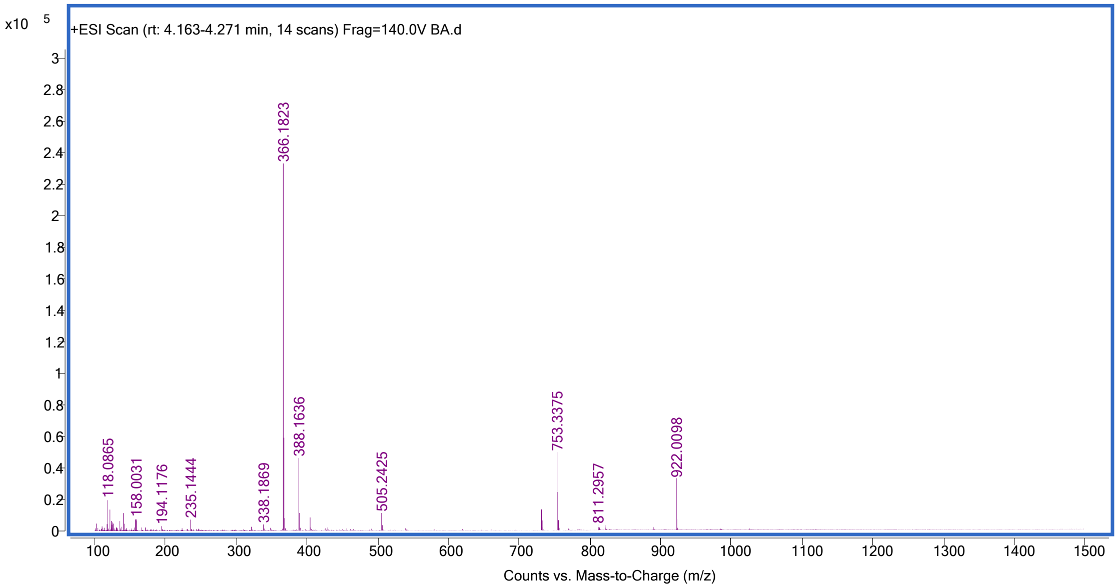

**Supplementary Figure 28.** Mass spectrum (TOF LC/MS) of (+)-brevianamide B (**6**) from fungi extraction.

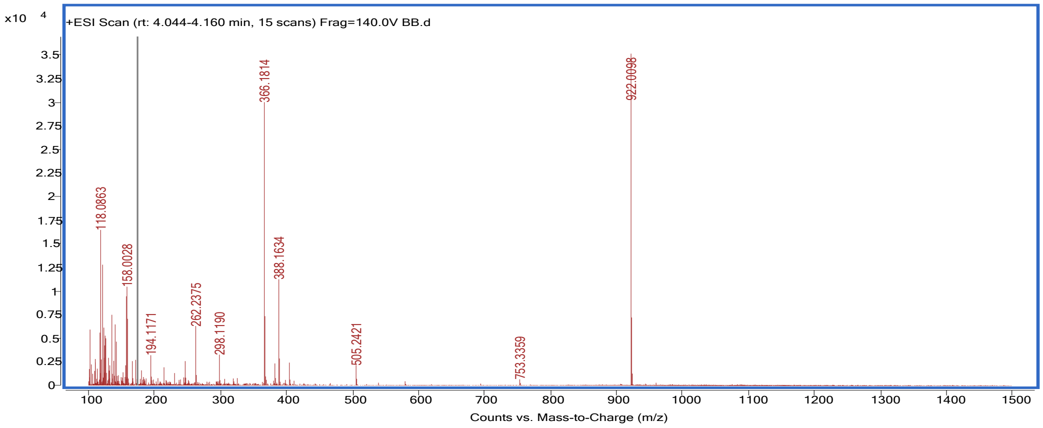

**Supplementary Figure 29.** Mass spectrum (TOF LC/MS) of (+)-brevianamide A (**5**) from LiOH reaction.

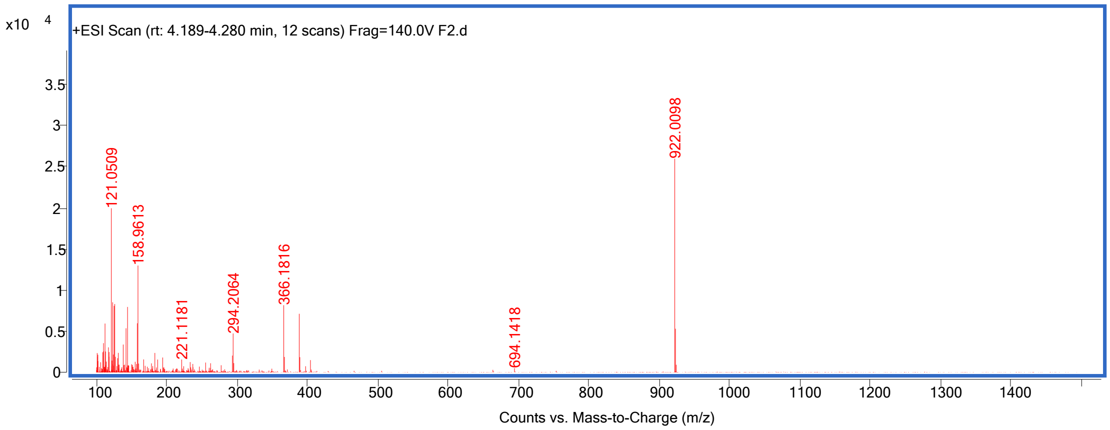

**Supplementary Figure 30.** Mass spectrum (TOF LC/MS) of (+)-brevianamide B (**6**) from LiOH reaction.

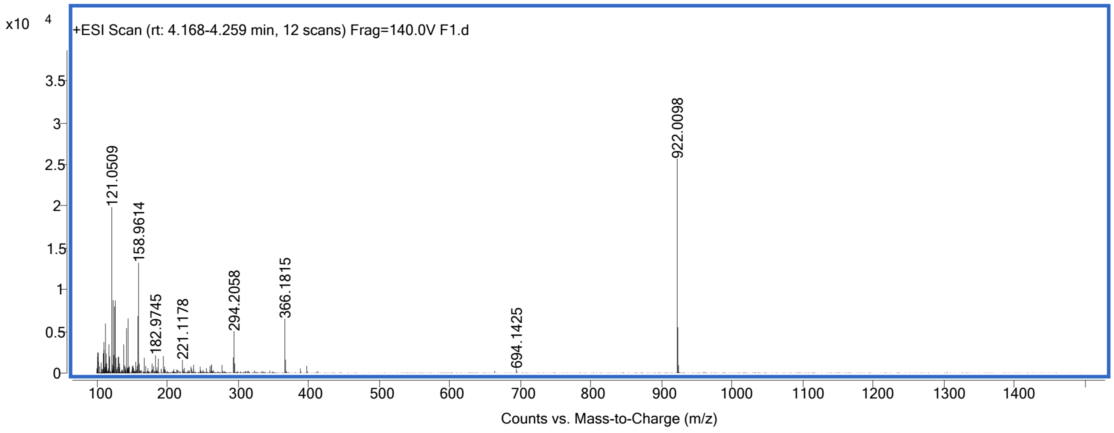

**Supplementary Figure 31.** [1]H NMR spectrum of brevianamide F (**1**) (599 MHz, $CDCl_3$)

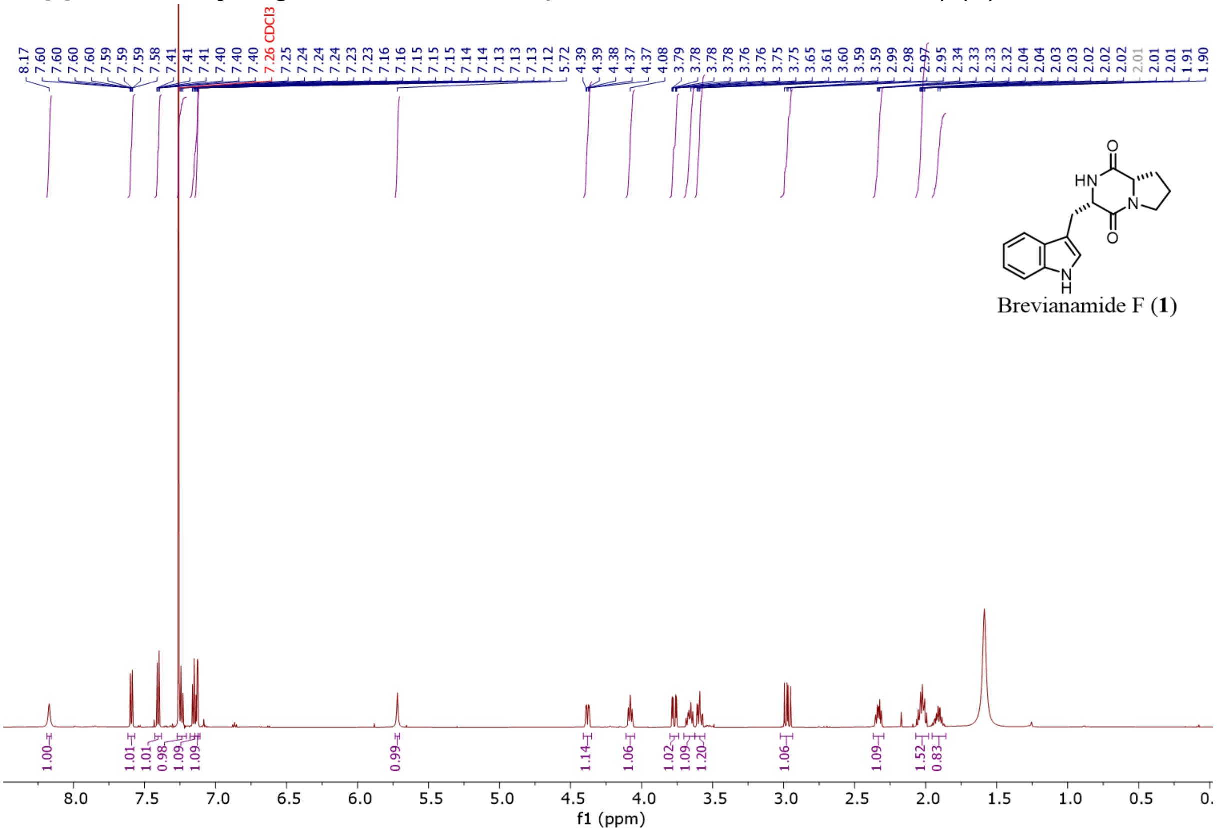

**Supplementary Figure 32**. $^{13}C$ NMR spectrum of brevianamide F (**1**) (151 MHz, $CDCl_3$)

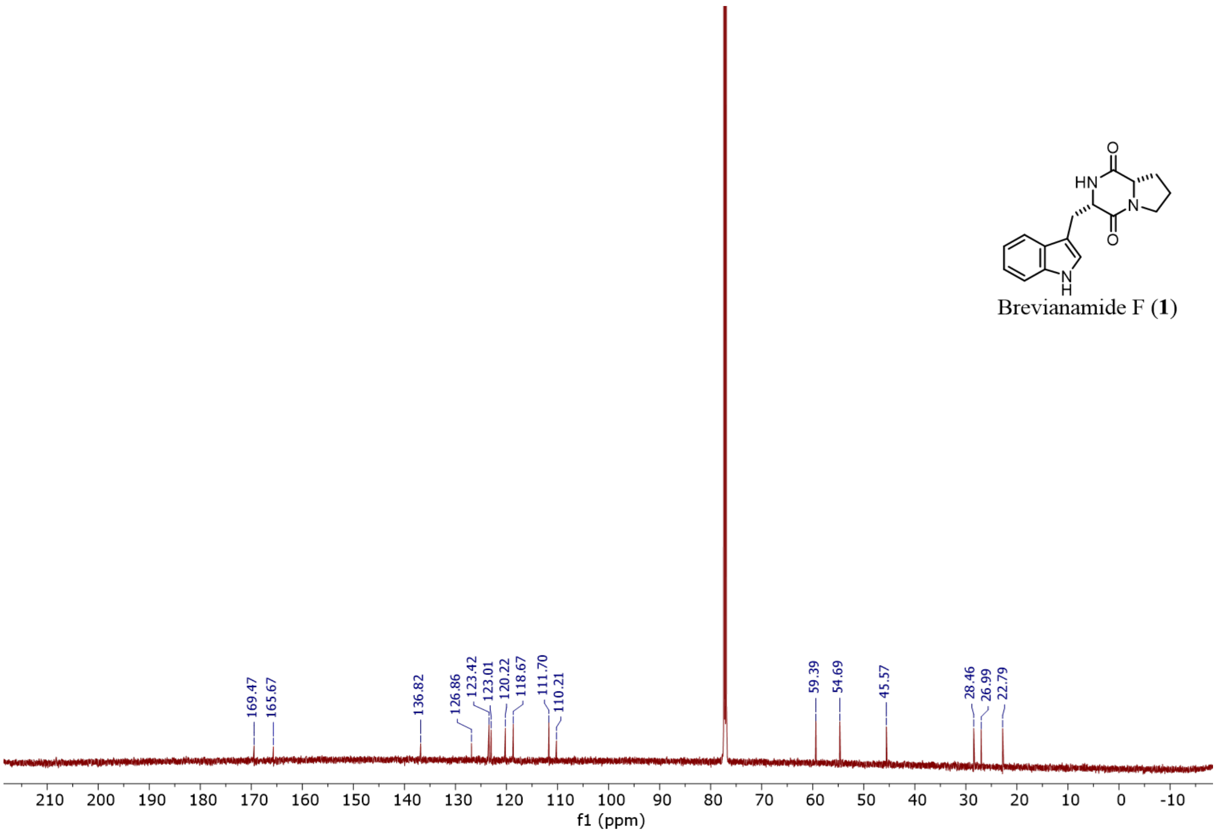

**Supplementary Figure 33.** $^{1}H$ NMR spectrum of dehydrobrevianamide F (**2**) (599 MHz, $CDCl_3$)

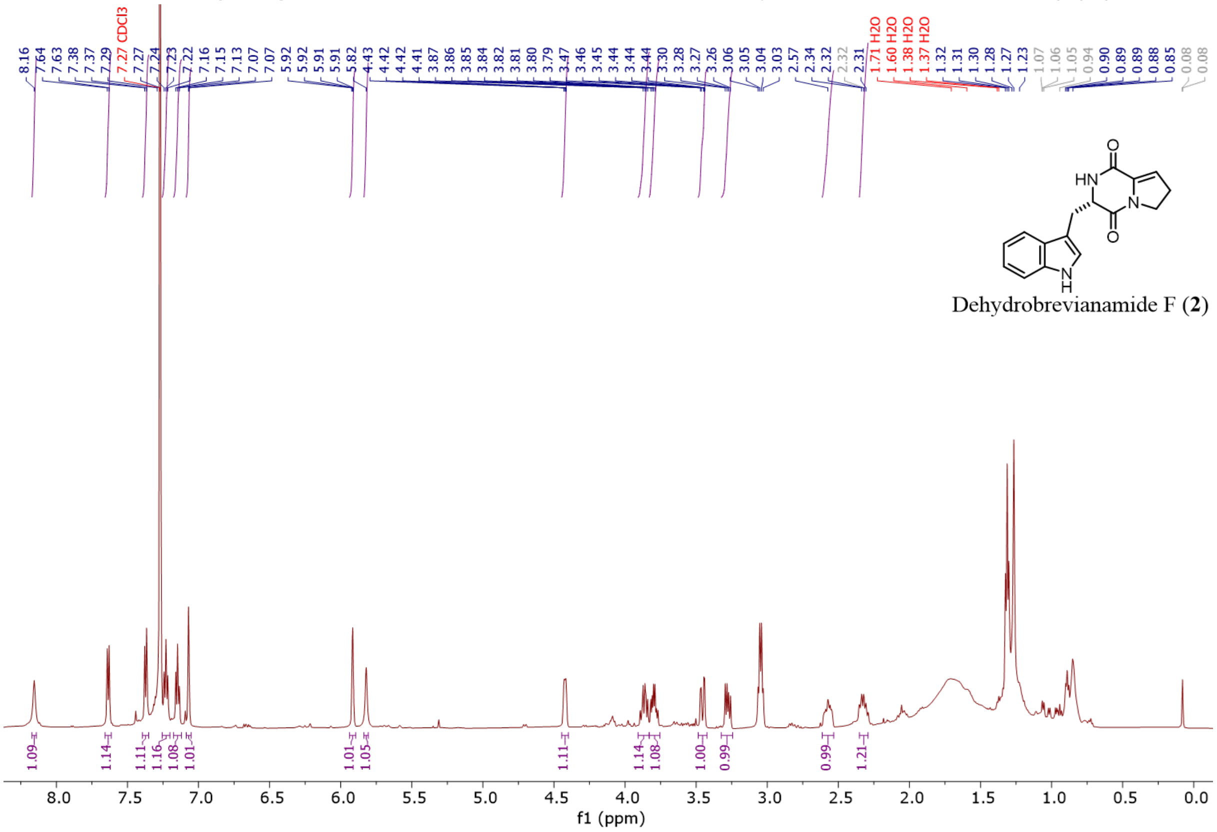

**Supplementary Figure 34.** $^{13}C$ NMR spectrum of dehydrobrevianamide F (**2**) (151 MHz, $CDCl_3$)

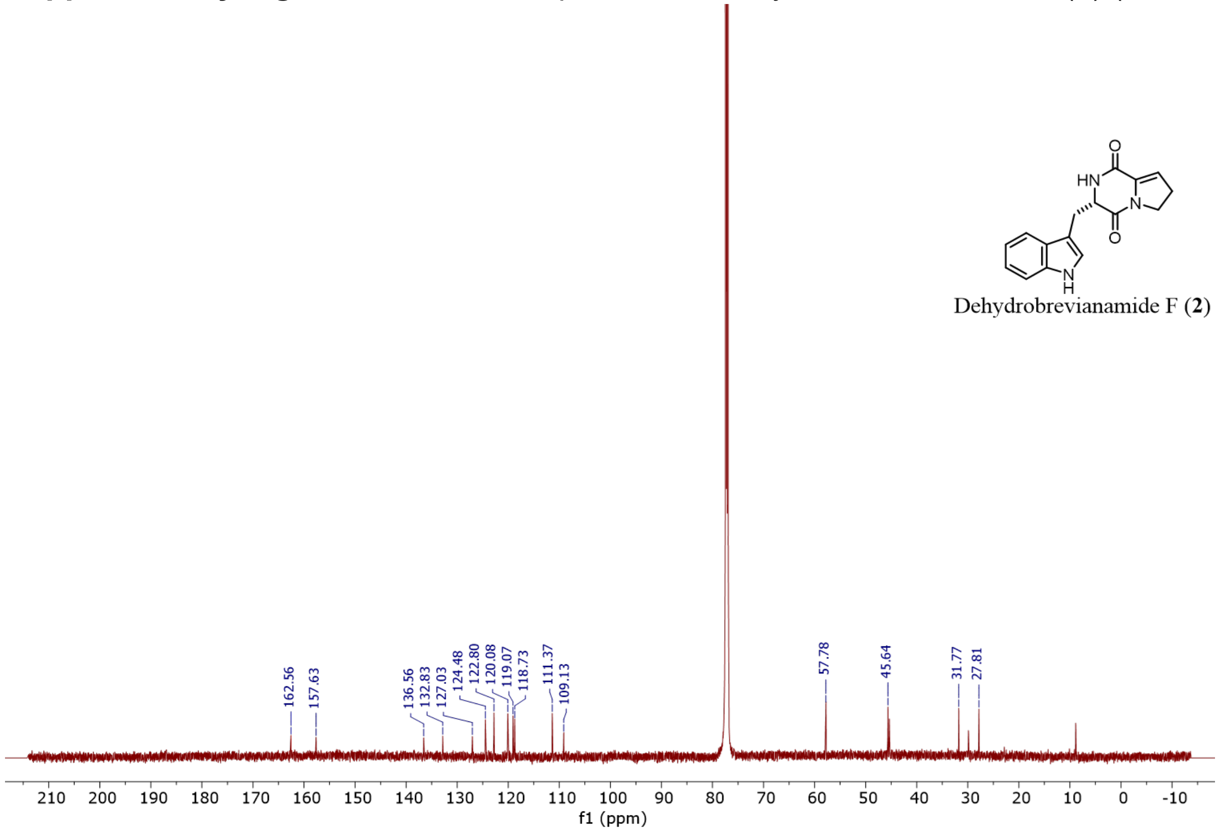

**Supplementary Figure 35.** COSY NMR spectrum of dehydrobrevianamide F (**2**) ($CDCl_3$)

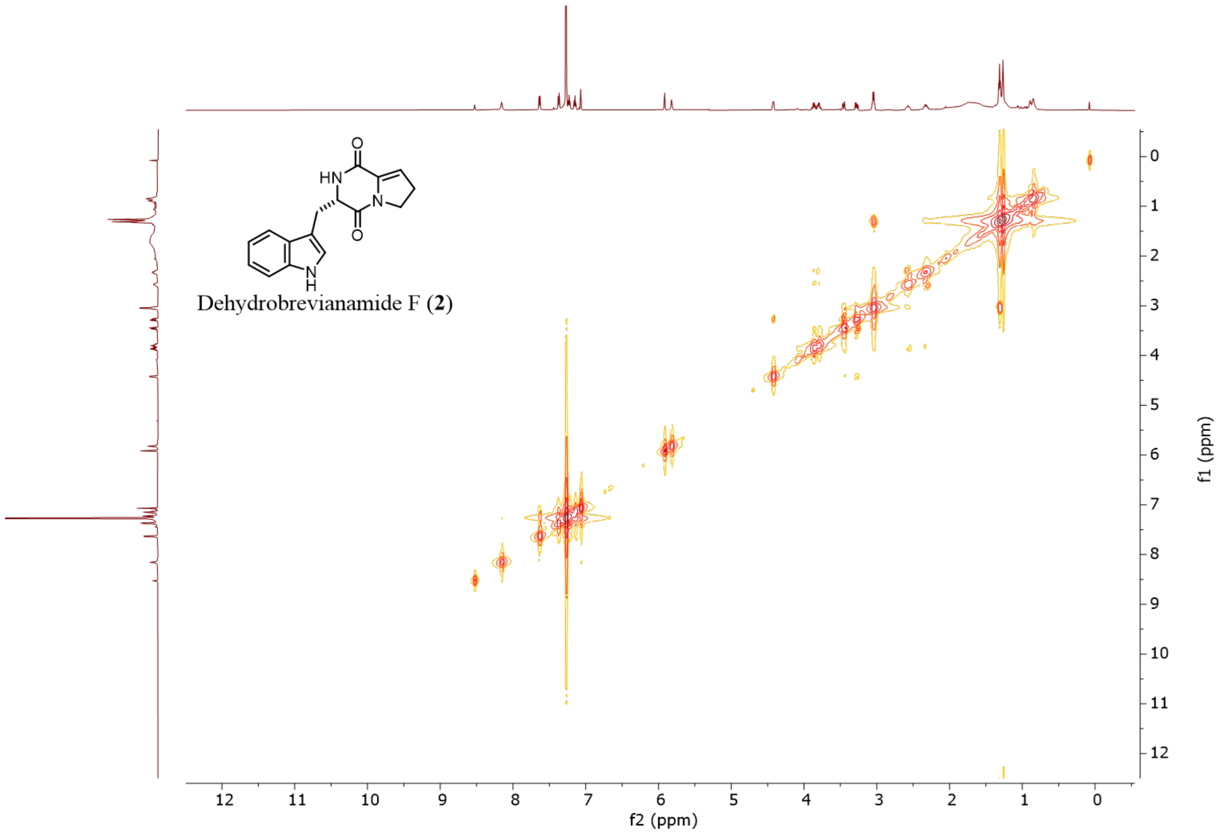

**Supplementary Figure 36.** HSQC NMR spectrum of dehydrobrevianamide F (**2**) ($CDCl_3$)

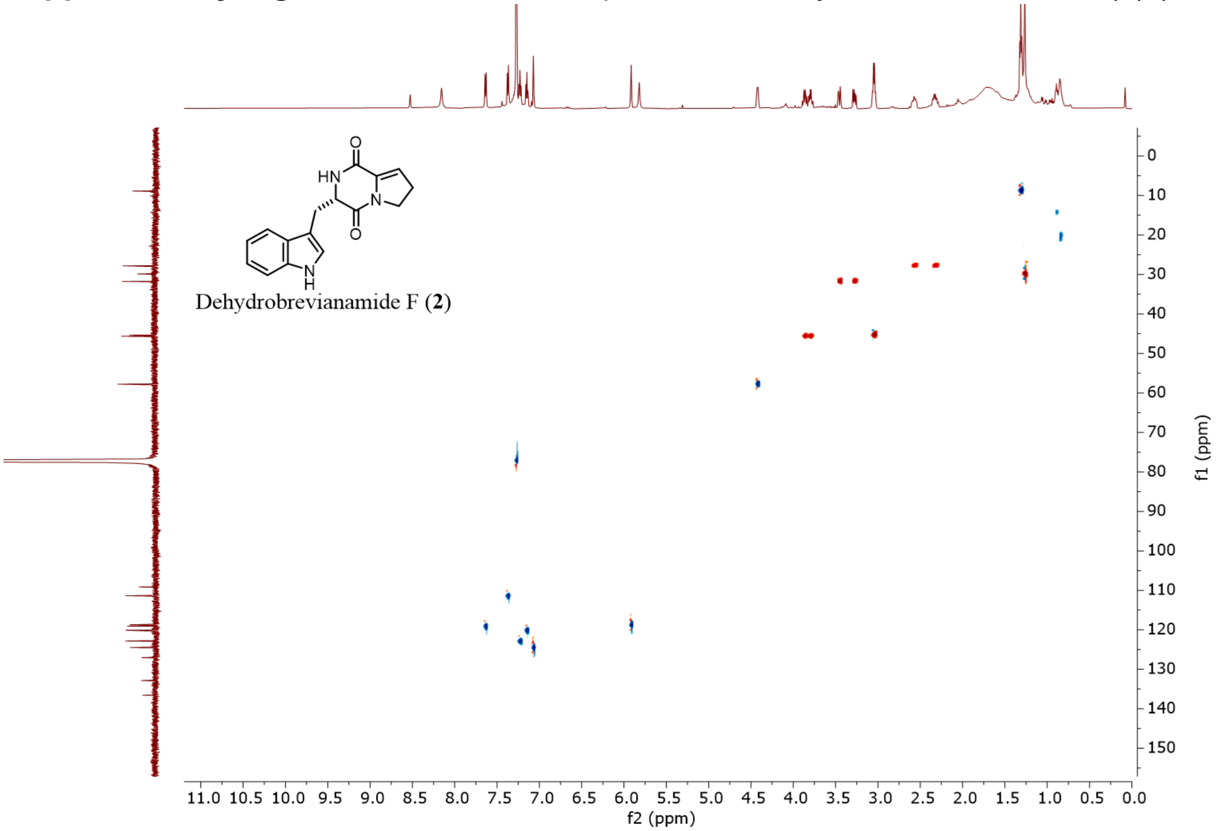

**Supplementary Figure 37.** HMBC NMR spectrum of dehydrobrevianamide F (**2**) ($CDCl_3$)

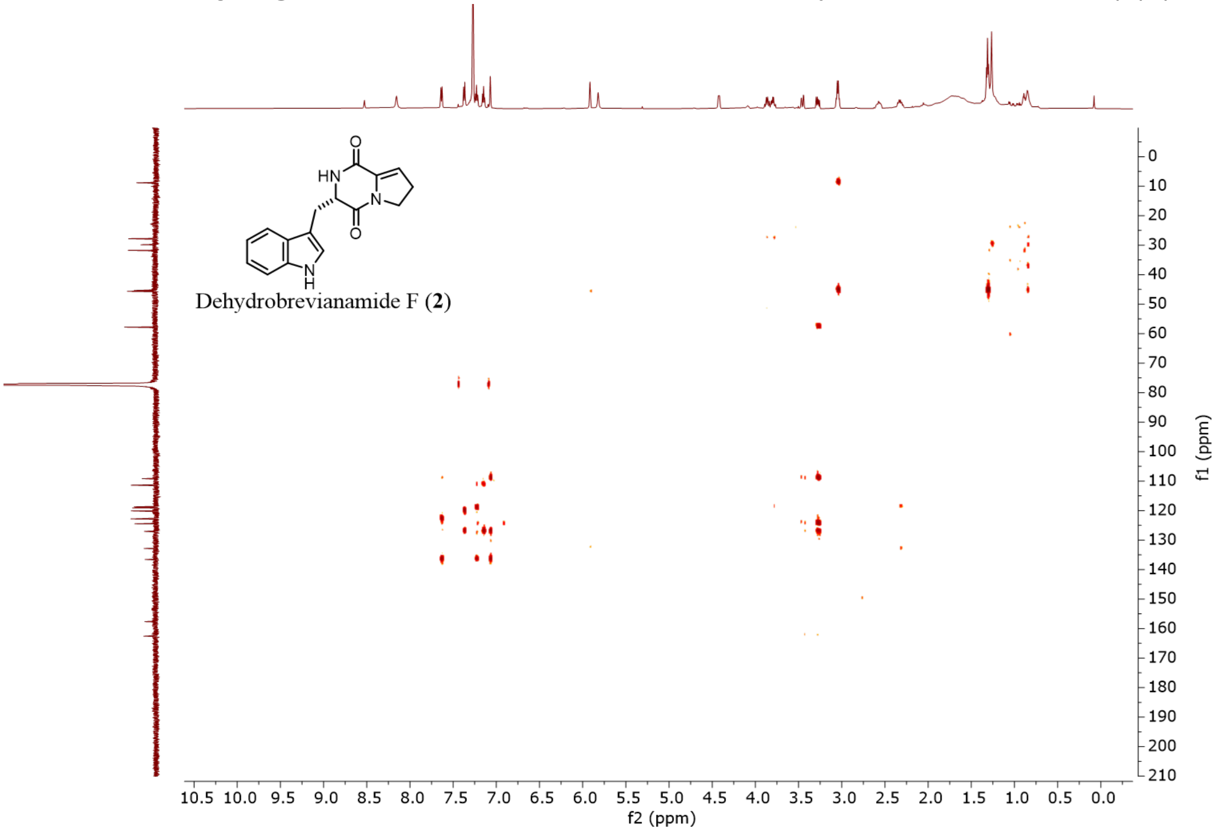

**Supplementary Figure 38.** NOESY NMR spectra of dehydrobrevianamide F (**2**) ($CDCl_3$)

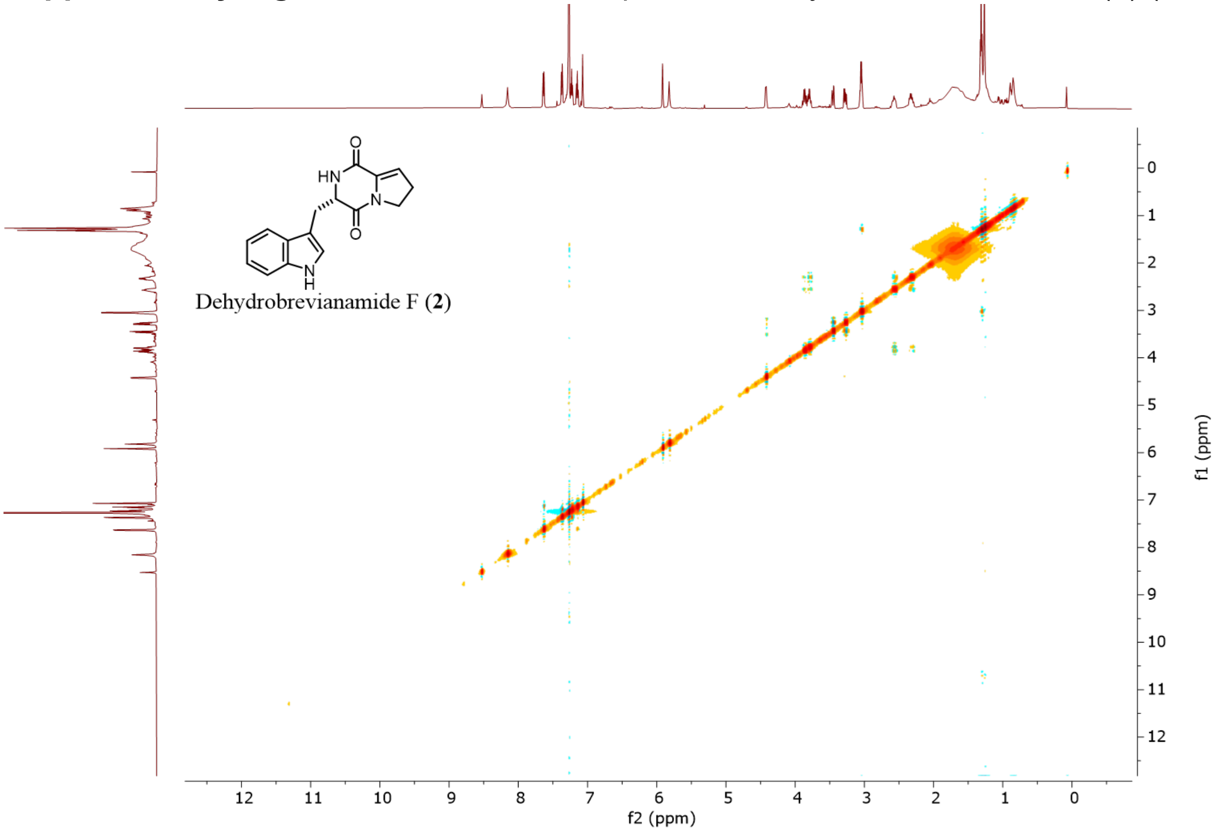

**Supplementary Figure 39.** [1]H NMR spectrum of dehydrobrevianamide F (**2**) (DMSO-d6)

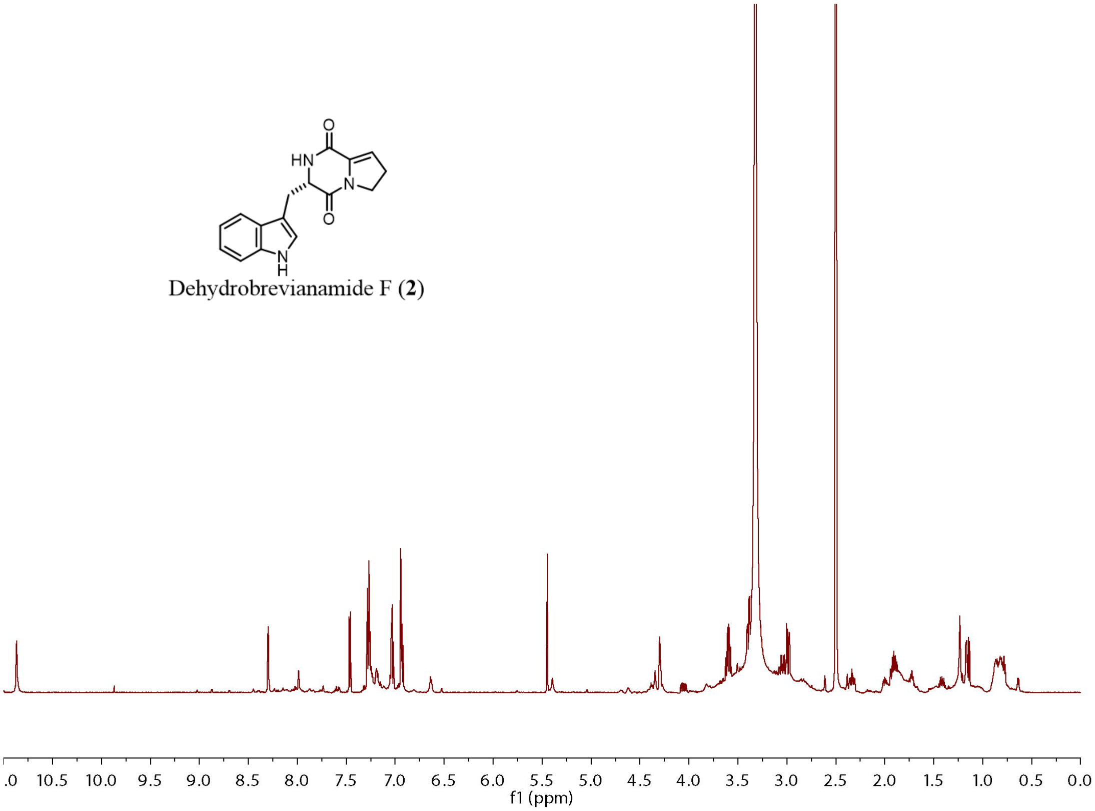

**Supplementary Figure 40.** $^{13}C$ NMR spectrum of dehydrobrevianamide F (**2**) (DMSO-d6)

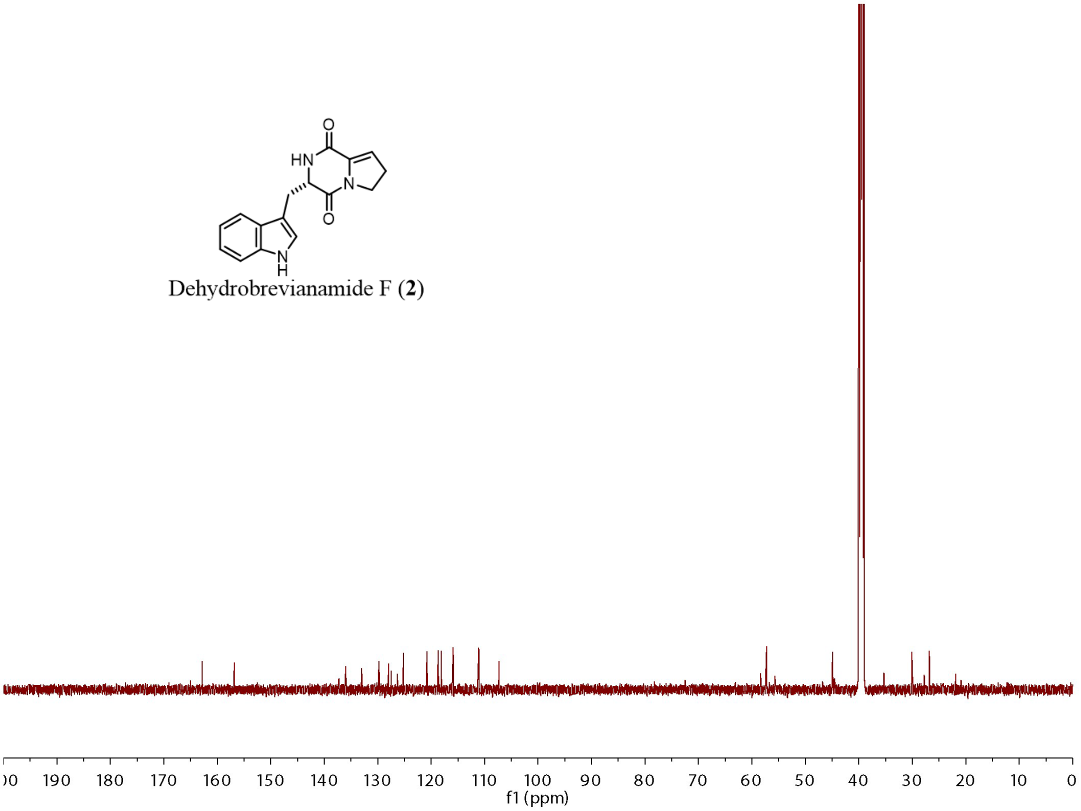

**Supplementary Figure 41.** COSY NMR spectrum of dehydrobrevianamide F (**2**) (DMSO-d6)

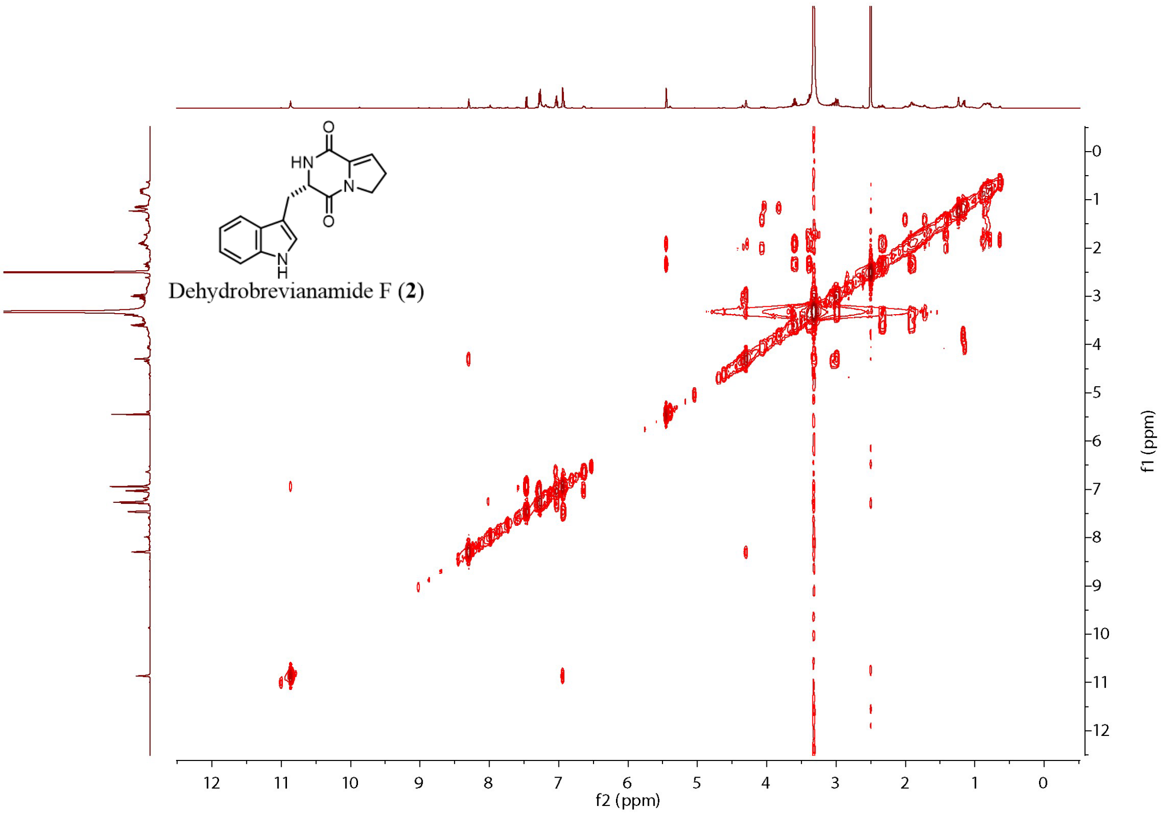

**Supplementary Figure 42.** HSQC NMR spectra of dehydrobrevianamide F (**2**) (DMSO-d6)

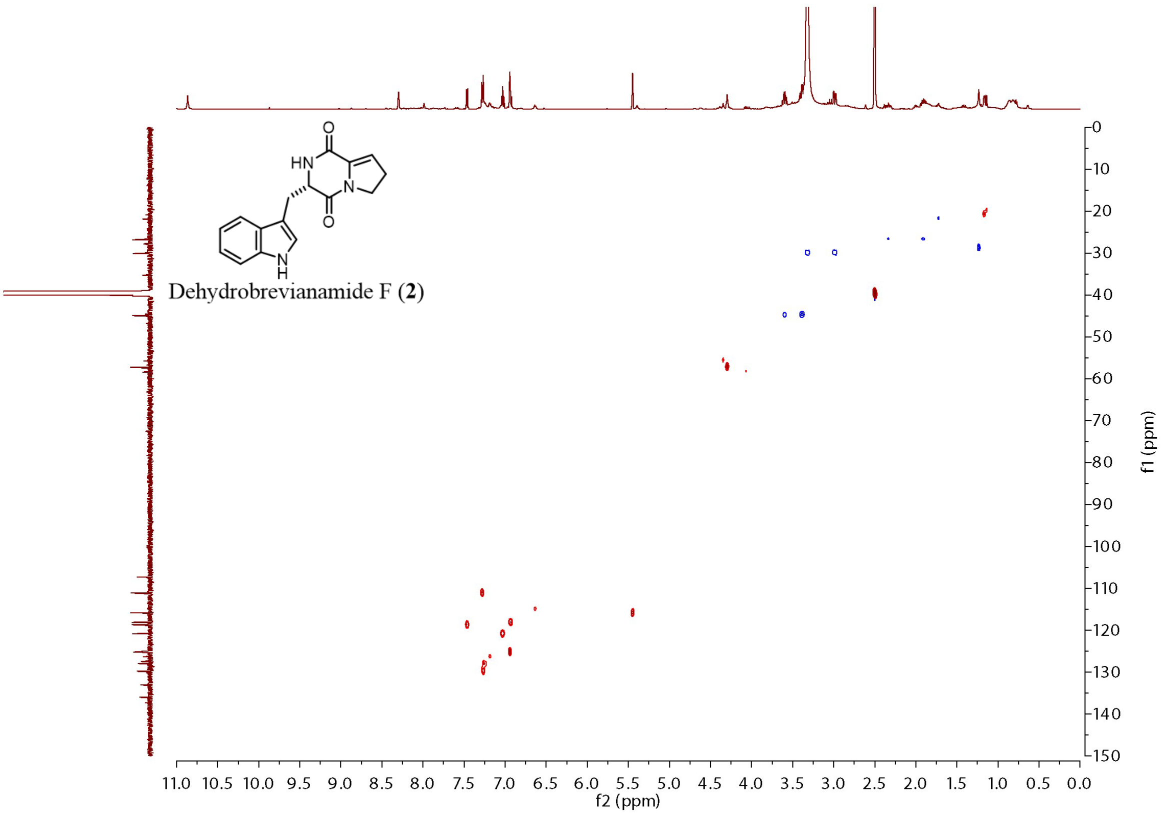

**Supplementary Figure 43.** HMBC NMR spectra of dehydrobrevianamide F (**2**) (DMSO-d6)

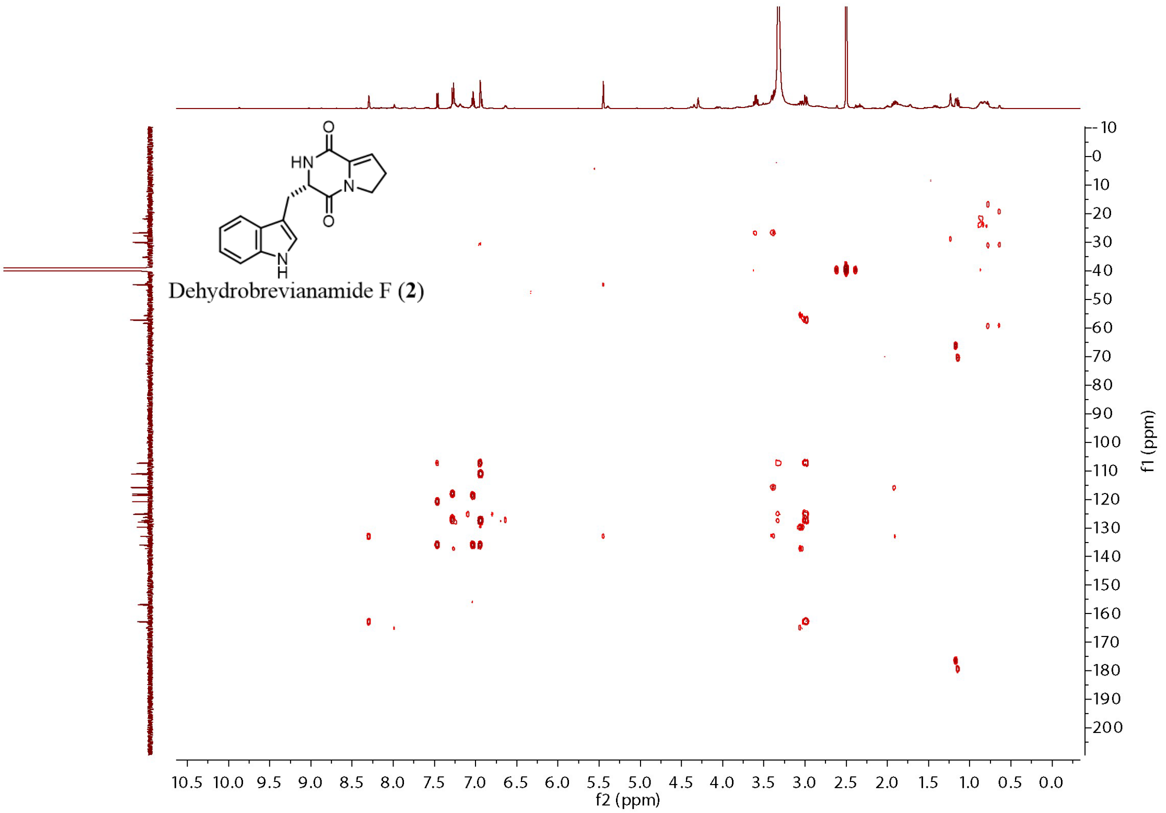

**Supplementary Figure 44.** $^{1}$H NMR spectrum of (+)-dehydrodeoxybrevianamide E (**3**) (599 MHz, $CDCl_3$)

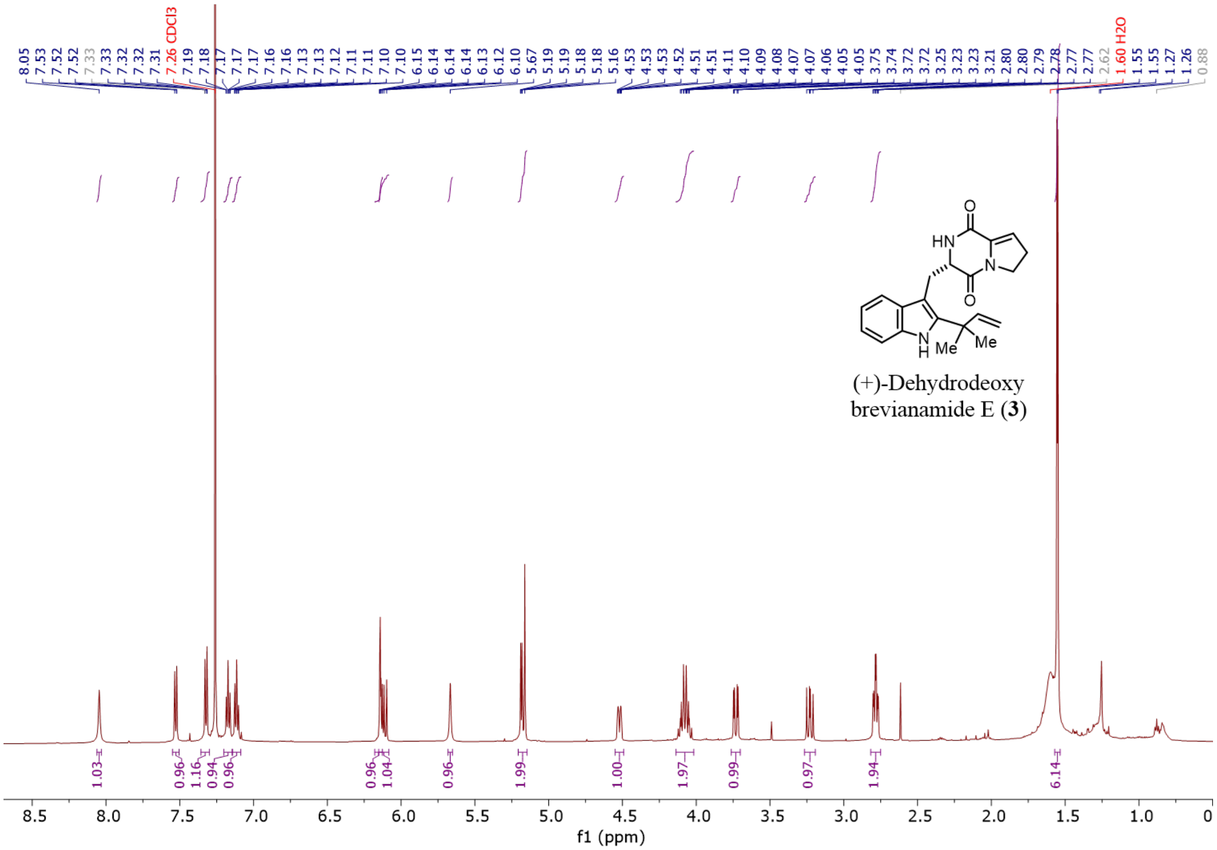

**Supplementary Figure 45.** $^{13}$C NMR spectrum of (+)-dehydrodeoxybrevianamide E (**3**) (151 MHz, $CDCl_3$)

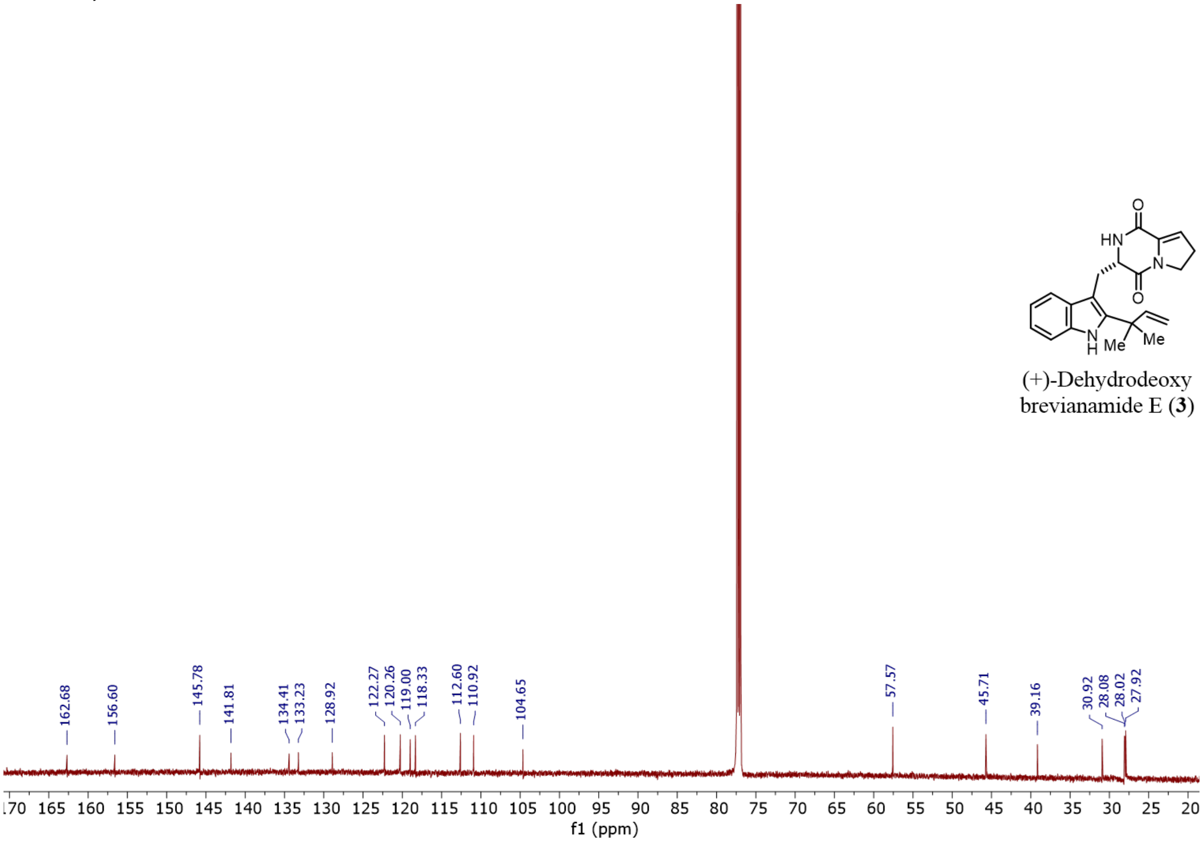

**Supplementary Figure 46.** $^1$H NMR spectrum of deoxybrevianamide E (**7**) (599 MHz, $CDCl_3$)

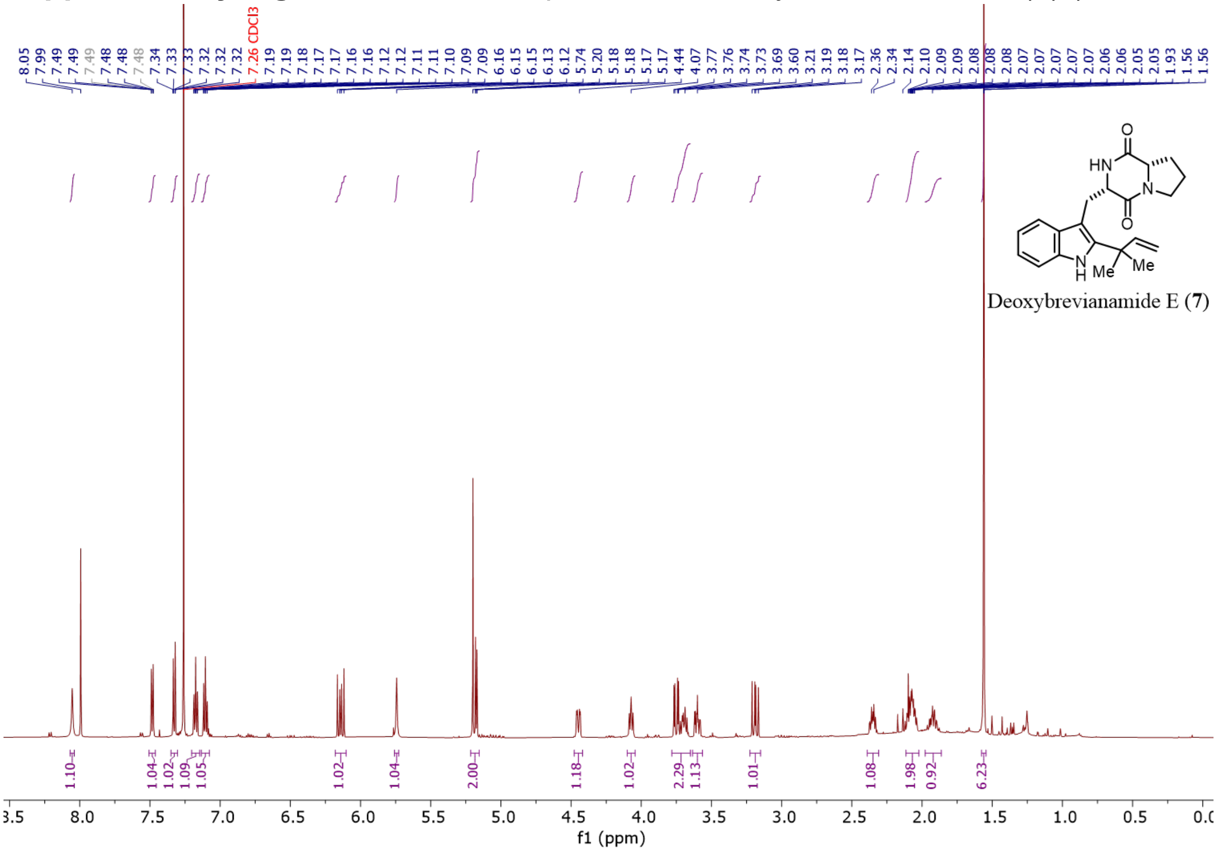

**Supplementary Figure 47.** $^{13}$C NMR spectrum of deoxybrevianamide E (**7**) (151 MHz, $CDCl_3$)

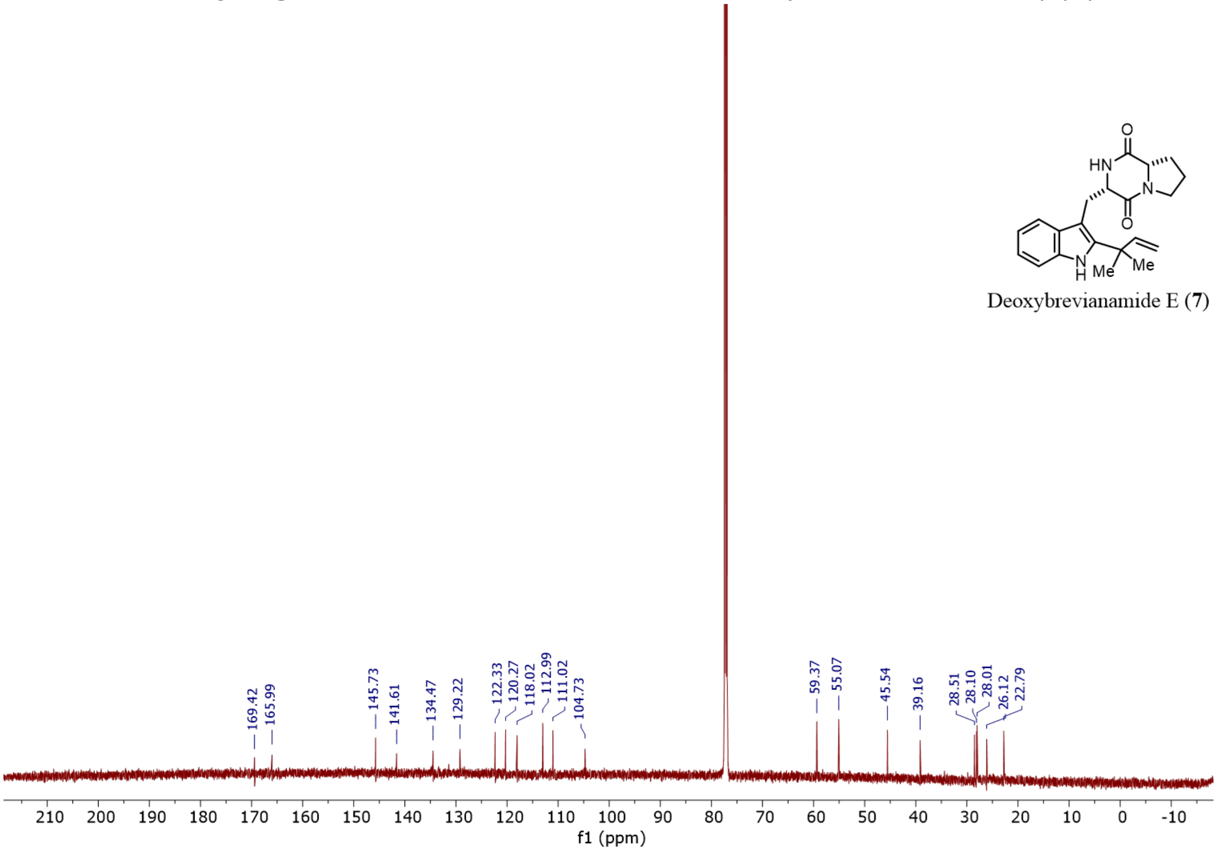

**Supplementary Figure 48.** $^{1}$H NMR spectrum of (-)-dehydrobrevianamide E (**4**) (599 MHz, $CDCl_3$)

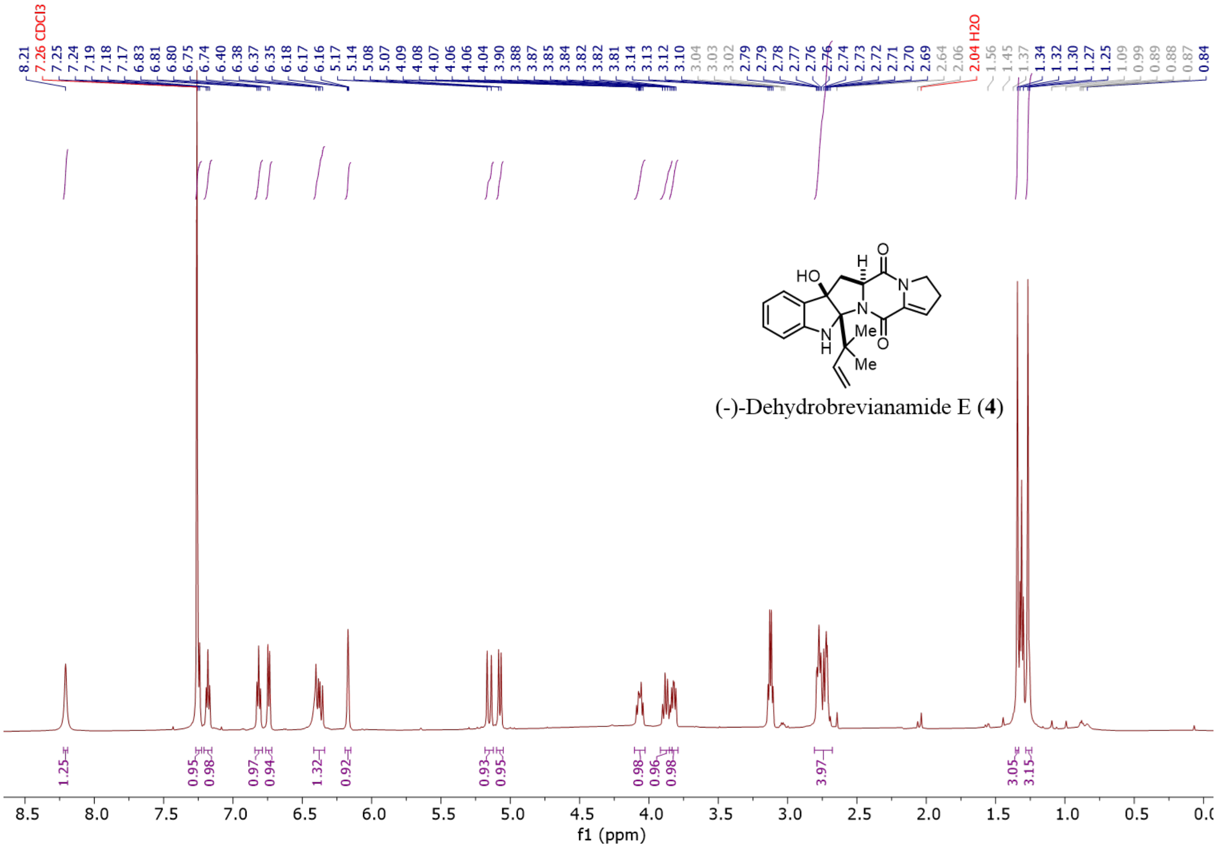

**Supplementary Figure 49.** $^{13}$C NMR spectrum of (-)-dehydrobrevianamide E (**4**) (151 MHz, $CDCl_3$)

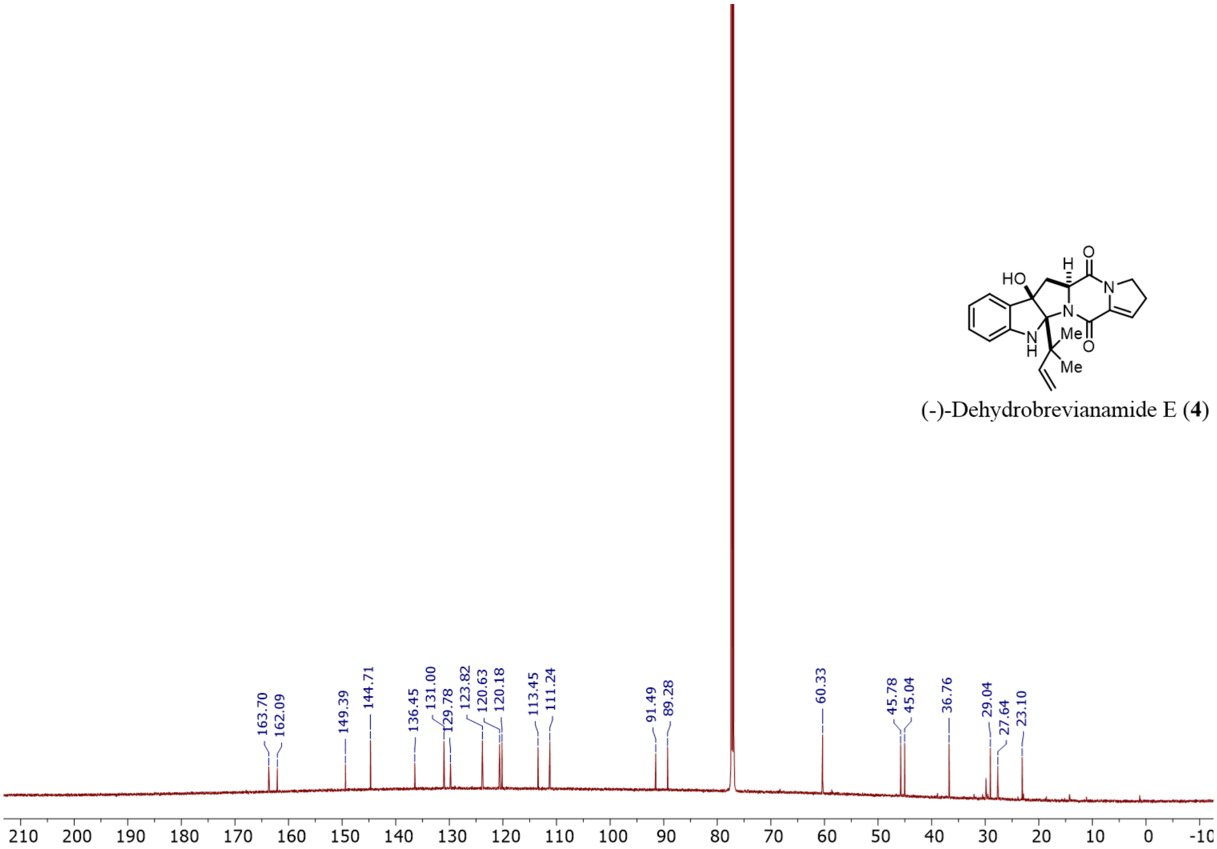

**Supplementary Figure 50.** $^{1}$H NMR spectrum of brevianamide E (**8**) (599 MHz, $CDCl_3$)

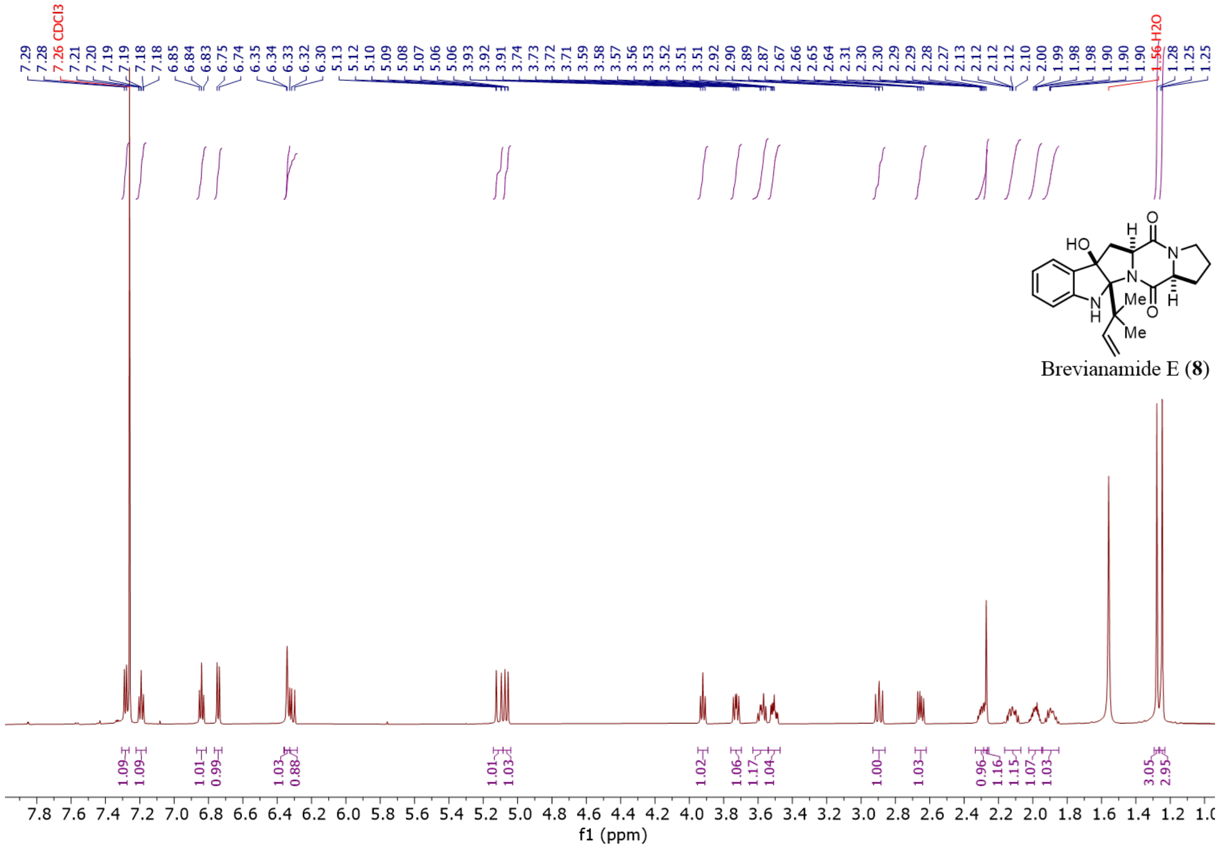

**Supplementary Figure 51.** $^{13}$C NMR spectrum of brevianamide E (**8**) (151 MHz, $CDCl_3$)

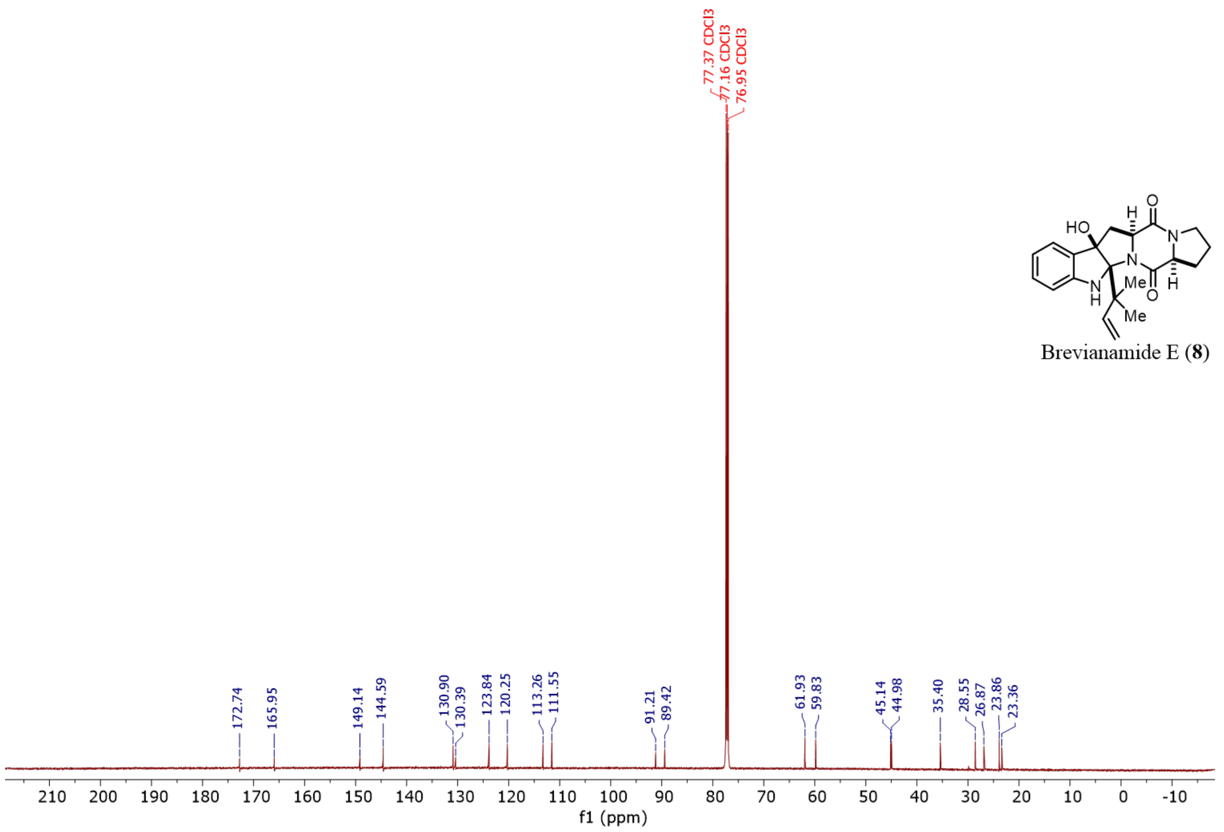

**Supplementary Figure 52.** $^{1}H$ NMR spectrum of (+)-brevianamide A (5) from fungal extraction (599 MHz, $CDCl_3$)

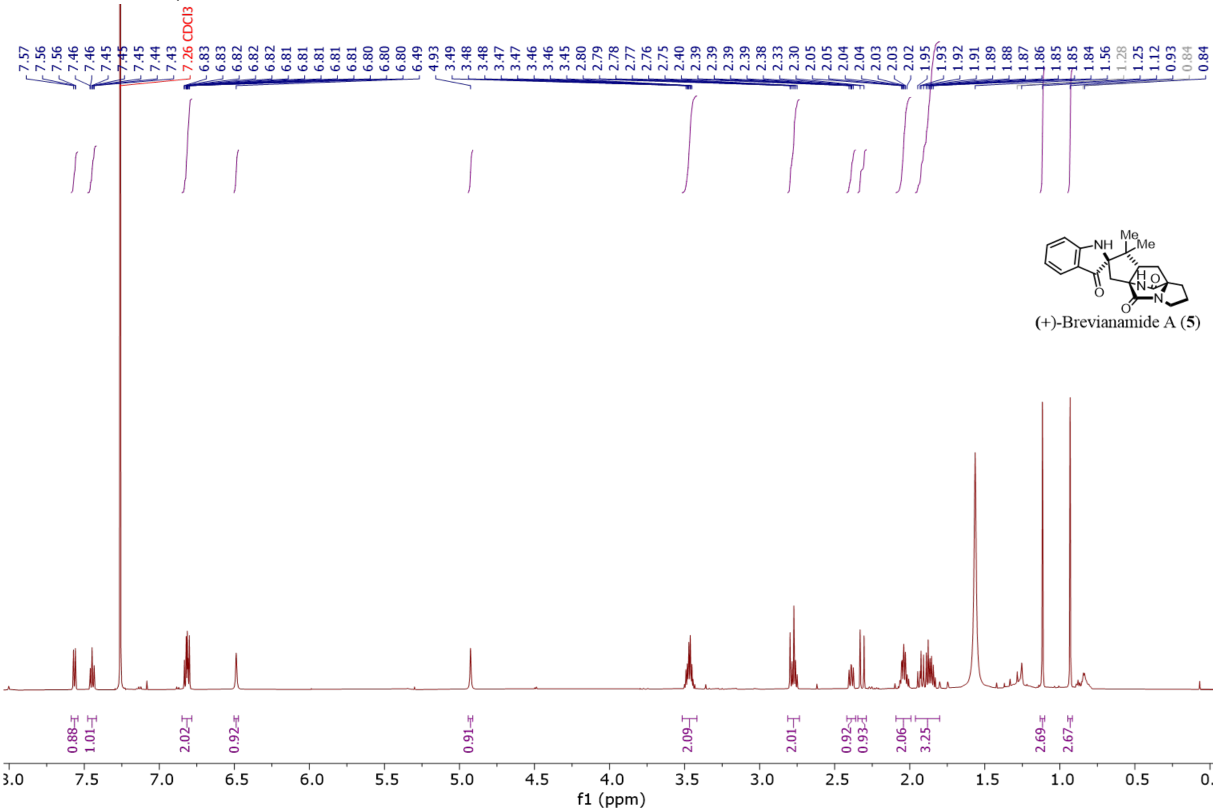

**Supplementary Figure 53.** $^{13}C$ NMR spectrum of (+)-brevianamide A (**5**) from fungal extraction (151 MHz, $CDCl_3$)

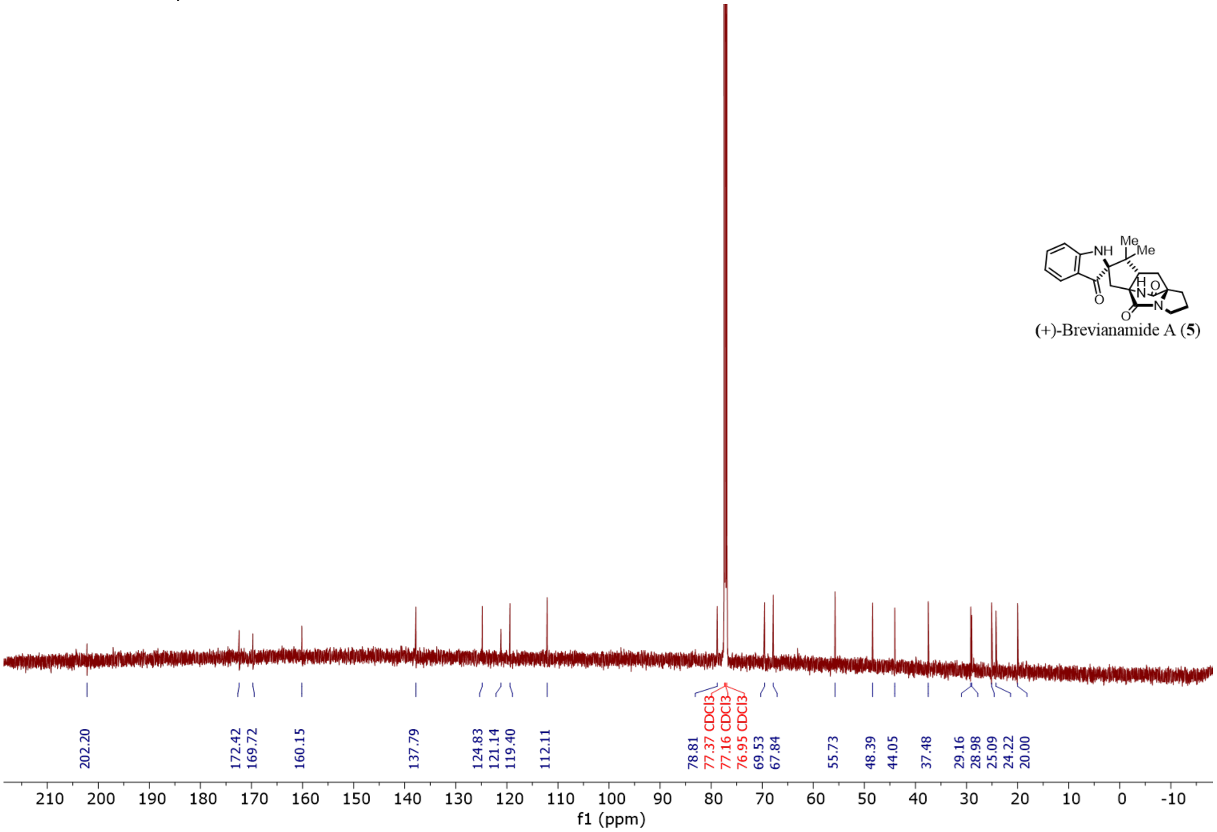

**Supplementary Figure 54.** $^{1}$H NMR spectrum of (+)-brevianamide B (**6**) from fungal extraction (800 MHz, DMSO-d6)

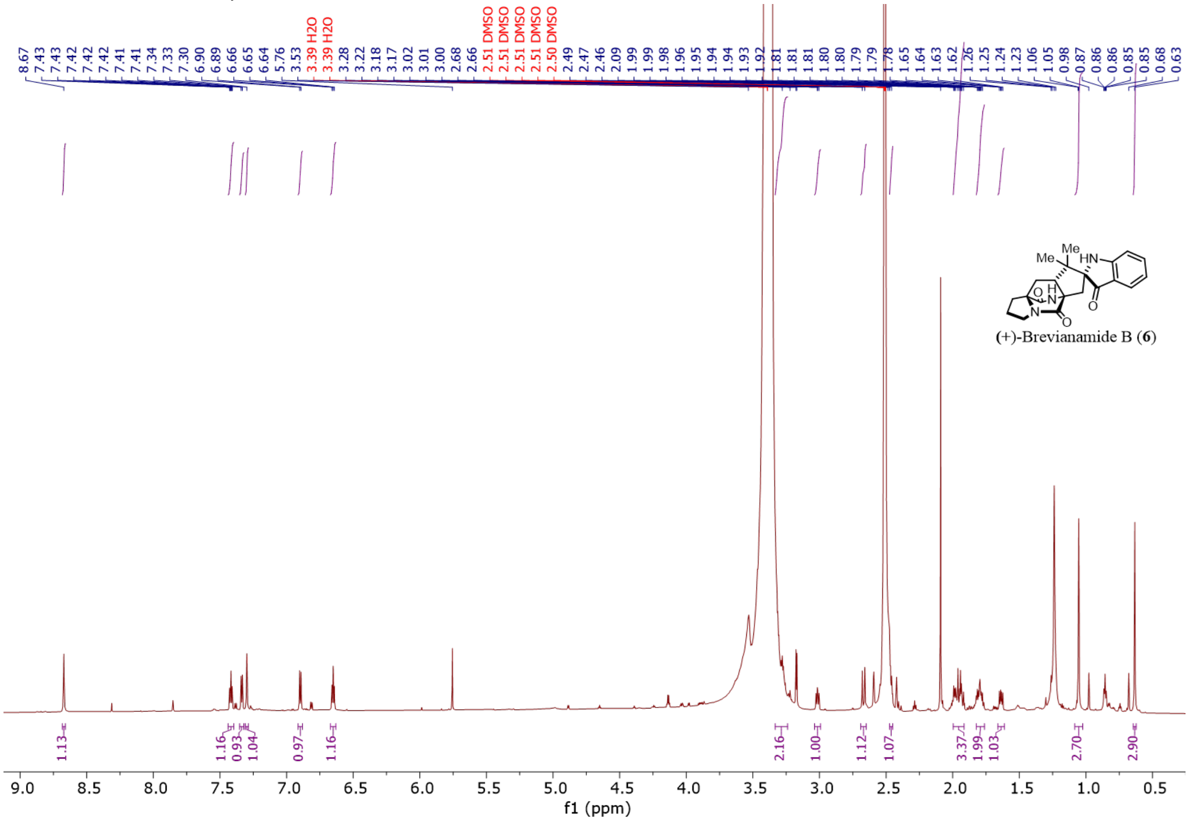

**Supplementary Figure 55.** $^{13}$C NMR spectrum of (+)-brevianamide B (**6**) from fungal extraction (201 MHz, DMSO-d6)

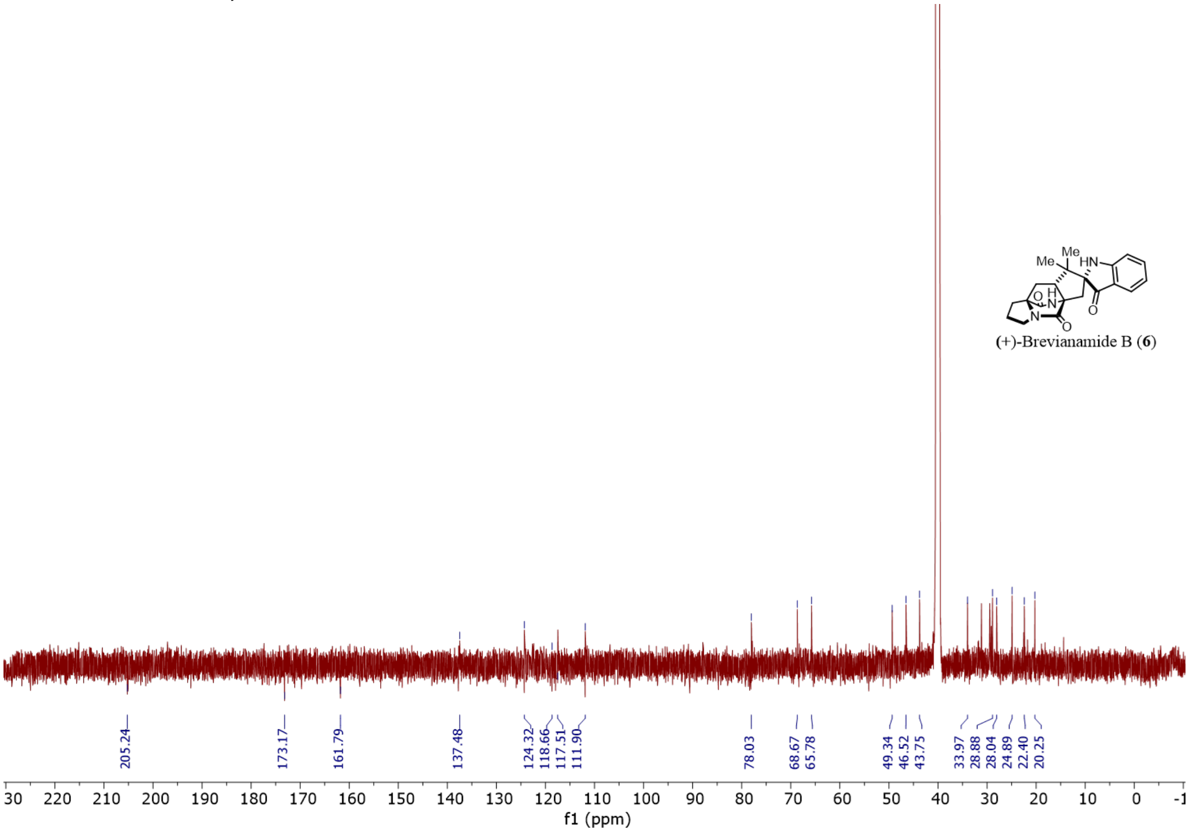

**Supplementary Figure 56.** $^{1}H$ NMR spectrum of (+)-brevianamide B (**6**) from fungal extraction (599 MHz, $CDCl_3$)

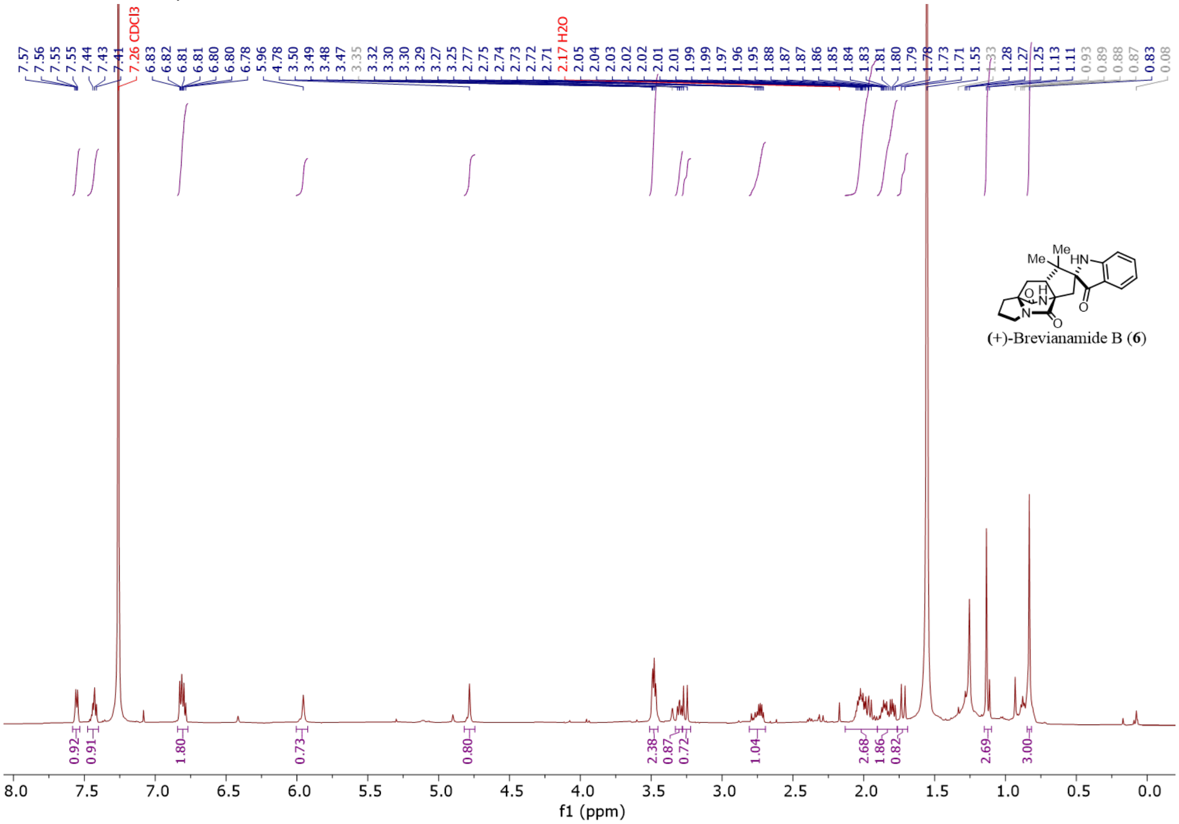

**Supplementary Figure 57.** $^{1}H$ NMR spectrum of (+)-brevianamide A (**5**) from LiOH reaction (599 MHz, $CDCl_3$)

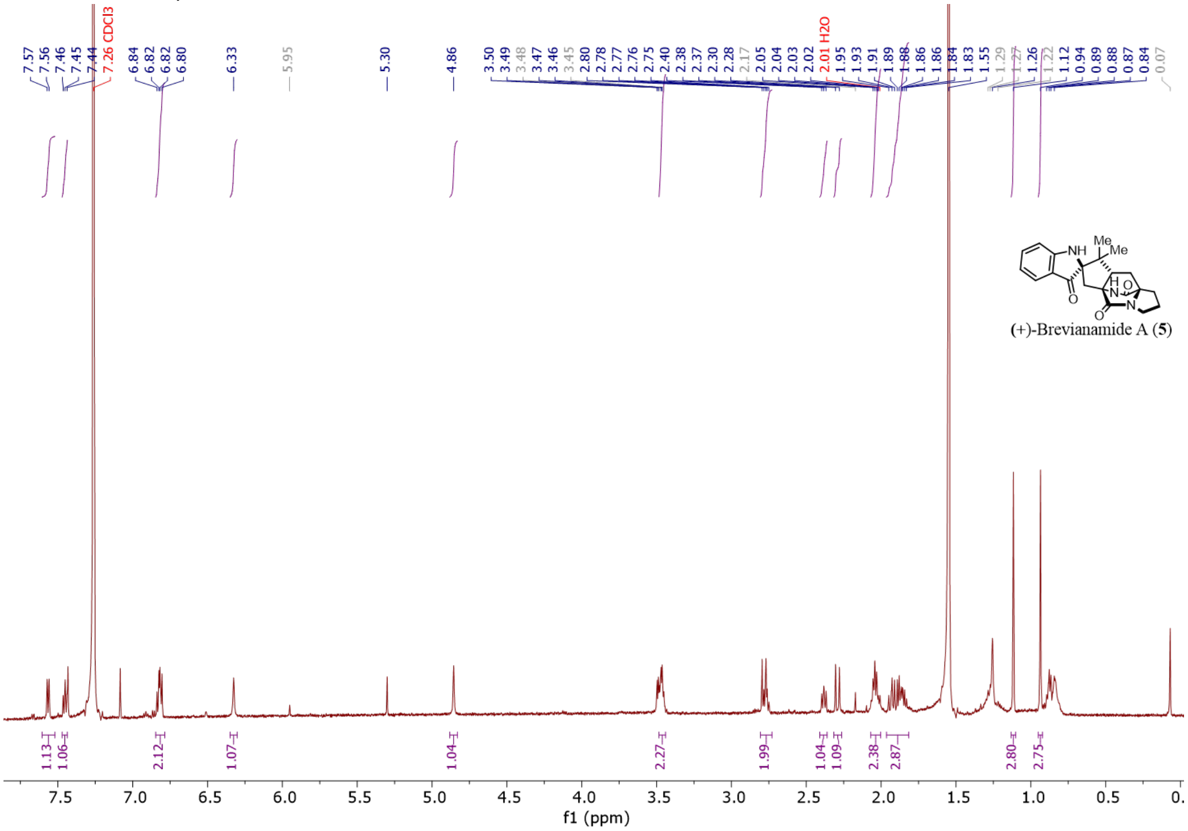

**Supplementary Figure 58.** $^{13}$C NMR spectrum of (+)-brevianamide A (**5**) from LiOH reaction (151 MHz, $CDCl_3$)

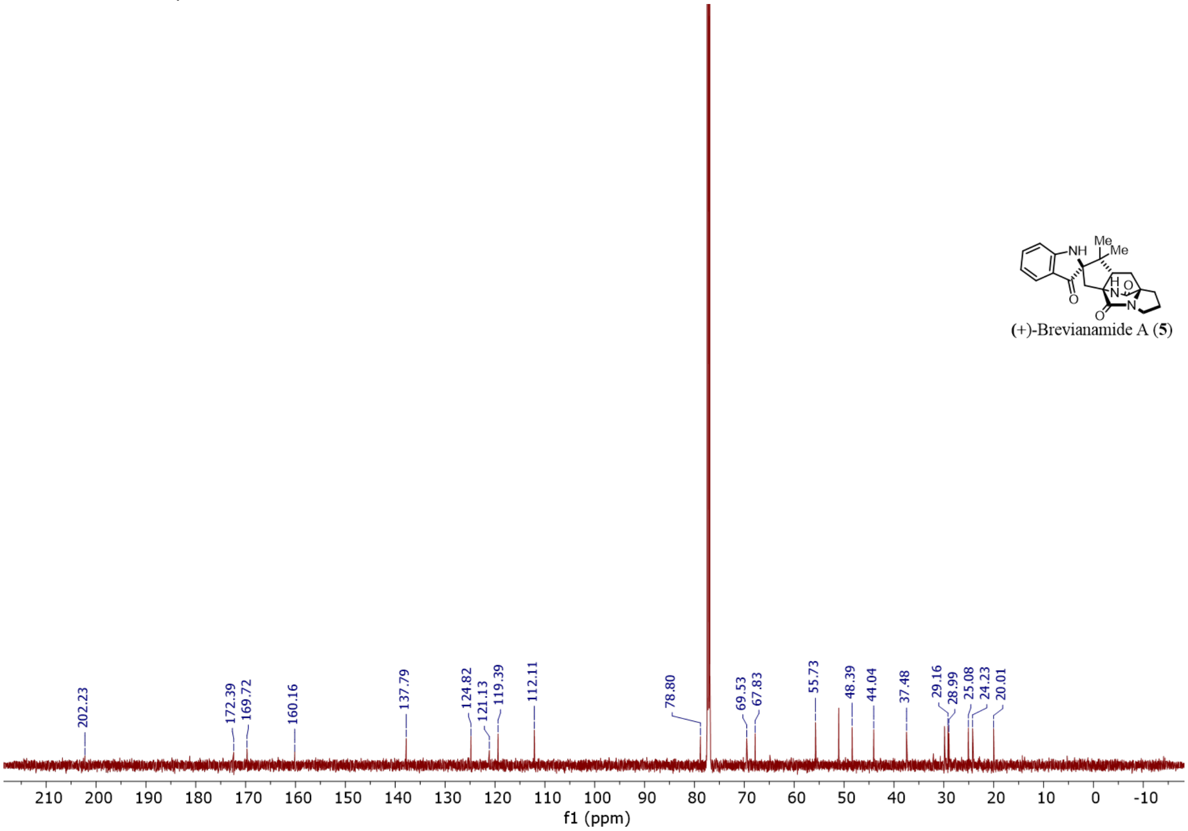

**Supplementary Figure 59.** $^{1}$H NMR spectrum of (+)-brevianamide B (**6**) from LiOH reaction (599 MHz, $CDCl_3$)

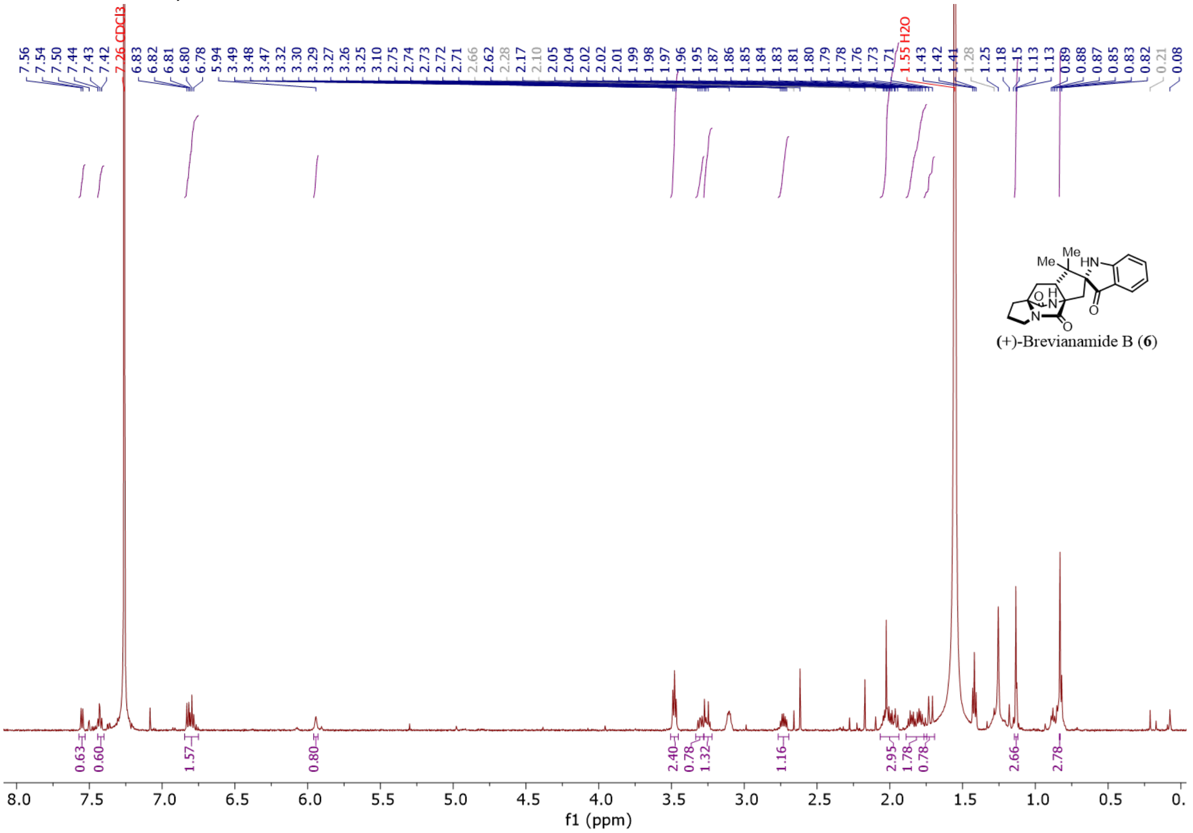
